# Punctuated memory change: The temporal dynamics and brain basis of memory stability in aging

**DOI:** 10.64898/2026.01.20.700557

**Authors:** Anders M. Fjell, Edvard O.S. Grødem, Didac Vidal-Piñeiro, Ole Rogeberg, Øystein Sørensen, Pablo F Garrido, Leiv Otto Watne, Anders Lundquist, the Alzheimer’s Disease Neuroimaging Initiative, the Vietnam Era Twin Study of Aging, Mayo Clinic Study of Aging, the Australian Imaging Biomarkers and Lifestyle flagship study of ageing, The Health and Aging Brain Study (HABS-HD) Study Team, Lars Nyberg, Kristine B. Walhovd

## Abstract

Are there individuals who resist episodic memory decline into older age? Analyzing 728,000 memory tests from 80,000 participants with at least 4 assessments, we introduce a simulation-calibrated framework to identify genuine memory stability. Across cohorts and models, ∼10% of adults ≥70 years showed stable performance over a decade. In an MRI subgroup (n≈2,000), stable performers exhibited lower rates of brain atrophy across widespread regions, anchoring cognitive stability in structural brain maintenance. However, stability was often transient rather than trait-like: many individuals followed trajectories with extended plateaus of stable performance punctuated by episodes of accelerated decline. Accordingly, 54% showed at least one period of observed stability, averaging 10 years, whereas only 0.4% upheld stable performance over 24 years under the strictest definition. These findings are consistent with a complex-systems model of cognitive aging in which decline often reflects critical transitions rather than continuous erosion.

## Introduction

How many, if any, individuals show stable episodic memory function as they age, and what does such stability imply about the fundamental shape of cognitive aging? Despite the well-documented average decline in memory after age 60 (1, 2), some individuals appear resistant to this trend (3, 4). This phenomenon has motivated the influential concept of “brain and memory maintenance” used to describe older adults who show little decline (5), yet estimates vary widely across studies (6–9). In one influential report, ∼18% of adults aged 35–85 in the Betula cohort were classified as memory maintainers over 15 years (10). An unresolved question is whether this broader class of low-decline trajectories includes a distinct subgroup with enduring near-zero change over long intervals, and whether apparent stability can reflect time-limited plateaus embedded within broader decline. Prevailing models typically treat cognitive aging as smooth and monotonic, with individual differences expressed mainly as variation in level, onset, and rate of decline (11–13); under this view, ‘memory maintenance’ largely reflects the shallow-slope end of a continuous distribution, and sustained stability should be relatively rare and trait-like. However, low average change can in principle arise from different temporal architectures. Here we propose a complementary, mechanistic interpretation of stability: that apparently stable memory trajectories in aging are often organized as periods of relative stability punctuated by briefer episodes of accelerated loss (Figure 1). Under this view, stability can reflect a transient state of resilience rather than a fixed lifelong trait, consistent with a complex-systems framework (14) in which systems shift between stable basins of attraction via critical transitions. We do not assume that punctuated dynamics replace smooth decline for all individuals; rather, we test whether they are sufficiently common to shape observed stability and population-average trajectories.

**Figure 1.**
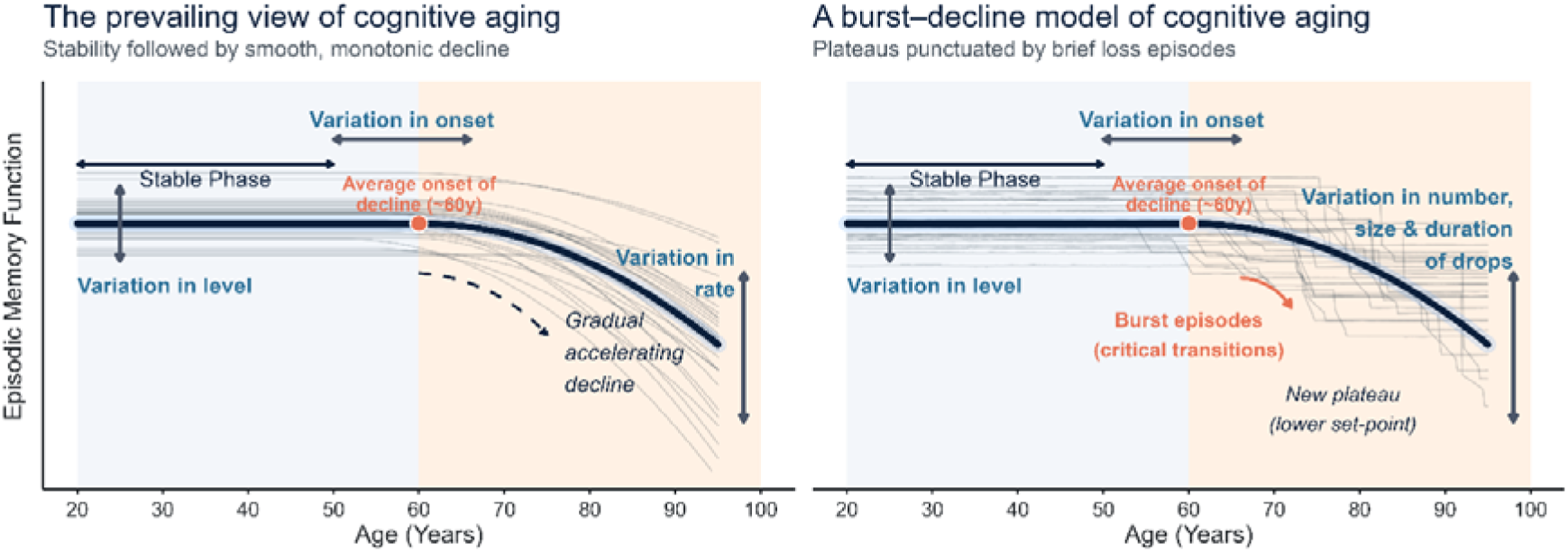
Conceptual models of cognitive aging: smooth decline versus burst–decline dynamics. Left: Prevailing models depict episodic memory as relatively stable through early adulthood, followed by a smooth, monotonic—and often accelerating—decline at older ages. Individual trajectories differ in level, age of onset, and rate of decline, but change is assumed to be gradual over time. Right: The burst–decline model proposed here posits that individual trajectories are often organized as extended plateaus of relative stability punctuated by brief episodes of accelerated loss (“bursts”), followed by stabilization at a new, typically lower, performance level. In this framework, stability can be a dynamic and potentially transient state rather than a fixed lifelong trait. The two panels share the same group-average trajectory, illustrating that similar population-level age curves can arise from fundamentally different underlying within-person dynamics.

Demonstrating genuine stability, however, is a profound methodological challenge that has obscured its theoretical interpretation. High late-life performance may simply reflect lifelong advantage rather than a lack of change (15), and estimates of stability are confounded by practice effects and measurement noise. Furthermore, because age-related change is modest relative to this noise, apparent stability often arises from chance rather than resilience. Study participants are also often healthier and have higher cognitive function than the target population, biasing naïve estimates. To resolve this, we introduce a rigorous, practice-adjusted framework for population-representative cohorts, calibrated against null simulations to strictly distinguish true signal from statistical noise. This allows us to move beyond simple prevalence estimates and use stability as a tool to interrogate the underlying dynamics of cognitive aging.

We use longitudinal episodic memory data from three premier population-based longitudinal cohorts - the Survey of Health, Ageing and Retirement in Europe (SHARE) (16), the Betula Prospective Cohort Study: Memory, Health and Aging (Betula) (17, 18), and the US Health and Retirement Study (HRS) (19) - to provide a new view on memory stability in aging. We go beyond quantifying prevalence to interrogate the temporal structure of stability itself. By combining multi-decadal observation windows with historical back-projections and neuroimaging, we test whether stable trajectories observed in old age are plausible continuations of midlife function or whether they can represent transient plateaus following unobserved decline. We anchor these cognitive patterns in the brain and health by assessing rates of brain atrophy and mortality risk, correcting for selection bias to ensure results are not driven by “healthy survivor” effects. Together, these analyses test whether stability is an enduring phenotype or a time-limited state, and whether apparently smooth population-level decline can arise from punctuated individual-level change.

## Results

We structured the analyses to move from identifying stability to understanding its implications for models of cognitive aging. First, we establish the existence and prevalence of statistically robust periods of memory stability in large population-based data. We validate these findings across independent cohorts and analytical frameworks. Second, we anchor stability in neurobiology by testing its association with longitudinal brain atrophy. Finally, we examine whether stability reflects a stable trait or a transitory state by analyzing within-person dynamics and back-projected trajectories. We show how these empirical findings motivate a burst–decline model of cognitive aging and demonstrate that smooth population-level decline can arise from heterogeneous, punctuated individual trajectories.

Sample details are presented in Online Methods (section *Samples*, Online Methods Table 1). A total of 211,431 participants with 72,066 memory test sessions were analyzed, including 80,679 participants with ≥ 4 sessions.

### Results section 1 – Identification of stable performers

Initial analyses were performed in SHARE, using the sum of the immediate and delayed verbal recall tests (max 20). An ordinary least squares model with a flexible age adjustment and a monotonic, diminishing-returns practice term, was fit to the full sample (n=155,606) to control for the number of prior tests, and the residuals were used in the analyses (Online Methods Figure 5). Further analyses were restricted to participants with ≥ 4 tests (n=49,602, follow-up = 11,1 years, range 6-18, restricted to baseline age 50-90 years). We estimated individual memory change using Theil–Sen regression on each participant’s practice-corrected recall scores (Online Methods Figure 6), yielding a robust change estimate less sensitive to measurement noise. Across age, longitudinal decline (-0.197 words/year, 95% CI: -0.192, -0.200) was consistently steeper than cross-sectional estimates (Online Methods Figure 7, SI Section 1), an expected discrepancy (20) reflecting selective survival of higher-functioning individuals, cohort effects, and the mathematical distinction between interval-averaged slopes versus instantaneous rates in the context of accelerating decline.

To classify participants as “stable performers,” we used the Theil–Sen slopes and quantified uncertainty via leave-one-out jackknifing (Online Methods Figure 6 and Table 4). A participant was labeled a stable performer if their slope and its lower confidence bound both exceeded a specified threshold (≥ 0), ensuring that apparent stability was not driven by outliers (see SI). Thus, 0 controls how much decline is allowed, and the lower bound (LB) criterion controls how certain we must be that the estimated slope is not more negative than 0.

To distinguish true stability from measurement noise, we calibrated these metrics against null simulations where participants followed the population-average decline with added individual noise extracted from the real data (Online Methods, section *Simulation experiment*). We tested a grid of 15 definitions varying in strictness (Online Methods Figure 8). Crucially, the observed prevalence exceeded the null simulation inter-quartile range for all 15 rules, demonstrating a robust signal beyond chance (SI Section 2). We observed a fundamental trade-off: stricter criteria (less negative 0, higher LB) yielded higher Positive Predictive Values (PPV), ensuring high confidence in individual classification, but reduced sensitivity. Conversely, more lenient criteria maximized the separation between observed and null curves, better capturing the magnitude of the stability phenomenon in the population despite lower specificity. We therefore focus primarily on a conservative rule to optimize individual-level certainty (slope > 0 & LB95%), while also reporting prevalence under a more lenient rule that better recovers population signal (slope > −0.1 & LB50%; Fig. 2 and SI Section 4 and 14).

**Figure 2.**
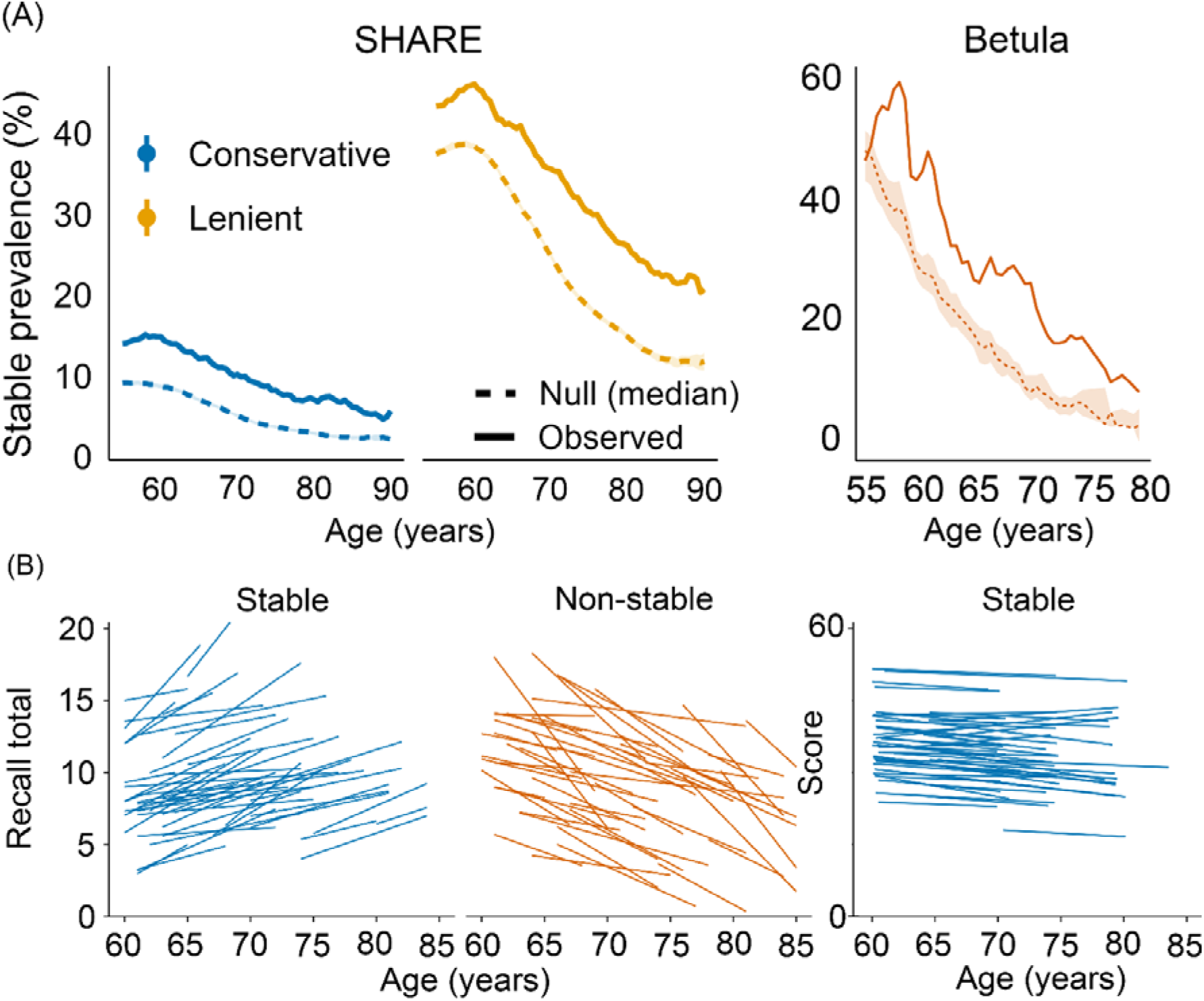
Prevalence of participants with stable memory function in SHARE and Betula. A: Observed prevalence of stable memory performers (solid line) vs. null expectation (dashed line) (median, shaded band = null IQR), separately for the conservative and the lenient stable performer criteria in SHARE (two left plots) and Betula (right plot). B. Lines show Theil–Sen slopes for 50 randomly selected stable performers (O > 0, LB95%, blue) and non-stable performers (O < -0.1 or LB50%, orange) in SHARE (two left plots) and Betula (right plot).

Stable performers were identified across the age spectrum, though their proportion declined approximately linearly (Figure 2). To estimate the true population prevalence, we corrected observed counts for false positives and negatives using simulation-derived PPV. Assuming 80% sensitivity, true stable performers comprised 5.3% (conservative) to 17.1% (lenient) of the population aged ≥ 70. Based on the midpoint of these bracketed estimates, we conclude that ≈10% of adults over 70 have stable episodic memory over a decade. Sensitivity analyses using different sensitivities (0.70–0.90) indicated these estimates are robust, varying by only 2-3 percentage points.

### Validation using hierarchical Bayesian modeling

To evaluate robustness under an independent statistical framework, we fit a hierarchical Bayesian model on the full dataset not restricted to participants with ≥ 4 observations, jointly estimating the population trajectory and individual deviations while explicitly accounting for uncertainty, practice effects, and non-Gaussian score distributions (Online Methods Figure 10). Bayesian estimates closely matched the robust Theil–Sen results, with prevalence of stable performers declining linearly from 60 to 90 under both slope criteria (SI Section 3). Bayesian and frequentist prevalence estimates were nearly identical. For slope ≥ 0, posterior means for >70 years was 5.2% (95% CI 4.28–6.03) compared to 5.3% according to the Theil-Sen estimates, and 15.9% (95% CI 13.6–17.5) vs 17.1% for slope ≥ -0.1. This alleviates concerns that Theil–Sen might be affected by finite-sample properties, the partly ad-hoc nature of the LB rules, and non-Gaussian errors affecting the results.

### Independent sample extension – analyses of the Betula cohort with extended follow-up

To assess whether stable performers can be identified in a sample starting at younger ages and followed for longer periods, we repeated the analyses in the Betula cohort (n = 4423/ 9731 observations, using participants with 4–5 memory assessments covering 14–25 years (Figure 2, n = 884/ 4344 observations, average follow up 19.5 years). The practice-corrected slope distribution was shifted to the left compared to SHARE (Online Methods Figure 7), likely reflecting less measurement noise due to longer follow-up time and use of a composite score based on 5 tasks (max 76 points). Although few participants qualified as stable performers under the strict conservative criterion (slope ≥ 0, LB95%), the observed stability signal still exceeded the null-distribution, yielding a PPV-adjusted prevalence for age ≥ 70 of ≈1.5%. When applying the lenient threshold (scale-adjusted slope ≥ −0.4 points per year [≈1% annual loss], LB50%), the PPV-adjusted true prevalence above age 70 was ≈12.5% (SI Figure 21), compared to 17.1% for SHARE under the equivalent rule, likely reflecting the longer follow-up intervals in Betula. These results were also replicated using hierarchical Bayesian modeling (SI Section 3 and 14). Collectively, these analyses confirm that stability can be identified over periods as long as 20+ years, though it is less prevalent than over 10-year windows and is rare under the most conservative definition.

### Selection and survival bias: Is stability a survival trait?

We used mortality as an external criterion to assess representativeness, selection or survival bias between stable performers (conservative rule, n = 5,261, 419 deaths) and non-stable performers (n = 42,924 non-stable performers, 4,434 deaths) (SI Section 8). In an initial sex-adjusted Cox model stratifying baseline hazards by country, we found a dissociation between memory performance and the stable performer label (See caption Figure 4). As expected, higher memory level and a shallower slope were robustly associated with lower mortality (Level HR = 0.79; Slope HR = 0.90; p < 0.001). However, the binary stable performer label itself was associated with a 17% higher hazard of death in the primary model (HR = 1.17, p = 0.003). This increased risk was not explained by differential dropout; weighting the models by the inverse probability of censoring (IPCW) yielded virtually identical results (HR = 1.17), indicating the finding is not a byproduct of attrition bias (see SI for full sensitivity analyses). Crucially, when the binary stable performer label was added to a model already containing the robustly protective continuous level and slope features, the label conferred an even stronger independent risk (HR = 1.51, p = 2 × 10^-12^, Figure 4). These findings caution against interpreting the stable performer label as a marker of a super-healthy elite. Instead, they suggest heterogeneity within the group and are consistent with stability sometimes reflecting a transient plateau that can mask underlying vulnerability. This supports the hypothesis that stability can represent a temporary plateau rather than lifelong resistance. To test this “transient state” hypothesis, while also investigating the rare subpopulation of long-term stable performers (20+ years) identified in Betula, we turned to the US Health and Retirement Study (HRS; n = 38,448/229,234 observations), taking advantage of its follow-up examinations extending to 25 years (n = 24,969/ 200,696 observations with ≥ 4 follow-ups over _x_ℒ =14.7 years, range 4-25 years).

### Stability as a Time-Limited State: Findings from the HRS

We shifted the analytical unit from the *person* to the *stability period*, allowing us to test whether we could capture stability as a phase within a longer trajectory. Accordingly, ‘stability’ is defined here as the occurrence of at least one qualifying stable bout, rather than as a property of the entire follow-up. Using this dynamic approach, we found that temporary stability is remarkably common. In HRS, 53.8% of participants exhibited at least one observed ≥4-wave (∼6 years) contiguous period - *bout* - meeting the conservative stable performer criterion (0 ≥ 0, LB95%), with the longest such period lasting on average 10.0 years across 6-7 waves. Pre-bout slopes could be estimated for 5,461 participants (40.5% of bout-positive cases had at least two timepoints preceding the stable period), and among these 57.7% showed decline prior to the stable period (mean pre-decline slope = −0.49 words/year). Post-bout slopes could be estimated for 5,580 participants (41.4% of bout-positive cases), and among these 63.6% showed decline after the stable period (mean post-decline slope = −0.50 words/year). Consequently, the prevalence of stable performer status dropped sharply with increasing follow-up duration. The result of this is clearly seen in Figure 3 (top panel), showing that participants > 70 years with a stability bout of up to 9 waves of no decline still showed a negative slope when calculated across their whole follow-up period.

**Figure 3.**
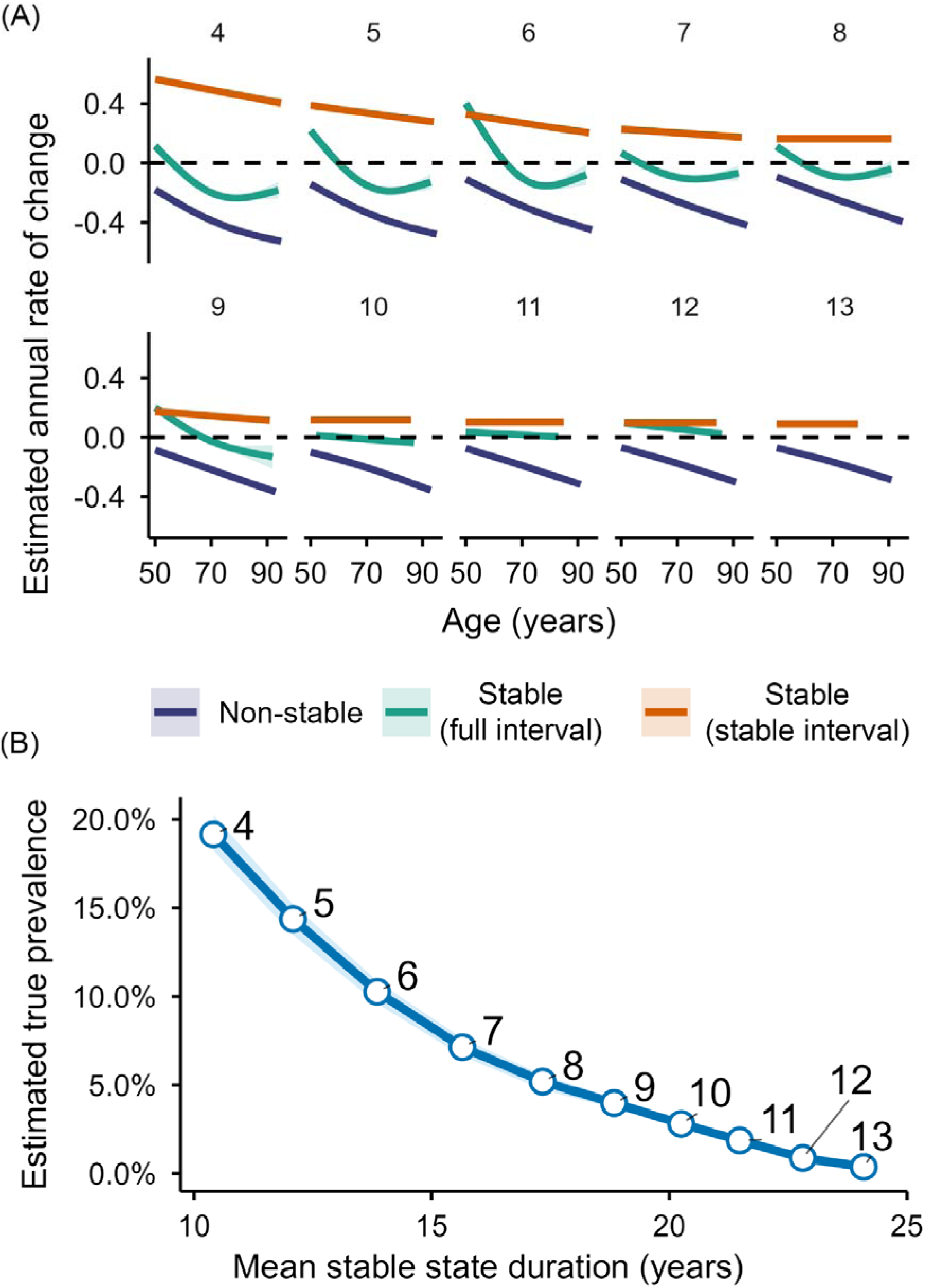
Stability-period dynamics and prevalence of memory stability in HRS. A: Age variation in memory-change rates during stability periods. Each panel shows GAM smooths of Theil–Sen annual change in practice-corrected episodic memory as a function of age for a specific stability-duration threshold, i.e. all participants with stable performance over ≥ 4 waves, ≥ 5 waves, etc. up to 13 waves. For stable performers, slopes are shown both for the identified stable window and for the full individual observation window; for comparison, non-stable performers show the overall slope across the full observation window among participants with at least the corresponding number of waves but without a stable period of that length. Shaded bands indicate 95% CI, the dashed line marks zero change. B: True prevalence of memory stability in ages 75–85 across bout durations. Points and line show the estimated true prevalence of having at least one contiguous stable period of length ≥W waves (labels), plotted against the mean duration (years) among those meeting each threshold. The shaded band reflects uncertainty from the null simulations. Prevalence estimates are corrected for false positives and false negatives assuming sensitivity Se = 0.80, using the simulated null prevalence as the false-positive rate.

Notably, the null model produced a substantial “stability” base rate, especially for shorter bout thresholds, underscoring that observed prevalences cannot be treated as ground truth and motivating PPV-based correction for false positives and false negatives. We used the same simulation framework as for SHARE to obtain PPV values to estimate true stable performer prevalence in the age-range 75–85 years across follow-up periods of different durations. Crucially, the observed proportion of stable performers was still significantly above chance even at the maximum duration (13 follow-ups, ≈25 years). Using a misclassification-corrected prevalence estimator (sensitivity=0.80) that subtracts the null ‘chance stability’ rate and adjusts for false negatives, the corrected prevalence in ages 75–85 was 2.80% for ≥10 waves (mean duration 20.3 years), 1.86% for ≥11 waves (21.5 years), 0.87% for ≥12 waves (22.8 years), and 0.37% for 13 waves (24.1 years) (Figure 3). Across the full HRS follow-up (any age), the corrected true prevalence of having at least one stable period was 14.7% for ≥4 waves, declining to 1.58% for ≥10 waves and 0.17% for ≥13 waves. Prevalence is higher in HRS than in SHARE in part because the HRS analyses quantify stability as the presence of at least one qualifying stable window that can occur anywhere within follow-up, whereas SHARE analyses treated stability as a person-level property across the available series, which is a stricter criterion. These results confirm that while transient stability is a widespread feature of cognitive aging, often reflecting a “post-burst” plateau, genuine multi-decadal stability exists as a rare, robust phenomenon distinct from the noise of gradual decline. For comparability, we also applied a window-based approach in SHARE; as expected, prevalence declined steeply with increasing duration threshold, with somewhat lower values reflecting shorter time coverage (SI Section 11). In all cases, observed prevalence exceeded the simulated null prevalence, indicating that the procedure detects a stability signal even at the longest durations.

Finally, we compared the prevalence of qualifying stable windows when restricted to the first waves versus the last waves. Across thresholds, prevalence in the first and last windows was remarkably similar (e.g., stability over the first 4 waves vs. the last 4 waves: 12.9% vs 14.5%; stability over the first 10 waves vs. the last 10 waves: 4.2% vs 4.9%), indicating that stability is not preferentially concentrated at either end of follow-up and is consistent with decline occurring both before and after stable periods (SI Section 11). In contrast, allowing stable windows to occur anywhere yielded much higher prevalence (W=4: 48.5%). Together, these results support stability as a time-limited state that can occur at different points within an individual’s observation window rather than reflecting a baseline trait or an end-stage plateau. Stability is therefore not a single phenotype but reflects at least two resilience pathways - post-burst stabilization and robust long-term stability, highlighting the role of timing of critical transitions in cognitive aging.

## Results section 2 – Biological and clinical validation of stable performers

### Brain basis of stable memory performers

To anchor the cognitive classification in the brain, we analyzed longitudinal MRI data from 10 independent studies (n =1,978 participants aged ≥ 50 with ≥ 4 episodic-memory test sessions covering 16,342 tests and 7,206 scans, see Online Methods Table 2). We extracted a principal component across memory subtests within each study and classified stable performers using the same Theil–Sen–based framework. The PPV-estimated proportion of stable performers was 9.8% (95% CI: 9.3–10.4) for ages ≥70 and 8.7% (95% CI: 8.2–9.2) for ages ≥80, slightly higher than in SHARE and BETULA, likely reflecting more selection and survival bias in MRI cohorts (SI Section 5). We used the same Theil–Sen framework to assess volumetric brain change and fitted interval-weighted logistic regressions comparing conservative stable performers (n = 215; mean age 68.5 years, mean memory/ MRI follow-up 9.3 and 6.1 years, respectively) to lenient non-stable performers (n = 1,387; mean age 72.4 years, mean memory/ MRI follow-up 8.1 and 5.0 years, respectively), adjusting for age, sex, and intracranial volume. Stable performers exhibited significantly less atrophy and ventricular expansion than non-stable performers in 27 of 44 regions (FDR < 0.05) (Figure 4, SI Section 6).

**Figure 4.**
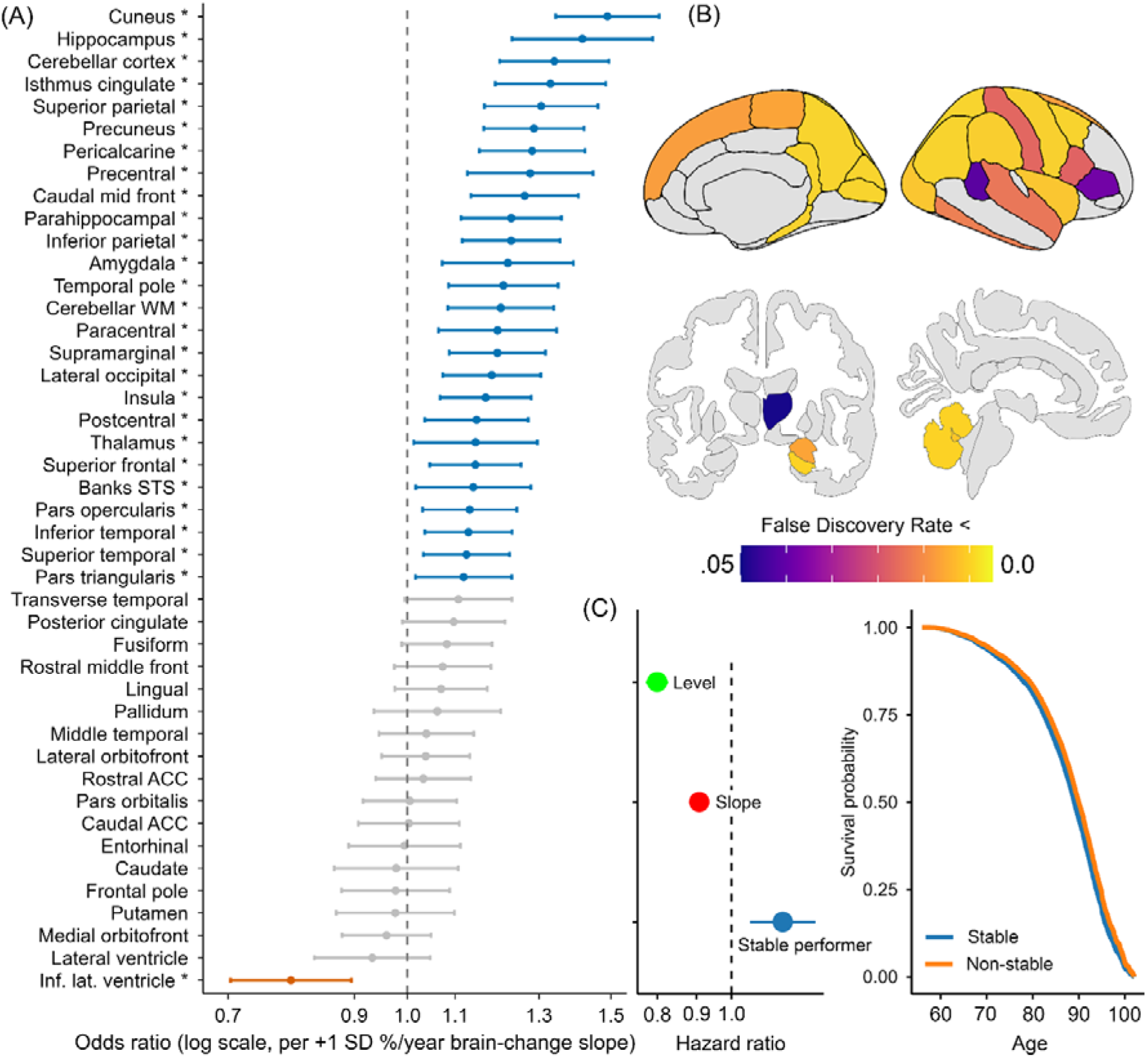
Associations between memory stability, brain atrophy and mortality. A: Forest plot showing odds ratios for being a stable performer vs. non-stable performer per one SD more positive brain-change (Δvolume/year) in bilateral regions. Models were adjusted for age at MRI midpoint, sex, and estimated total intracranial volume, and were weighted by individual brain follow-up span (longer trajectories contributed more). Error bars show 95% CI, and asterisks indicate FDR < 5% (Benjamini–Hochberg). ACC: Anterior Cingulate Cortex. WM: White Matter. STS: Superior Temporal Sulcus. Inf. Lat.: Inferior Lateral. B: Anatomical location of brain change relationships. Regions with FDR ≤ 0.05 projected onto an inflated template for cortical (top) or subcortical (bottom) volume using ggseg software(21). Analyses were done on bilateral volume and shown on the right hemisphere for illustrative purposes. Inferior lateral ventricles are not shown. C: Mortality analyses using chronological age Cox regression anchored at each participant’s classification age (the observed visit closest to the person’s median testing age). Left panel shows hazard ratios (HRs, log scale) with 95% CIs for three predictors: memory level (per SD at the classification age), memory slope (per SD less decline), and the binary stable performer label. Level and slope estimates come from a sex-adjusted and the stable performer estimate from an IPCW Cox model, all models were stratified by country. Right panel displays IPCW-weighted survival curves for stable vs non-stable performers predicted for a typical profile (sex and country set to their most frequent categories in the sample).

Slower decline was particularly evident in the medial temporal lobe (bilateral hippocampus and parahippocampal cortex) and widespread frontoparietal cortices. Quantitatively, 1 SD less volume loss was associated with up to 50% higher odds of being a stable performer. Repeating this analysis using robust volume levels yielded no significant associations, reinforcing that memory stability is specifically linked to the preservation of brain structure over time, not brain size per se (details in SI).

### Health and psychosocial characteristics

We used IPCW univariate age-adjusted logistic models to relate a broad set of life-course, health, and psychosocial variables to stable performer status (conservative stable performers vs. lenient non-stable performers). Stable performers were generally healthier (e.g., lower rates of smoking and depressive symptoms) and had better orientation scores (Figure 5), but effect sizes were universally small (0.9 < OR < 1.1, see SI Section 7 for a series of sensitivity analyses confirming the results). Testing all variables simultaneously by multivariate elastic-net modeling achieved only modest discrimination (AUC = 0.614), offering negligible improvement over a model based on age alone (AUC = 0.608). This indicates that while stable performers are on average slightly healthier, they do not constitute a distinct sociodemographic or healthier class, and cannot be robustly predicted by standard risk factors.

**Figure 5.**
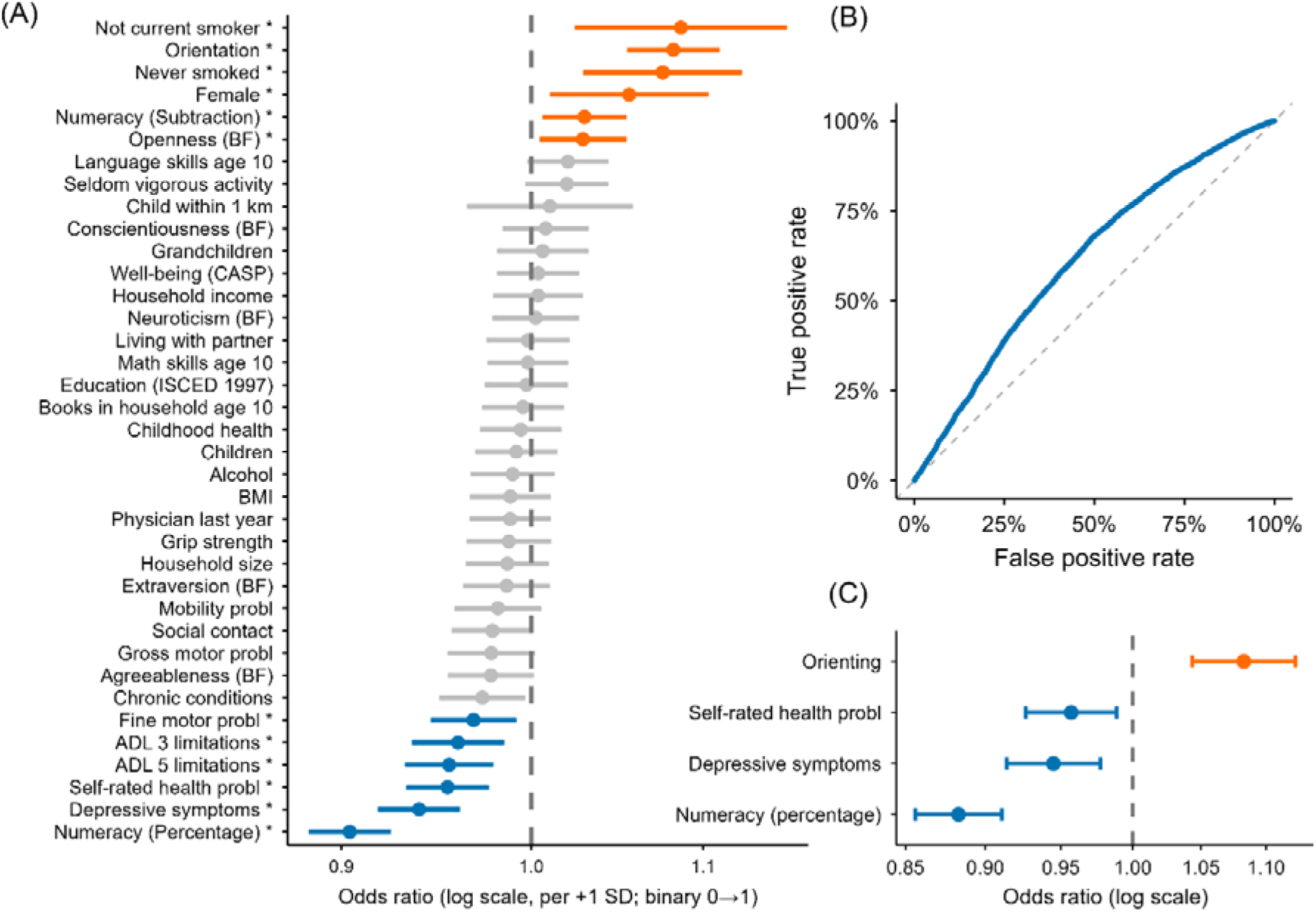
Classification of stable performers. A: Forest plot of all variables in the univariate analysis. Points are odds ratios (OR) and horizontal bars are 95% CIs (per +1 SD for continuous measures), and asterisks indicate FDR < 5% (Benjamini–Hochberg). Bottom: Multivariate classification of stable performers. B: Receiver–operating characteristic (ROC) curve from an elastic-net logistic model predicting stable performer status from all candidate predictors. The diagonal dashed line marks chance. C: Forest plot of the variables retained in the multivariate model after refitting with maximum likelihood for inference. Points are odds ratios (OR) and horizontal bars are 95% CIs (per +1 SD for continuous measures). Age was included in all models but is not displayed.

## Results section 3 - Distinguishing lifelong slopes from episodic decline dynamics

### Level–slope dynamics and the back-projection paradox

Performance level was a weak predictor of stable performer status; a +1 SD higher robust intercept increased the odds of being a stable performer by only 11% (OR = 1.11, 95% CI: 1.08–1.15, SI Section 10), confirming that classification captures trajectory stability rather than lifelong high function. Plotting the performance of stable performers showed that many have stable scores at low levels (Figure 6). This raises a critical question: have these individuals always performed low, did they decline earlier and then stabilize, or can cohort differences account for this pattern?

**Figure 6.**
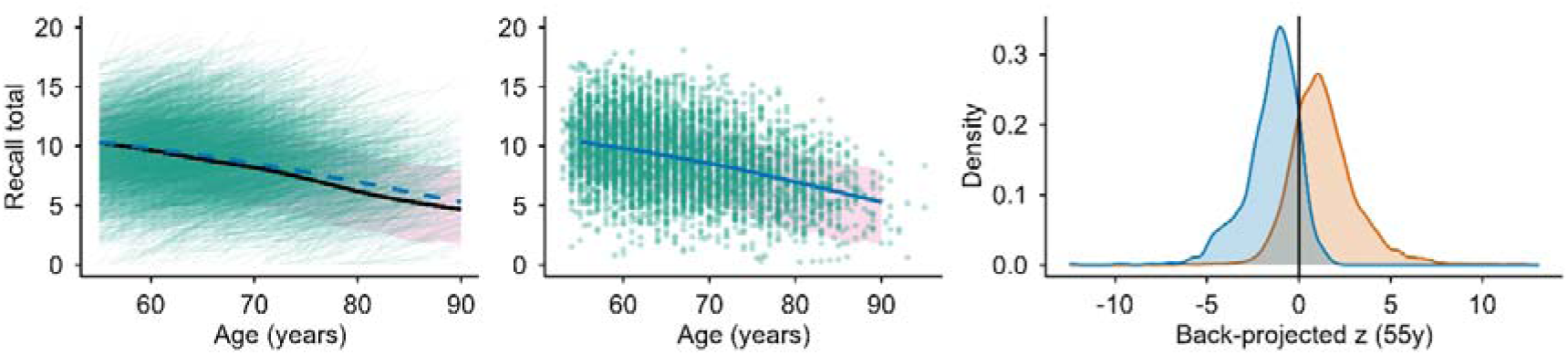
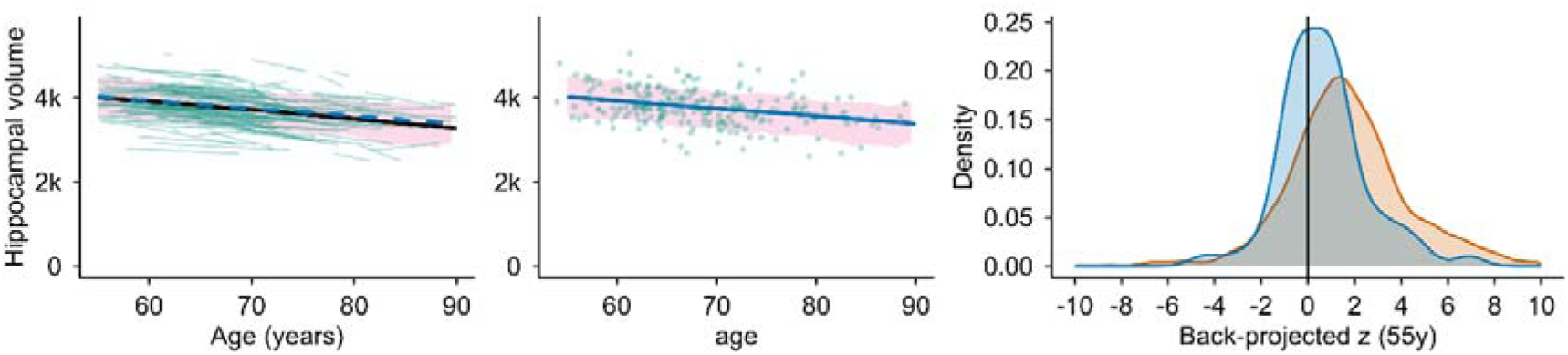
TS-estimated memory and hippocampal volume level in all SHARE participants and stable performers. Each participant’s memory and hippocampal volume level was defined as the Theil–Sen–estimated score at their mean age. Top left: Practice-corrected mean memory level as a function of age for all participants with ≥4 observations (black line) and for stable performers according to the conservative rule (blue dashed line). The pink band shows the central 75% of levels in the full sample, and the thin lines show the trajectory for each stable performer. Middle: Memory level for each stable performer, the age-trajectory for the stable performers and the central 75% in the full sample (pink). Right: Distribution of back-projected performance at age 55 (z-score within birth cohort) among participants aged ≥75 at any measurement, for stable performers (blue) and non-stable performers (orange). The vertical line marks the cohort mean (z = 0). Bottom panels: The same analyses as above for hippocampal volume averaged across hemispheres.

To distinguish between these possibilities, we back-projected the estimated trajectories of older SHARE participants (≥75 years) to age 55, adjusting for birth cohort (stable performers n=970; non-stable n=9,531). The results were incompatible with an enduring, near-flat trajectory for the stable performers: if their current level and slope were extrapolated backwards, 79.8% of stable performers would fall below the 25th percentile of their cohort at age 55, and they were 11.6 times more likely than non-stable performers to fall below the 5th percentile (Figure 6). These implausibly low reconstructed scores strongly suggest that for many, the observed stability was preceded by unobserved, substantial loss periods (“bursts”).

We confirmed this dynamic in the HRS, where longer follow-up allowed us to define stability as a transient state (a qualifying ≥4-wave bout) rather than as a stable trait, permitting stability periods to characterize only part of each individual’s observation window. The back-projection paradox persisted: when we back-projected memory to age 55 using Theil–Sen-estimated linear models anchored to the stable bout itself, 73.5% of bout stable performers (2,489/3,387) fell below the 25th percentile of their birth cohort at age 55, compared with 6.7% of non-stable performers (233/3,500) (SI Section 10). Bout stable performers were also ∼40× more likely than non-stable performers to fall below the 5th percentile (27.5% vs 0.69%; risk ratio [RR] = 40.2). Crucially, this paradox was strongly attenuated when bout stable performers were instead back-projected using their slope across the full observation window (10.5% below the 25th percentile and 1.6% below the 5th percentile, compared with 6.7% and 0.7% among non-stable performers; RR <5th = 2.3, Figure 7), indicating that the stable slope is locally valid for a time-limited plateau but not representative of long-run change. Finally, using a purely window-based definition, participants classified as stable in the last four waves did not show reduced decline earlier: among those with ≥10 observations, last-window stability was not associated with the first-window rate of change (β = −0.013 words/year, p = 0.45). Together with the within-person state analyses showing that stability periods commonly occur in the context of broader decline, these results indicate that much of the observed late-life stability in HRS reflects a plateau embedded within a non-monotonic change rate trajectory rather than a lifelong state of resistance.

**Figure 7.**
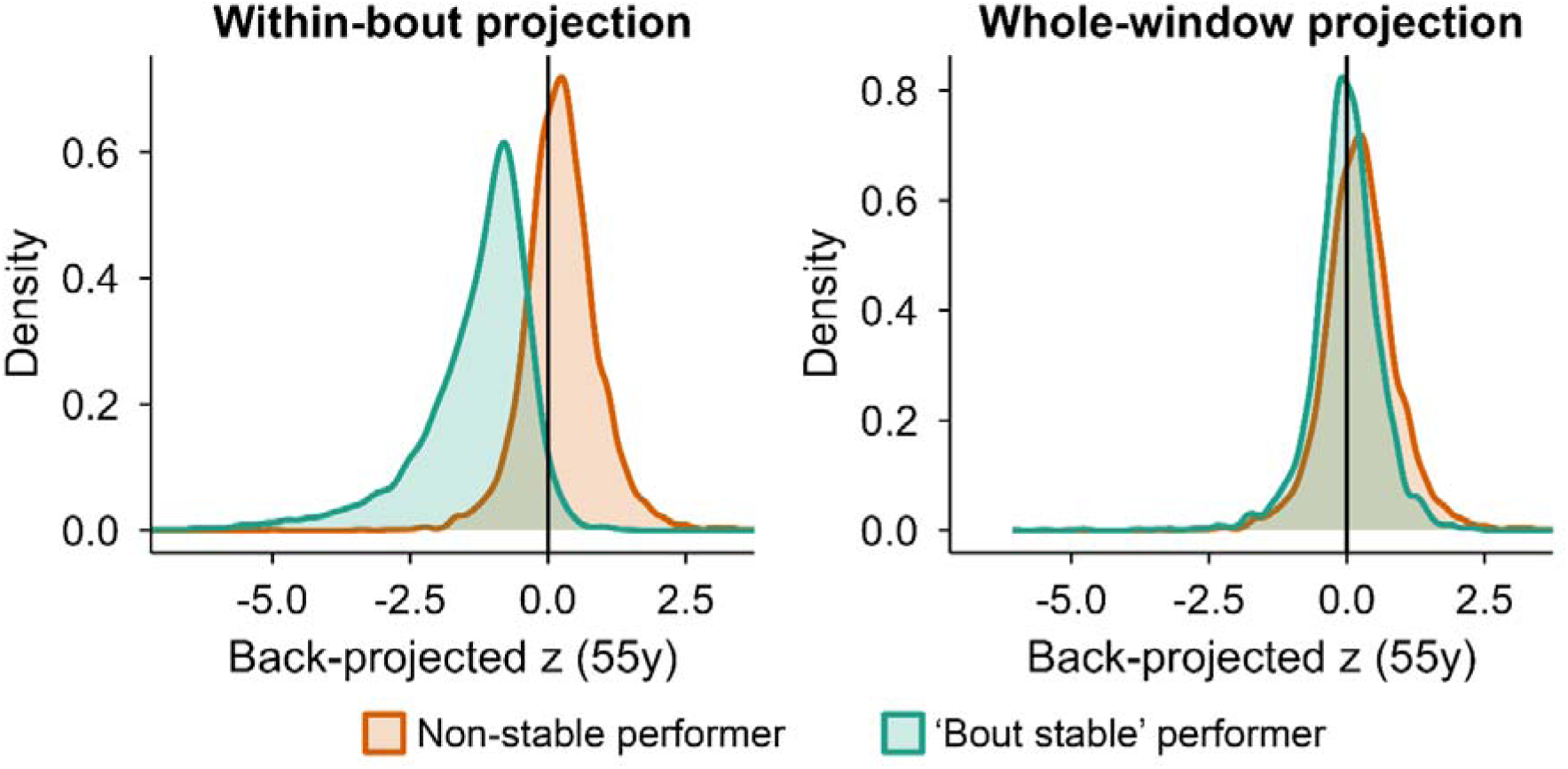
Back-projection paradox supports stability as a time-limited state in HRS. Density plots show the distribution of birth-cohort–standardized back-projected memory performance at age 55 (z-scores) among older HRS participants (defined as having maximum observed age ≥75), contrasting bout stable performers with non-stable performers (orange). The vertical line marks the birth-cohort mean (z = 0). Left (Within-bout projection): For bout stable performers, the back-projection is anchored to the identified ≥4-wave stable bout (Theil–Sen slope and intercept estimated within the bout), producing a strong “paradox” in which stable performers back-project to implausibly low midlife performance. Right (Whole-window projection): Repeating the same back-projection but estimating the stable slope over the entire observation window substantially attenuates the paradox, indicating that the near-zero slope is locally valid for a plateau state rather than representing long-run resistance to decline. Non-stable performers are projected using their overall Theil–Sen line across all observations in both panels; cohort standardization uses midlife norms derived from observations in ages 50–60 (see Online Methods).

We also explored brain structural correlates in the MRI subsample, finding only a slight skew towards smaller hippocampal volumes in stable performers at age 55 (Figure 6 and SI), although this secondary analysis had limited precision given the small number of stable performers (n = 64) relative to non-stable performers (n = 873).

Finally, we evaluated offset–slope dynamics in Betula, where participants were followed from midlife over ≥20 years, comparing the GAM-derived age-trajectories of Theil–Sen-estimated intercepts for stable performers and non-stable performers. There was no main effect (p=.77) or age × stable performer interaction (p=.32), supporting the finding that stability is weakly related to current performance level across the age range (SI Section 10).

Together, the three datasets reveal heterogeneity within stable performance, where many late-life stable trajectories likely reflect post-burst stabilization, whereas a smaller subgroup shows genuine long-term stability over 20+ years. This pattern implies that stability can be locally real but globally transitory, and motivates testing whether smooth population-average decline can arise from punctuated within-person change.

### Identification of a burst-decline mechanism

We simulated longitudinal data for 5,000 participants following stable baselines interrupted by 1–3 age-dependent “bursts” of accelerated loss (Online Methods, section *Individual vs. group trajectory simulation*). Despite the punctuated nature of all individual change in this simulation, the aggregated group curve reproduced the smooth, accelerating decline observed empirically (Figure 8), demonstrating that smooth population-level memory decline can arise even when individual change occurs in sporadic bursts.

**Figure 8.**
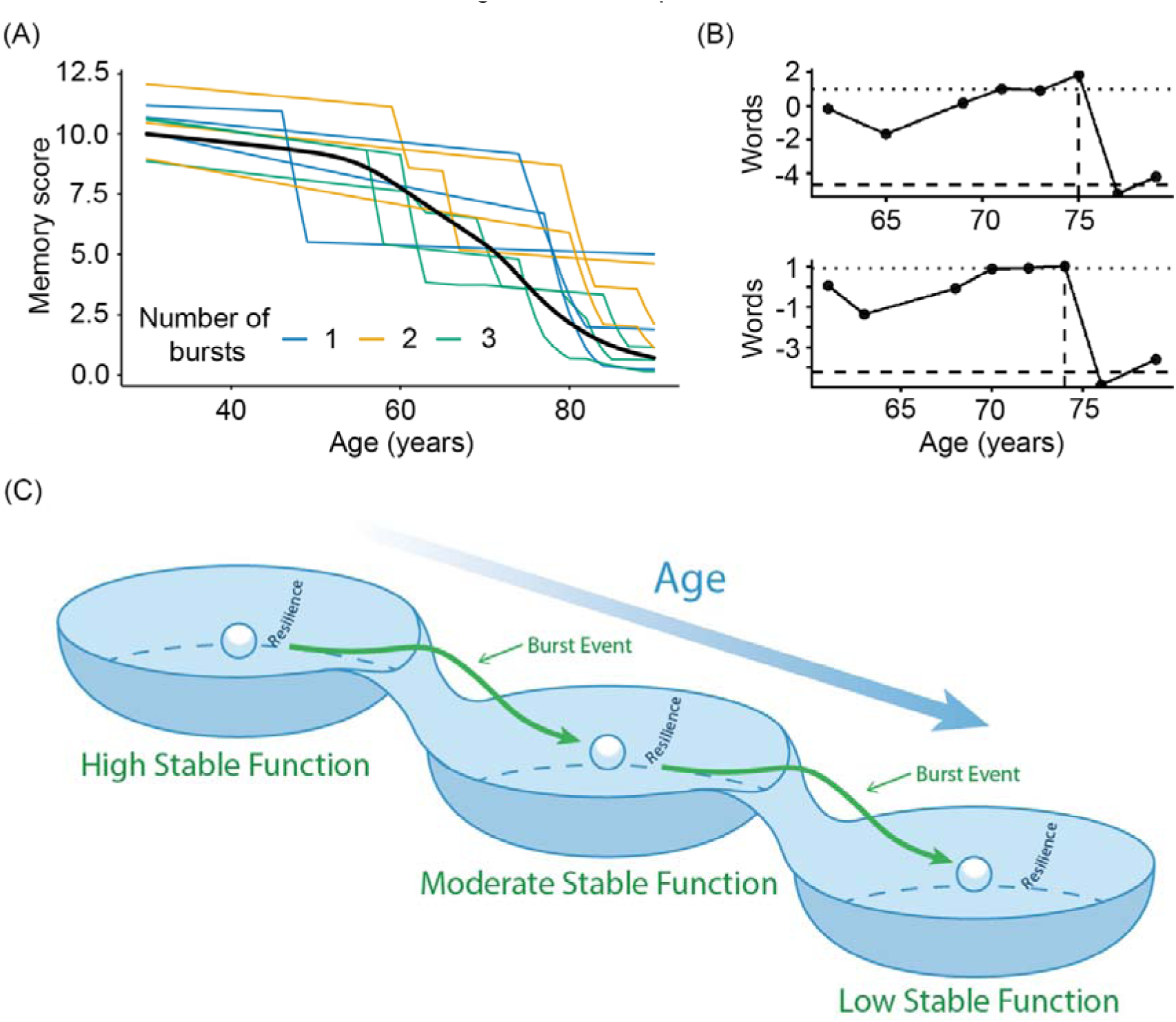
Cognitive resilience and burst-declines: A complex systems model of aging. A: Aggregating 5,000 individual “burst–decline” trajectories produced the smooth, age-normative episodic-memory curve (black). Nine randomly selected simulated curves are highlighted, color-coded by number of bursts. B: Examples of non-recovered burst declines in two real participants. Points and lines show practice-corrected recall residuals after subtracting each person’s Theil–Sen trend. The dotted horizontal line is the pre-burst mean residual; the dashed horizontal line is the post-burst mean residual; the dotted vertical line marks burst onset. These bursts meet three criteria: drop ≥ 1.5 words, slope ≤ −0.5 words/year, interval ≤ 3 years. A burst is labeled non-recovered if the best subsequent residual ≥ 2 years after onset remains below the 50% recovery threshold. C: The Cognitive Resilience Basins model illustrating burst–decline dynamics. This conceptual figure adapts the complex-systems “ball-in-a-cup” metaphor (basins of attraction) to explain heterogeneous, non-linear episodic-memory trajectories in aging. **Model components.** Basins of attraction (cups) represent discrete, locally stable states of cognitive function; basin depth and width index resilience, that is, how much perturbation can be absorbed before the system transitions to another state. The ball represents an individual’s current episodic-memory performance. Perturbations—internal or external stressors such as illness, vascular events, or cumulative allostatic load—can displace the system from equilibrium and precipitate burst events. **Trajectories.** The green trajectory illustrates a high-performing stable participant who remains within a deep, resilient basin. A sufficiently strong perturbation can trigger a rapid, non-linear drop (“burst”) as the system crosses a tipping threshold into a lower-capacity basin, followed by a new period of stability at a lower level. Consistent with our back-projection results, many late-life low-performing stable participants may reflect such post-burst stabilization rather than lifelong resistance to decline. With increasing age, basin walls gradually lower, reducing resilience and increasing the likelihood of burst transitions, consistent with greater prevalence of bursts in later life.

To test for such discontinuities in real data, we analyzed deviations from within-participant linear trends in SHARE. We identified “bursts” as intervals of fast (≤ −0.50 words/year), substantial (drop ≥ 1.5 points), and short (≤ 3 years) decline (Figure 8). Bursts were common and sizeable (median drop ∼3.4 points), and the majority (≈80–94%) did not recover to baseline. Naturally, stable performers experienced fewer bursts overall and were significantly less likely to exhibit non-recovered “big bursts” (≥2× each person’s residual SD) even after adjusting for exposure duration (OR ≈0.16, 95% CI 0.15–0.17). Permutation tests disrupting the temporal order of residuals confirmed that these patterns exceed chance expectations for all age bands (50–80+) (Section 9). Thus, it is possible that specific, time-structured loss events can drive episodic memory decline in addition to constant erosion, and stability in many appear to reflect the transient avoidance or containment of these bursts, which can help explain how some individuals remain stable for a decade while still performing at relatively low levels.

## Discussion

The present analyses yield two fundamental insights into cognitive aging. First, we confirm that long-term episodic memory stability is a real phenomenon: approximately one in ten older adults maintain stable performance over a decade, anchored in lower rates of brain atrophy. Second, this stability is often a transient state rather than a lifelong trait. These results help reconcile the descriptive concept of “maintenance” with a more mechanistic account of trajectories: low average decline can reflect at least two temporal architectures, a subgroup with genuinely stable performance over long intervals (‘trait-like stability’), and a more common pattern of extended plateaus punctuated by episodes of accelerated loss (‘state-like stability’). Back-projection analyses and individual trajectory modeling reveal that many of these stable performers have likely stabilized following earlier periods of decline. Thus, the monotonic, accelerating curves seen in group averages can mask a common punctuated, non-linear reality at the individual level, characterized by periods of resilience interspersed with bursts of loss. While a subgroup exhibits robust, multi-decadal stability, for the majority, stability may represent a temporary plateau. These findings motivate a refinement of cognitive aging models toward a complex-systems view of resilience and loss.

### Implications for models of cognitive aging

Our results challenge a view of cognitive decline in aging dominated by slow, monotonic erosion. We emphasize that this does not exclude gradual decline in many individuals; rather, our findings suggest that the temporal organization of change is often state-like, such that low average decline can arise from periods of relative stability punctuated by brief episodes of accelerated loss. We therefore support a model in which cognitive function behaves as a dynamical system, often visualized as a “ball in a cup” (Figure 8), that withstands perturbations until a tipping point triggers a critical transition (“burst”) to a lower functional state. This aligns with prior evidence of accelerated decline phases in preclinical dementia (22–24) and terminal drop (25), but extends the concept to the broader aging population, suggesting that stability is often a post-decline equilibrium rather than lifelong preservation.

This complex-systems framework of aging and resilience (14) resolves the paradox of our back-projection and mortality findings. The implausibly low historical projections for many stable performers suggest they have already undergone a state transition, settling into a lower but internally stable basin of attraction. Their stable performer status may therefore reflect current resilience (depth of the new basin) rather than a high baseline. Crucially, our mortality analysis also suggests that while high cognitive level is protective, the stability label itself captures a heterogeneous group, including fragile individuals likely residing close to a critical threshold. One plausible explanation is that stability can sometimes mark a post-event plateau: after an adverse health shock produces a step-like drop relative to an individual’s prior trajectory, performance may temporarily stabilize at a lower set-point, yielding near-zero slopes while simultaneously signaling elevated underlying risk. Thus, stability can coexist with underlying fragility, reinforcing the view that memory stability can reflect a transient state within dynamic trajectories rather than a fixed, protective trait. This dissociation - stability masking underlying vulnerability - is a hallmark of complex systems near or following a phase transition, analogous to “critical slowing down” in ecosystems or financial markets prior to collapse (26, 27). In this sense, “maintenance” may be viewed as a descriptive umbrella that pools trait-like stability and common state-like plateaus, helping to explain why maintainer prevalence estimates vary widely across designs and definitions.

Biologically, these downward shifts (“bursts”) likely represent moments where the cumulative burden of physiological dysregulation (allostatic load) exhausts the brain’s capacity (28). The lower rate of ’non-recovered’ bursts in stable performers suggests they might currently have superior network robustness or greater reserves. To capture these dynamics, future research can move beyond linear models toward latent-state approaches (e.g., Hidden Markov Models) that explicitly characterize aging as probabilistic transitions. This perspective aligns with Catastrophe Theory (29, 30), which mathematically describes how continuous pressure (e.g., accumulated pathology) can precipitate sudden functional shifts.

This dynamic view is further corroborated by the profile of the stable performers themselves. While stable performers were, on average, slightly healthier and had fewer depressive symptoms, broadly in line with previous studies (8, 10, 31, 32), effect sizes were uniformly small (33), and multivariate models added negligible discriminative power beyond age alone. Crucially, factors like education were strongly linked to level but weakly to slope (34–36). The absence of a sharply distinct ‘stable performer phenotype’ suggests that stability does not reflect a singular “super-aging” protection factor or a specific lifestyle profile. This pattern aligns with prior maintenance work in suggesting that “good aging” is rarely attributable to a single protective factor (10), but stability could instead often reflect the combined effect of many small advantages interacting with time-varying resilience.

The transient nature of memory stability implies that long periods of stability do not guarantee future protection, and individuals can shift from stability to decline over relatively short intervals. Consequently, risk screening based on single-timepoint performance, or even short-term stability, may overlook patients who are currently stable but critically vulnerable to future bursts, parallel to the “tipping point” view on neurodegeneration (24). Effective prevention may require identifying early-warning signals of resilience loss, such as slowed recovery from minor stressors, rather than waiting for gross decline. In other complex systems, small increases in variability or slowed recovery from perturbations can serve as early-warning signals of impending transitions (29, 30), and analogous markers in cognitive, behavioral, or neurophysiological data may help identify individuals approaching a decline episode.

### The neural foundation of memory stability

The stable performer construct was anchored in neurobiology by demonstrating that it is not merely a behavioral classification but tracks with brain structural change. Stable participants exhibited reduced atrophy across large regions of the cortex and subcortex, consistent with the brain maintenance framework (5, 37). Here we complement this framework with temporal dynamics, showing that the same underlying structural preservation can support either enduring stability or state-like plateaus within punctuated trajectories. Crucially, the effect sizes for structural preservation were substantially larger than for self-reported health and lifestyle factors, with a 1 SD reduction in volume loss associated with almost 50% higher odds of stability. Regions driving the effect included the hippocampus and other temporal lobe areas, structures central to episodic memory, but associations extended well beyond canonical memory regions. This topography aligns with recent evidence that age-related memory decline reflects both a specific vulnerability of the medial temporal lobe and a global factor of brain aging (37, 38) potentially driven by distinct biological modes of structural change (39–41). The anatomically widespread effects could suggest that atrophy in networks supporting attention, executive control, and other non-memory processes can erode memory stability indirectly, in line with meta-analytic evidence that roughly 60% of the variance in cognitive change is shared across functions (42). This implies a clear prediction: if memory stability relies on partly global brain preservation, these individuals should exhibit superior stability in other cognitive domains as well, while retaining a relative advantage in episodic memory. Determining whether stable participants are “domain-general brain maintainers” or a specific “hippocampal-sparing” subgroup will be essential for mapping the biological boundaries of healthy aging (38, 43).

While structural preservation is a hallmark of memory stability, the temporal dynamics of brain and cognition need not be identical. Our back-projection analysis for the hippocampus found only a modest skew at age 55, contrasting with the dramatic cognitive paradox. This dissociation is coherent with a complex-systems view where cognitive bursts arise from the non-linear interaction of multiple physiological and environmental factors, of which macroscopic atrophy is only one slowly changing component. It is plausible, and predicted by the model, that stable participants may experience transient periods of accelerated memory decline (bursts) without simultaneous accelerated volumetric loss. For example, a “burst” triggered by psychosocial stress or late-life depression (44), could precipitate a rapid state transition in attention and performance (43) while the accompanying change in hippocampal atrophy rate remains comparatively modest or lagged (45). Thus, structural brain maintenance provides the capacity (deep basin) for cognitive stability, but specific functional trajectories are shaped by the dynamic interplay of acute perturbations and reserve.

### Limitations

First, while our analyses spanned up to 24 years, lifelong trajectories are inferred rather than fully observed. Second, “burst-like” change is inherently challenging to capture in standard longitudinal data; while our simulations and back-projections provide converging evidence for a burst–decline process, the specific timing of acute loss events remains defined indirectly. Finally, the neuroimaging analysis was restricted to a subset with shorter follow-up and volumetric measures. Future research with denser (46), multi-modal sampling will be essential to map the precise coupling between cognitive bursts and corresponding non-linear dynamics in brain structure and function.

### Conclusion

The results suggest that episodic memory in later life is neither uniformly declining nor well captured by a single monotonic trajectory. Instead, many individuals appear to follow punctuated trajectories - extended plateaus of relative stability interrupted by bursts of loss, with a distinct subgroup of approximately one in ten older adults showing genuine, robust stability over a decade embedded within these heterogeneous paths. We show that this stability is real, biologically grounded in brain preservation, and only modestly predicted by conventional risk factors. Going forward, the field should move beyond searching for a static ‘*stability phenotype*’ to developing models that can resolve the dynamics of these critical transitions. In this sense, our ambition is that the present work can contribute to a broadened perspective, from asking who declines to also asking when, how, and under what conditions resilience fails or stability is sustained.

## Acknowledgement

LCBC is supported by the European Research Council under grant agreements no. 283634 and no. 725025 (to A.M.F.) and no. 313440 (to K.B.W.), as well as the Norwegian Research Council (325878, 262453 to A.M.F.; 325001, 301395, 239889 to K.B.W.; 249931 to A.M.F & K.B.W.; 324882 to DVP; 325415 to HG), the National Association for Public Health’s dementia research program, Norway (to A.M.F.), and the University of Oslo through the UiO:Life Science convergence environment (to A.M.F). Betula is supported by a scholar grant from the Knut and Alice Wallenberg foundation to L.N. See SI for funding of the different data sources.

## Author contributions

AMF, LN, KBW designed the work. AMF, EOSG, DV-P & AL analysed data. AMF drafted the manuscript. All authors critically revised and approved the final version.

