## Supplementary material for "Punctuated memory change: The temporal dynamics and brain basis of memory stability in aging": Online Methods

The research complies with all relevant ethical regulations, and all participants provided informed consent. The main project was approved by the Norwegian Regional Committee for Medical Research Ethics South (approval 8122), and each sub-study was approved by the relevant ethical review board, as specified in SI Section 13.

**Samples**

Each sample is thoroughly described in multiple previous publications, so only the main features relevant for the present study is presented in the following. An overview of the samples is presented in Online Methods Table 1.

| Cohort | Sample | n | Obs | Age | Waves | Interval |
| --- | --- | --- | --- | --- | --- | --- |
| SHARE | Full | 155606 | 443734 | 67.4 (40–111) | 2.9 (1–8) | 5.2 (0–18) |
| SHARE | >=4 sessions | 49602 | 258025 | 68.1 (40–104) | 5.2 (4–8) | 11.1 (6–18) |
| HRS | Full | 38448 | 229234 | 66.7 (40–109) | 6.0 (1–13) | 10.4 (0–25) |
| HRS | >=4 sessions | 24969 | 200696 | 67.3 (40–107) | 8.0 (4–13) | 14.7 (4–25) |
| Betula | Full | 4423 | 9731 | 62.5 (25-100) | 2.2(1-6) | 6.0 (0-25) |
| Betula | >=4 sessions | 884 | 4344 | 61.4 (34-100) | 4.9(4-6) | 19.5 (14-25) |
| MRI | Full | 12954 | 45367 | 72,5 (40-92) | 3.5 (1-31) | 4.1 (0-31.6) |
| MRI | >=4 sessions | 5224 | 32673 | 73,5 (40-92) | 8.3 (4-31) | 8.0 (2-31.6) |
| Total | Full | 211431 | 728066 | 25-111 | 1-13 | 0-31.6 |
| Total | >=4 sessions | 80679 | 495738 | 34-107 | 4-31 | 4-31.6 |

***Online Methods Table 1. Sample characteristics.*** *SHARE – The Study of Health and Retirement in Europe; HRS – The Health and Retirement Study (US); Betula - the Betula Prospective Cohort Study (Swe); MRI – Subsamples with MRI and memory tests, note that these numbers refer to the memory observations across these samples, which are greater than the numbers when restricted to only participants with longitudinal MRI data (see Online Methods Table 2). Most analyses were restricted to participants ≥ 50 years.*

**SHARE – sample and test characteristics**

The Survey of Health, Ageing and Retirement in Europe is a research infrastructure for studying the effects of health, social, economic and environmental policies over the life-course of European citizens and beyond (<https://share-eric.eu/>) (1). We used the SHARE easySHARE release (rel9_0_0) for the main analyses, restricting to respondents with baseline age 50–90 years. SHARE contains observations of individuals from 50 years of age from 28 countries, recruited to be representative of the population in each country. Data for the present analyses was extracted from *easy*SHARE (release 8.0.0, February 10^th^ 2022, doi:10.6103/SHARE.easy.800), see (2, 3) for methodological details. The *easy*SHARE release 8.8.0 is based on SHARE Waves 1, 2, 3, 4, 5, 6, 7, and 8 (DOIs:10.6103/SHARE.w1.800, 10.6103/SHARE.w2.800, 10.6103/SHARE.w3.800, 10.6103/SHARE.w4.800, 10.6103/SHARE.w5.800, 10.6103/SHARE.w6.800, 10.6103/SHARE.w7.800, 10.6103/SHARE.w8.800) (1, 4).

Memory was assessed with a 10-word verbal recall test. The word list is read out loud to the participants, and then recall is tested immediately after the presentation (Recall 1) and then after a delay of approximately 5 minutes (Recall 2). The sum of immediate and delayed recall was used in the analyses. Multiple versions of the lists are assigned to the respondents (5). There were no ceiling effects, which is important when assessing longitudinal change for the best-performing participants. There were some floor effects for recall 2, but less for recall 1, suggesting that we can estimate longitudinal chance well for most baseline levels of memory (6).

In addition to the memory measures, we extracted the variables age, sex, birth year, education (based on the International Standard Classification of Education 1997), and country of current residency, as well as several functional, social and health-related variables with possible associations with memory function.

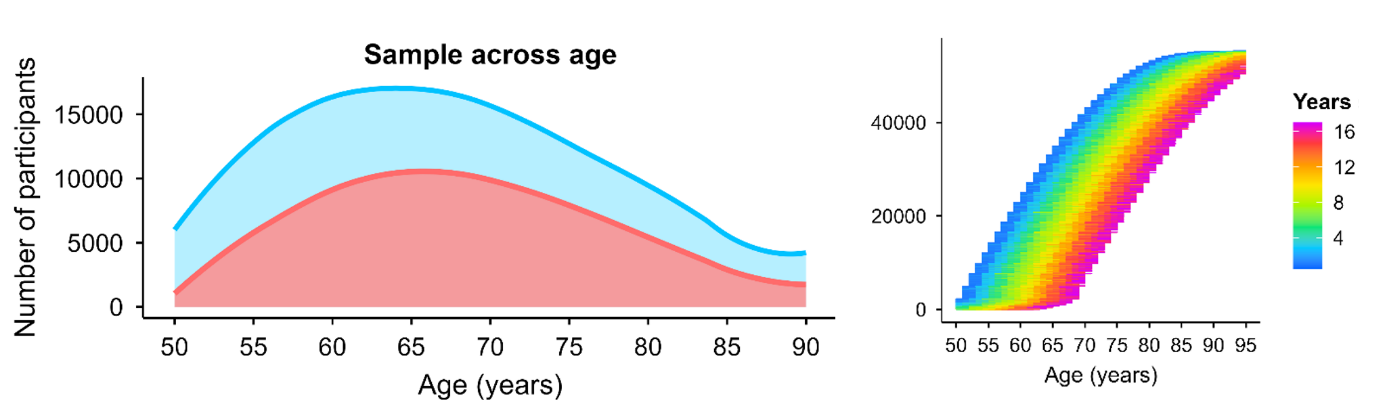

***Online Methods Figure 1. SHARE sample distribution.*** *Left panel: LOESS [locally estimated smoothing spline] smoothed age distribution of participants from 50–90 years. The aqua area shows all unique participants observed at each age and the coral area overlays the subset with ≥4 visits. Right panel: Spaghetti overview of within-person age coverage in the core sample. Each horizontal line represents one participant, stacked by baseline age. Line color encodes time since baseline.*

**Betula – sample and test characteristics**

The Betula project (Umeå, Sweden) is a prospective longitudinal study on aging, memory, and dementia, which used a population-based sampling of healthy middle-aged and older adults for recruitment (7, 8). Participants were followed with repeated in-person assessments at regular intervals, with comprehensive neuropsychological testing and detailed health and demographic characterization collected at each wave. The study was designed to support both normative cognitive aging research and investigations of preclinical and clinical dementia, with long follow-up and linkage to clinical evaluations enabling identification of incident cognitive impairment and dementia outcomes. Standardized test protocols and consistent procedures across waves provide high-quality longitudinal data for examining individual differences in level and change in memory and related cognitive functions.

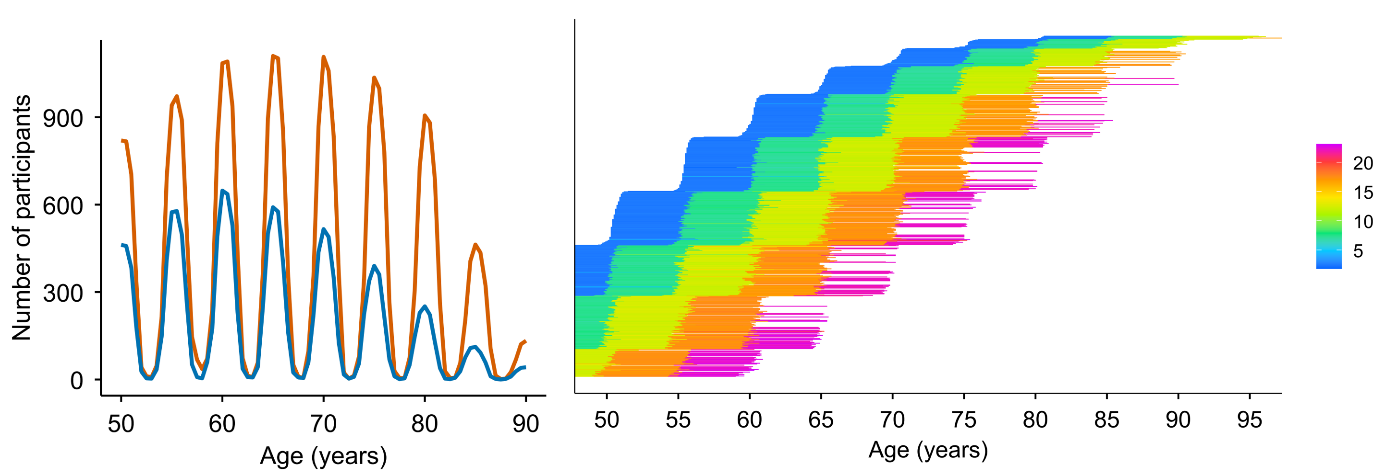

***Online Methods Figure 2. BETULA sample distribution.*** *Left panel: LOESS smoothed age distribution of participants from 50–90 years. The orange line shows all unique participants observed at each age and the blue line overlays the subset with ≥4 visits. Right panel: Spaghetti overview of within-person age coverage in the core sample. Each horizontal line represents one participant, stacked by baseline age. Line color encodes time since baseline, indexed by the color bar.*

**HRS – sample and test characteristics**

The Health and Retirement Study (HRS) is a longitudinal panel study designed to investigate health, cognitive functioning, retirement, and aging in the United States, based on a nationally representative sample of community-dwelling adults aged 50 years and older and their partners, with replenishment cohorts added over time to maintain population representation. Participants are followed at regular intervals with repeated interviews and assessments, providing harmonized measures of sociodemographic factors, health, and cognition across waves and enabling analyses of late-life trajectories and individual differences. For the present analyses, we used the RAND HRS Longitudinal File (1992–2022 release, Stata format), which provides cleaned and consistently coded variables across waves.

Episodic memory was assessed using immediate and delayed recall of a 10-word list; the list is presented orally and recall is tested immediately and after a short delay. To ensure comparability across follow-up, analyses were restricted to waves with 10-word lists (early waves 3–13, and later waves 14–15 where mode-specific variables were harmonized). At each wave, immediate and delayed recall scores were summed to form a composite episodic memory score (0–20). For later waves with separate mode-specific variables (FTF/Tel vs web), values were harmonized by coalescing the available mode-specific score. Participants contributed repeated assessments with age at interview as the time axis. We retained individuals with at least four valid observations after score construction and practice correction (≥4 waves), which defines the analysis sample for all stable performers and bout analyses.

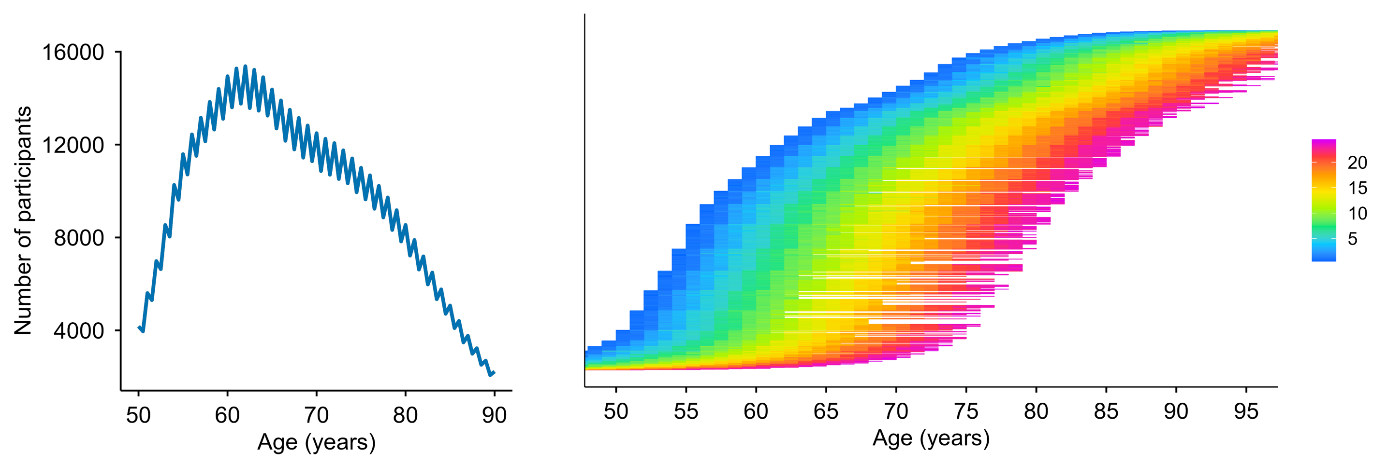

***Online Methods Figure 3. HRS sample distribution.*** *Left panel: LOESS smoothed age distribution of participants from 50–90 years. The blue line shows all participants with ≥4 visits. Right panel: Spaghetti overview of within-person age coverage in the sample. Each horizontal line represents one participant, stacked by baseline age. Line color encodes time since baseline, indexed by the color bar.*

**MRI subsample – sample and test characteristics**

We analyzed longitudinal MRI data from 10 independent studies (n =1,978 participants aged > 50 with > 4 episodic-memory test sessions covering 16,342 tests and 7,206 scans). The datasets include the the Alzheimer’s Disease Neuroimaging Initiative (ADNI) database (<https://adni.loni.usc.edu>) (Mueller et al., 2005), AIBL (Ellis et al., 2009), BLSA (open subset) (Ferrucci, 2008), Cognorm (Idland et al., 2020), HABS_HD (Petersen et al., 2025), LCBC (Walhovd et al., 2016), Mayo (Roberts et al., 2008), PREVENT-AD (Breitner et al., 2016; Tremblay-Mercier et al., 2021), OASIS3 (LaMontagne et al., 2019), and WRAP (Sager et al., 2005) datasets. The initial dataset included individuals with 1 to 14 MRI acquisitions with longitudinal structural MRI data spanning up to 15.8 years. Similarly, memory assessments range from 1 to 30 observations per individual with a follow-up up to 31.6 years. An overview of the studies included are shown in Online Methods Table 2 and 3, and a more thorough description of each is provided in the SI Section 12.

| Cohort | Sample | n | Obs | Age | Waves | Interval |
| --- | --- | --- | --- | --- | --- | --- |
| ADNI | Full | 821 | 3769 | 76,6 | 4,6 | 3,98 |
| ADNI | >=4 sessions | 447 | 3097 | 77,6 | 6,9 | 6,4 |
| AIBL | Full | 608 | 1260 | 73,7 | 2,1 | 1,74 |
| AIBL | >=4 sessions | 115 | 535 | 73,7 | 4,7 | 5,75 |
| BLSA | Full | 132 | 396 | 82,7 | 3 | 3,27 |
| BLSA | >=4 sessions | 46 | 237 | 83,9 | 5,2 | 5,98 |
| HABS_HD | Full | 2832 | 4576 | 66,4 | 1,6 | 1,44 |
| HABS_HD | >=4 sessions | 131 | 524 | 67,2 | 4 | 6,56 |
| MCSA | Full | 5609 | 20936 | 74,7 | 3,7 | 3,72 |
| MCSA | >=4 sessions | 2742 | 15988 | 75,3 | 5,8 | 6,47 |
| Oasis3 | Full | 923 | 5985 | 74,1 | 6,5 | 7,32 |
| Oasis3 | >=4 sessions | 623 | 5359 | 74,6 | 8,6 | 9,78 |
| COGNORM | Full | 114 | 677 | 75,9 | 5,9 | 5,28 |
| COGNORM | >=4 sessions | 100 | 651 | 75,8 | 6,5 | 5,88 |
| PreventAD | Full | 348 | 1916 | 66,1 | 5,5 | 4,38 |
| PreventAD | >=4 sessions | 265 | 1710 | 66,3 | 6,5 | 5,17 |
| LCBC | Full | 256 | 559 | 65,4 | 2,2 | 4,94 |
| LCBC | >=4 sessions | 53 | 215 | 66 | 4,1 | 12,26 |
| Wrap | Full | 1094 | 5021 | 65,1 | 4,6 | 10,93 |
| Wrap | >=4 sessions | 702 | 4357 | 65,7 | 6,2 | 15,4 |

***Online Methods Table 2. Sample characteristics for the MRI subsample****. The numbers refer to participant characteristics tied to the memory tests. The essential numbers for the number of MRI scans are provided in the main text. ADNI – Alzheimer’s Disease Neuroimaging Initiative; AIBL – Australian Imaging, Biomarker & Lifestyle Study of Ageing; BLSA – Baltimore Longitudinal Study of Aging; HABS-HD - Health and Aging Brain Studies–Health Disparities; MCSA – Mayo Clinic Study of Aging / Mayo Alzheimer’s Disease Research Center cohorts; OASIS3 – Open Access Series of Imaging Studies (OASIS-3); COGNORM – Oslo University Hospital studies; PREVENT-AD – Pre-symptomatic Evaluation of Experimental or Novel Treatments for Alzheimer Disease; LCBC – Center for Lifespan Changes in Brain and Cognition cohorts; WRAP – Wisconsin Registry for Alzheimer’s Prevention.*

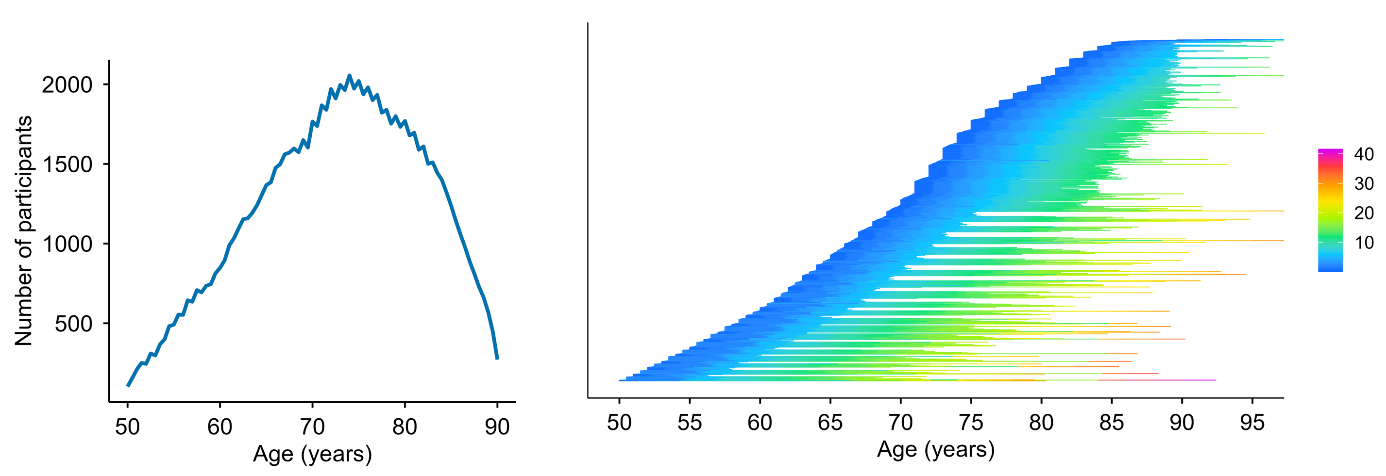

***Online Methods Figure 4. MRI subsample distribution.*** *Left panel: LOESS smoothed age distribution of participants from 50–90 years. The blue line shows all participants with ≥4 visits. Right panel: Spaghetti overview of age coverage for each person in the sample. Each horizontal line represents one participant, stacked by baseline age. Line color encodes time since baseline, indexed by the color bar.*

We used all available memory tests in each MRI subsample and computed the principal component (PC) at baseline on the z-scaled subtests, representing memory function separately within each sample (mean = 0, SD = 1). The included memory tests are shown in Online Methods Table 3. Structural T1-weighted (T1w) MPRAGE and FSPGR scans were collected using 1.5 and 3T MRI scanners. Information regarding scanners and scanner parameters across datasets are presented in Online Methods Table 3. Data was converted to BIDS (Gorgolewski et al., 2016) and preprocessed using the longitudinal FreeSurfer v.7.1.0 stream (Reuter et al., 2012) for cortical reconstruction and volumetric segmentation of the structural T1w scans (Dale et al., 1999; Fischl et al., 1999). Data was tabulated based on the *Desikan* (cortical; *N* = 34 regions per hemisphere) (Desikan et al., 2006) and *aseg* (subcortical, *N* = 18 regions) atlases (Fischl et al., 2002).

| Dataset | Memory test | Scanner | Field | Sequence parameters |
| --- | --- | --- | --- | --- |
| ADNI | ADNI-MEM^1^ | Multisite | 1.5/  3.0 | https://adni.loni.usc.edu/methods/documents/mri-protocols/ |
| OUS | CERAD short and long delay recall | Siemens Avanto | 3.0 | MPRAGE. TR: 2400 ms; TE: 3.79 ms, TI: 1000 ms; flip angle 8°, slice thickness: 1.2 mm, FoV 240 x 240, 160 slices. |
|  |  | SiemensPrisma | 3.0 | MPRAGE. TR: 2400 ms; TE: 2.22 ms, TI: 1000 ms; flip angle 8°, slice thickness: 0.8 mm, FoV 240 x 256, 208 slices, iPat = 2. |
| HABS_HD | SEVLT (learning & long delay recall); Logical memory (immediate & delay recall); | SiemensSkyra | 3.0 | MPRAGE. TR: 2300 ms; TE: 2.93 ms, TI: 900 ms; flip angle 9°, slice thickness: 1.2 mm, FoV 240 x 256, 176 slices. |
|  |  | Siemens Vida | 3.0 | MPRAGE. TR: 2300 ms; TE: 2.98 ms, TI: 900 ms; flip angle 9°, slice thickness: 1 mm, FoV 240 x 256, 208 slices. |
| LCBC | CVLT (learning, short & long delay recall) | SiemensAvanto | 1.5 | MPRAGE. TR: 2400 ms; TE: 3.79 ms, TI: 1000 ms; flip angle 8°, slice thickness: 1.2 mm, FoV 240 x 240, 160 slices. |
|  |  | SiemensSkyra | 3.0 | MPRAGE. TR: 2300 ms; TE: 2.98 ms, TI: 850 ms; flip angle 8°, slice thickness: 1 mm, FoV: 256 x 256, 176 slices. |
|  |  | SiemensPrisma | 3.0 | MPRAGE. TR: 2400 ms; TE: 2.22 ms, TI: 1000 ms; flip angle 8°, slice thickness: 0.8 mm, FoV 240 x 256, 208 slices, iPat = 2. |
| OASIS3 | Logical memory immediate | Vision | 1.5 | MPRAGE. TR: 9,7 ms; TE: 4.0 ms, TI: 20 ms; flip angle 10°, slice thickness: 1.25 mm, FoV: 256 x 256, 160 slices. |
|  |  | Siemens Sonata | 1.5 | MPRAGE. TR: 9,7 ms; TE: 3.9 ms, TI: 20 ms; flip angle 15°, slice thickness: 1 mm, FoV: 224 x 256, 160 slices. |
|  |  | Siemens Tim Trio | 3.0 | MPRAGE. TR: 2400 ms; TE: 3.1 ms, TI: 1000 ms; flip angle 8°, slice thickness: 1 mm, FoV 256 x 256, 176 slices. |
|  |  | Siemens Magnetom Vida | 3.0 | MPRAGE. TR: 2300 ms; TE: 2.3 ms, TI: 900 ms; flip angle 9°, slice thickness: 1.2 mm, FoV 240 x 256, 176 slices. |
|  |  | Siemens BioGraph mMR | 3.0 | MPRAGE. TR: 2300 ms; TE: 2.3 ms, TI: 900 ms; flip angle 9°, slice thickness: 1.2 mm, FoV 240 x 256, 176 slices. |
| PreventAD | RBANS list recall and learning, story immediate and delayed recall | Siemens Tim Trio | 3.0 | MPRAGE. TR: 2300 ms; TE: 2.98 ms, TI: 900 ms; flip angle 9°, slice thickness: 1 mm, FoV 240 x 256, 176 slices. |
| Mayo | pzmemory^2^ | GE Discovery MR 750 | 3.0 | SPGR. TR: 7.4 ms; TE: 3.0 ms, TI: 900 ms; flip angle 8°, slice thickness: 1.2 mm, FoV 256 x 256, 166 slices. |
|  |  | GE Signa HDxt | 3.0 | SPGR. TR: 7.0 ms; TE: 2.8 ms, TI: 900 ms; flip angle 8°, slice thickness: 1.2 mm, FoV 256 x 256, 166 slices. |
|  |  | GE Signa HDx | 3.0 | SPGR. TR: 6.9 ms; TE: 3.0 ms, TI: 900 ms; flip angle 8°, slice thickness: 1.2 mm, FoV 256 x 256, 166 slices. |
|  |  | GE Signa Excite | 3.0 | SPGR. TR: 7.2 ms; TE: 3.1 ms, TI: 900 ms; flip angle 8°, slice thickness: 1.2 mm, FoV 256 x 256, 166 slices. |
| WRAP | Logical Memory short & long delay recall; RAVLT learning & long delay recall; BVMT learning & delay recall | GE Signa Premier | 3.0 | SPGR. TR: 8.2 ms; TE: 3.2 ms, TI: 450 ms; flip angle 12°, slice thickness: 1 mm, FoV 256 x 256, 156 slices. |
|  |  | GE Signa Excite | 3.0 | SPGR. TR: 8.4 ms; TE: 1.7 ms, TI: 600 ms; flip angle 10°, slice thickness: 1.2 mm, FoV 256 x 256, 124 slices. |
|  |  | GE Signa HDx | 3.0 | SPGR. TR: 6.6 ms; TE: 2.8 ms, TI: 900 ms; flip angle 8°, slice thickness: 1.2 mm, FoV 256 x 256, 166 slices. |
|  |  | GE Discovery MR 750 | 3.0 | SPGR. TR: 8.2 ms; TE: 3.2 ms, TI: 450 ms; flip angle 12°, slice thickness: 1 mm, FoV 256 x 256, 156 slices. |
| BLSA | CVLT (learning, short & long delay recall) | Philips Achieva | 3.0 | MPRAGE. TR: 6.5 ms; TE: 3.1 ms, TI: 0 ms; flip angle 8°, slice thickness: 1.2 mm, FoV 256 x 256, 170 slices. |
| AIBL | Logical Memory short & long delay recall | Avanto Siemens | 1.5 | MPRAGE. TR: 2300 ms; TE: 2.98 ms, TI: 900 ms; flip angle 9°, slice thickness: 1.25 mm, FoV: 240 x 256, 160 slices. |
|  |  | Verio Siemens | 3.0 | MPRAGE. TR: 2300 ms; TE: 2.98 ms, TI: 900 ms; flip angle 9°, slice thickness: 1.25 mm, FoV: 240 x 256, 160 slices. |
|  |  | Tim Trio Siemens | 3.0 | MPRAGE. TR: 2300 ms; TE: 2.98 ms, TI: 900 ms; flip angle 9°, slice thickness: 1.25 mm, FoV 240 x 256, 160 slices. |

***Online Methods Table 3.*** *Memory tests and MRI scanner acquisition parameters in each MRI subsample****.*** *Tests related to episodic memory included in the analyses for each sample****.*** *A PC was estimated based on the first time point for which multiple memory measures were available. RAVLT = Rey Auditory Verbal Learning Test; CVLT = California Verbal Learning Test; Logical memory = Memory subtest of the Wechsler Memory Scale. CERAD = Consortium to Establish a Registry for Alzheimer's Disease (CERAD) Word List Memory test. SRT = Buschke Selective Reminding Task. RBANS = Repeatable Battery for Assessment of Neuropsychological Status. Story recall = Story recall and recognition task of episodic memory from Wechsler Neuropsychological Battery. ^1^ADNI-MEM score was computed developed by (9) and consists of a composite score of memory which includes measures from Rey Auditory Verbal Learning Test (learning trials, list, recognition and recalls), Alzheimer Disease Assessment Scale (learning trials, recall, and recognitions), Mini-mental State Examination words, and Logical memory. ^2^pzmemory score was developed by (Knopman et al., 2015) and consists of a composite score of memory tests which includes measures from Logical memory (delayed recall), Weschler’s Visual Reproduction (delayed recall), and RAVLT (delayed recall). For MRI: TR = Repetition Time; TE = Echo Time; TI = inversion time; FoV = Field of View, iPat = in-plane acceleration. ^a^Two matched scanners.*

For datasets not provided in Brain Imaging Data Structure (BIDS) format, data was converted to BIDS(10). BIDS transformation of ADNI, AIBL, OASIS3, and HABS_HD data was performed with Clinica software (Routier et al., 2021; Samper-González et al., 2018). For sessions with multiple scans, data from the scanners were averaged. Briefly, the images were processed using the cross-sectional stream, which includes the removal of nonbrain tissues, Talairach transformation, intensity correction, tissue and volumetric segmentation, cortical surface reconstruction, and cortical parcellation. Next, an unbiased within-subject template space based on all cross-sectional images was created for each participant, using robust, inverse-consistent registration (Reuter et al., 2010). The processing of each time point was then reinitialized with common information from the within-subject template to increase reliability and statistical power. Except for the BETULA dataset, all data was preprocessed on the Colossus processing cluster, part of the Tjenester for Sensitive Data (TSD) (<https://www.uio.no/tjenester/it/forskning/sensitiv/>), University of Oslo (UiO).

For each sample, we first z-normalized all measures based on the first time point and the different available memory tests. When multiple measures were available, we estimated a main component using Principal Component Analysis (PCA; *prcomp*) with all measures at the first time point as inputs. Missing values were imputed using *imputePCA* from the *missMDA* r-package(Josse and Husson, 2016). Only for OASIS3, the imputed number of values was not negligible (> .5%).

***Statistical analyses***

Analyses were mainly conducted in R (v4.4+). False Discovery Rate (FDR) was controlled across comparisons by use of the Benjamini–Hochberg procedure.

*Memory analyses and practice corrections*

For each participant we created prior_tests, a 0-based visit counter (0, 1, 2, …) ordered by age at assessment. To adjust memory scores for practice (retest) effects while retaining age-related variation, we fit an ordinary least squares regression model predicting the raw composite recall score from a flexible spline function of age (natural spline, df = 8) and a monotonic, diminishing-returns retest term defined as log(1 + prior_tests). The estimated coefficient for the retest term was then used to remove the practice component from each observation: the practice-adjusted recall score equals the raw recall score minus the estimated retest coefficient multiplied by log(1 + prior_tests). By construction, no adjustment is applied at a respondent’s first test, and the magnitude of the correction increases monotonically with additional prior exposure while allowing diminishing returns. All downstream analyses used the practice-adjusted recall variable. Distributions of raw and practice-adjusted scores and the implied correction factors are shown in Online Methods Figure 5.

| SHARE | HRS |
| --- | --- |
| 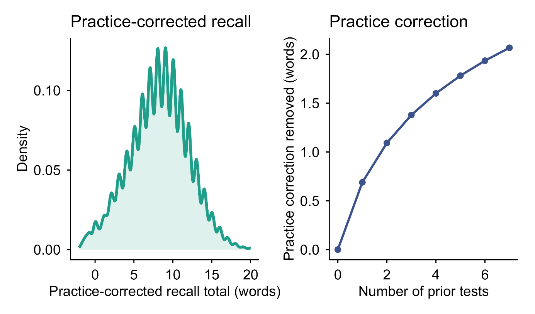 | 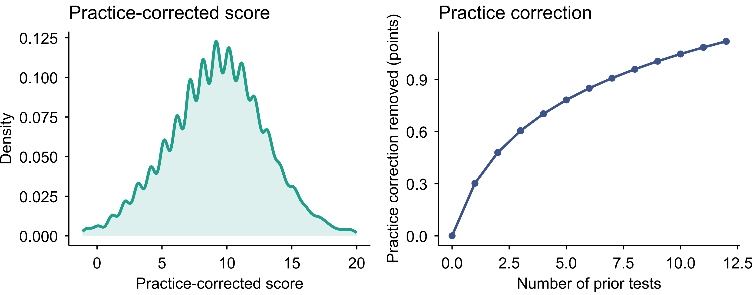 |
| BETULA | MRI subsample |
| 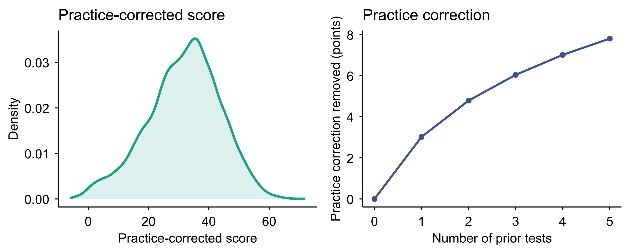 | 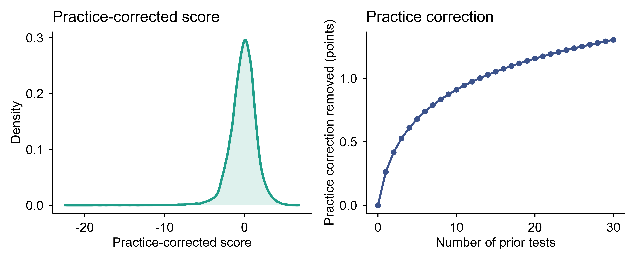 |

***Online Methods Figure 5****. Memory scores and practice effects pr sample. The left part of each panel shows distribution of practice-corrected memory composite scores pr sample. For SHARE and HRS, the test range is 0-20. For Betula, a memory composite score with range 0-76 is used. For the MRI subsample, a PCA was fitted to all memory-related tests in each subsample to extract a principal component (z-standardized). The right part of each panel shows the derived practice effect correction factor, starting at 0 for the first timepoint (0 previous tests).* *Practice effects were estimated separately for each subsample by fitting an ordinary least squares model with a flexible age adjustment and a monotone, diminishing-returns practice term, on the number of prior tests, using the full sample.*

Unless explicitly noted, we restricted the main analyses to respondents with at least four valid memory assessments (≥4 visits) to ensure stable within-person change estimation. We estimated each participant’s annual rate of change using Theil–Sen regression on the practice-corrected recall scores (Online Methods Figure 6) defined as the median of all pairwise slopes across that individual’s observations, yielding change in original recall units per year (Δrecall/Δyears). This estimator is robust to single-point anomalies and non-Gaussian noise and provides stable slope estimates in sparse longitudinal data. Uncertainty in the slope estimate was quantified non-parametrically using a leave-one-out (LOO) jackknife across waves. For each respondent with n observations, we recomputed the Theil–Sen slope n times, each time excluding one observation, and summarized the resulting distribution using empirical lower-bound quantiles (e.g., the 5th percentile, lb95, and the median, lb50). These lower-bound summaries down-weight classifications driven by any single unusually high or low observation (Online Methods Table 4). To avoid borderline flips due to numerical noise, we applied a small tolerance (τ = 0.01 units/year) when comparing lower-bound estimates to the thresholds used in the conservative stable performers rule (see stable performers definition).

| 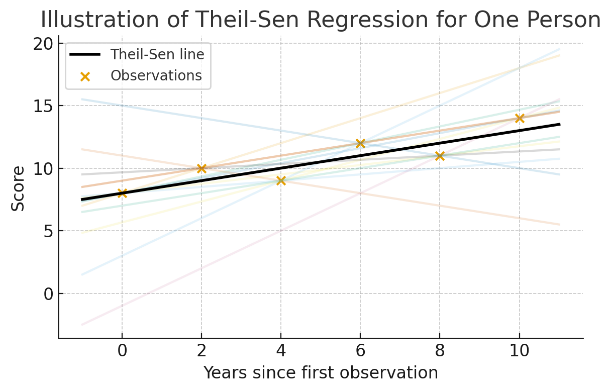 | 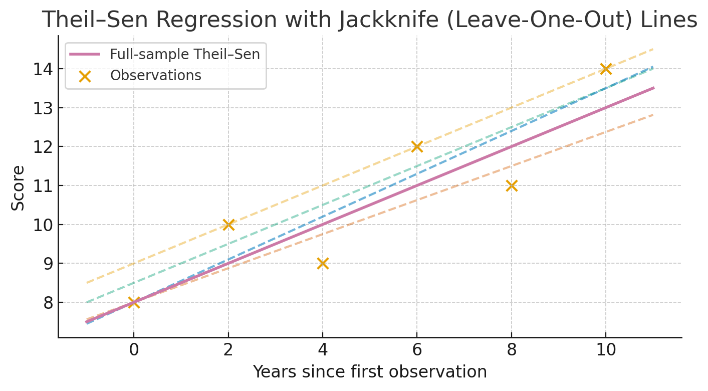 |
| --- | --- |

***Online Methods Figure 6. Theil-Sen regression for robust slope estimation with Jackknife Leave-One-Out.*** *Left: For each participant, we computed all pairwise rates of change in recall (Δrecall/Δyears) across their repeated assessments and defined the individual annual slope as the median of these pairwise slopes (Theil–Sen estimator). With n observations, the number of pairwise slopes is n(n−1)/2 (e.g., 4 observations → 6 slopes; 6 observations → 15 slopes; 8 observations → 28 slopes). Right: Uncertainty in the individual slope estimate was quantified using a leave-one-out jackknife over that participant’s observations: the Theil–Sen slope was recomputed n times, each time excluding one observation, yielding a distribution of leave-one-out slopes from which empirical lower-bound quantiles (e.g., the 5th percentile) were derived. This procedure reduces sensitivity to any single unusually high or low observation and supports conservative classification.*

| LB level | N (visits) | Max # slopes ≤ T if LB > T | Max % slopes ≤ T | Min # slopes > T | Min % slopes > T |
| --- | --- | --- | --- | --- | --- |
| 95 | 4 | 1 | 25% | 3 | 75% |
| 75 | 4 | 1 | 25% | 3 | 75% |
| 50 | 4 | 2 | 50% | 2 | 50% |
| 95 | 6 | 1 | 17% | 5 | 83% |
| 75 | 6 | 2 | 33% | 4 | 67% |
| 50 | 6 | 3 | 50% | 3 | 50% |
| 95 | 8 | 1 | 12% | 7 | 88% |
| 75 | 8 | 2 | 25% | 6 | 75% |
| 50 | 8 | 4 | 50% | 4 | 50% |

***Online Methods Table 4*.** *Number of Jackknife slopes violating the lower bound slope criterion as a function of LB level and number of visits.*

The distribution of practice corrected scores, longitudinal and cross-sectional age-relationships per sample are shown in Methods Figure 7.

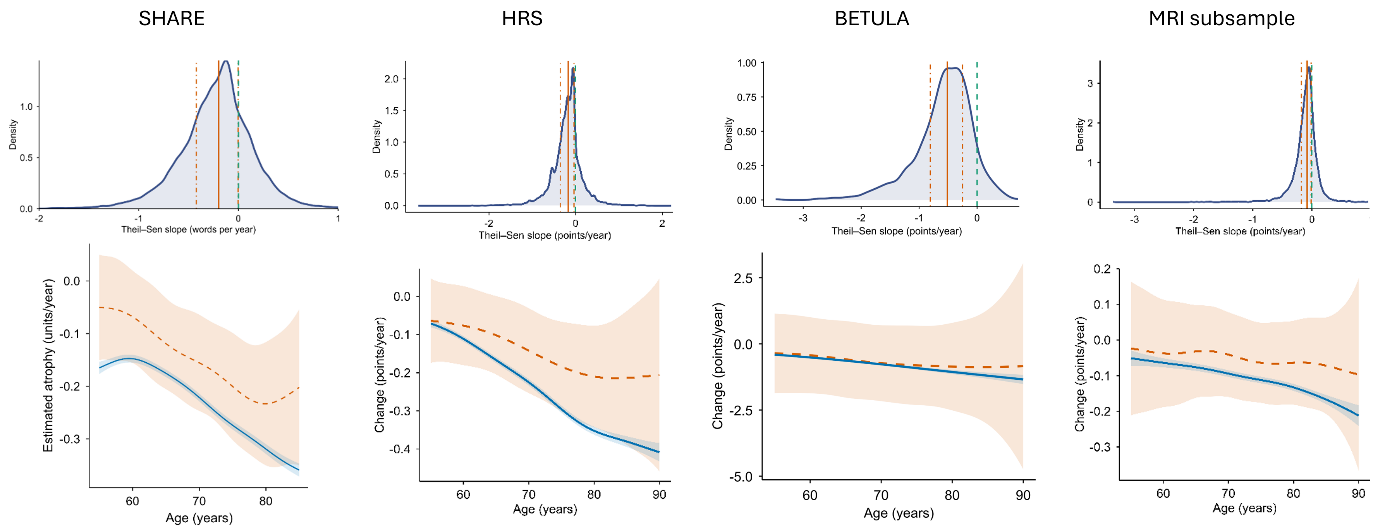

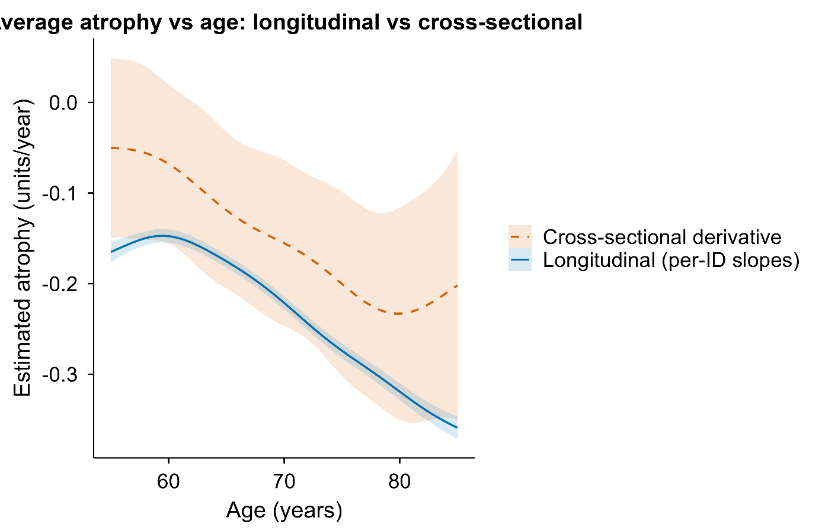

***Online Methods Figure 7. Distribution of Theil–Sen slopes and comparison of longitudinal and cross-sectional age trends across samples.*** *Top row: Distribution of individual annual memory-change estimates (Theil–Sen slopes) based on practice-corrected recall. The green dashed line marks zero change (flat slope). The orange solid line marks the median slope (−0.197 words/year; 95% CI −0.200 to −0.192), and the orange dot–dash lines indicate the interquartile range (25th and 75th percentiles). Bottom row: Longitudinal and cross-sectional age trends. The solid blue curve shows the longitudinal trend obtained by smoothing participants’ Theil–Sen slopes over median age using a generalized additive model (GAM). The dashed red curve shows the cross-sectional instantaneous rate of change, computed as the derivative of a GAM fitted to practice-corrected recall as a function of age using all observations. Shaded ribbons denote 95% confidence intervals. Units are words recalled per year for SHARE and HRS, points per year on the Betula composite scale, and standardized units (SD/year) for the MRI subsample.*

*Stable performers definition (primary and sensitivity thresholds)*

Participants were classified as stable performers using robust within-person slope estimates based on Theil–Sen regression applied to practice-corrected recall. For each participant, we summarized uncertainty in the slope using leave-one-out jackknife slopes and extracted empirical lower-bound quantiles (LB), such as the 5th percentile (LB95) and the median (LB50). A participant was classified as a stable performer under threshold δ if the relevant jackknife lower bound exceeded δ (with a small tolerance for the conservative rule). The two primary definitions were:

Conservative: LB95 ≥ 0 − τ, with τ = 0.01 units/year.

Lenient: LB50 ≥ −0.10 units/year.

The conservative definition therefore requires strong evidence that the participant’s slope is not negative even under single-wave perturbations, whereas the lenient definition allows small average decline (up to 0.10 units/year) while requiring that the median jackknife slope remains above the threshold. By construction, these rules are nested and monotonic: all conservative stable performers are also classified as lenient stable performers. In sensitivity analyses, we systematically varied the slope threshold δ (0.00, −0.25, −0.50, −0.75, −1.00 units/year) and the jackknife lower-bound quantile (95th, 75th, and 50th percentiles) to evaluate robustness of results to classification stringency. Note that “units/year” refer to the native scale of each outcome: words/year for SHARE and HRS, points/year for Betula’s composite memory score, and standardized units (SD/year) for the MRI subsample.

*Simulation experiment*

To estimate how often individuals would satisfy the stable performers criteria by chance, under the assumption that everyone follows the typical age-related decline and that apparent stability can arise from noise and sparse sampling, we conducted Monte Carlo simulations that preserve each participant’s observed visit ages and within-person noise structure.

First, we quantified each participant’s noise level as the within-person residual standard deviation around their own Theil–Sen line fitted to the practice-corrected recall scores. If a participant’s residual SD was not finite or equaled zero, we replaced it with the sample median residual SD. Second, we estimated the age-typical slope of memory change by fitting a generalized additive model (GAM) to the observed per-person Theil–Sen slopes as a smooth function of median age (REML estimation). This model provided an expected “group-typical” slope for each participant at their median age. Third, for each simulation replicate, we generated a synthetic memory trajectory for every participant on that person’s actual assessment ages. Each simulated trajectory followed the participant’s expected age-typical slope and included Gaussian noise with standard deviation set to that participant’s estimated residual SD. A robust intercept was used to anchor the simulated series at a realistic level for that participant (so that only the slope and noise determine whether stable performance is detected).

Finally, we applied the full stable performers pipeline to each simulated dataset exactly as in the observed data. This included recomputing each participant’s Theil–Sen slope and leave-one-out jackknife lower-bound summaries, applying the same stable performers rules (including the conservative tolerance), and then estimating stable performers prevalence as a function of age using the same moving-window procedure as in the observed analyses (ages 55–90, step 0.5 years, half-window 1.5 years, minimum n=30). For each age point, we summarized the simulation-based prevalence distribution across replicates using the median and interquartile range to form the null envelope. Because the person-specific noise SDs were estimated from residuals around each person’s own Theil–Sen line rather than around the group slope, the null model is conservative in that it can slightly inflate noise in individuals whose true trajectories deviate from the group-typical decline.

We ran 200 simulation replicates on the practice-corrected data. The simulations provide the expected prevalence of “stable performers” under a no-stable performers scenario, given the real sampling schedule and noise. Using the observed and simulated prevalence curves, we computed two closely related quantities. First, we estimated the excess prevalence of stable performers above chance (observed prevalence minus null prevalence) as a function of age. Second, to interpret the observed stable performers prevalence in terms of signal versus false positives, we computed a PPV-style estimate of the fraction of observed stable performers that exceed what is expected by chance at each age, using the null prevalence as the false-positive rate and an assumed sensitivity for detecting true stable performers (Se = 0.80). To provide conservative and lenient bounds, we propagated uncertainty from the null distribution by using the upper or lower null quantiles in the correction step (upper for conservative, lower for lenient), consistent with the bounds defined above.

To evaluate robustness, we repeated this simulation-based calibration across a grid of stable performers definitions by varying the slope threshold (δ) and the jackknife lower-bound requirement (LB quantile). Plotting observed prevalence against the simulation envelope showed that the observed curves exceeded the null envelope across the tested rule set (SI Figure X), demonstrating a stable performers signal beyond chance. As expected, stricter criteria (less negative δ and more conservative LB quantiles) increased confidence that any single identified stable performer reflects true signal (higher PPV), whereas more lenient criteria increased sensitivity and yielded larger observed–null separation (Online Methods Figure 8), at the cost of including more chance classifications. Based on this quantitative calibration and qualitative inspection of individual trajectories, we interpreted the population stable performers prevalence as lying between the lower-bound estimates from the conservative rule and the upper-bound estimates from the lenient rule. See *Online Methods Figure 9* for individual examples.

| 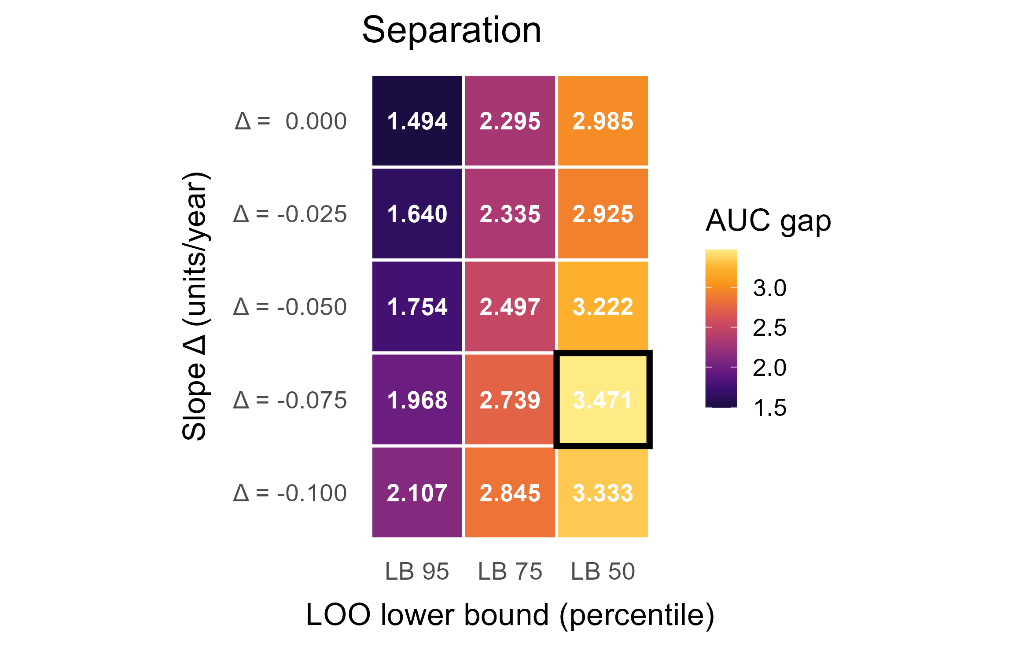 | 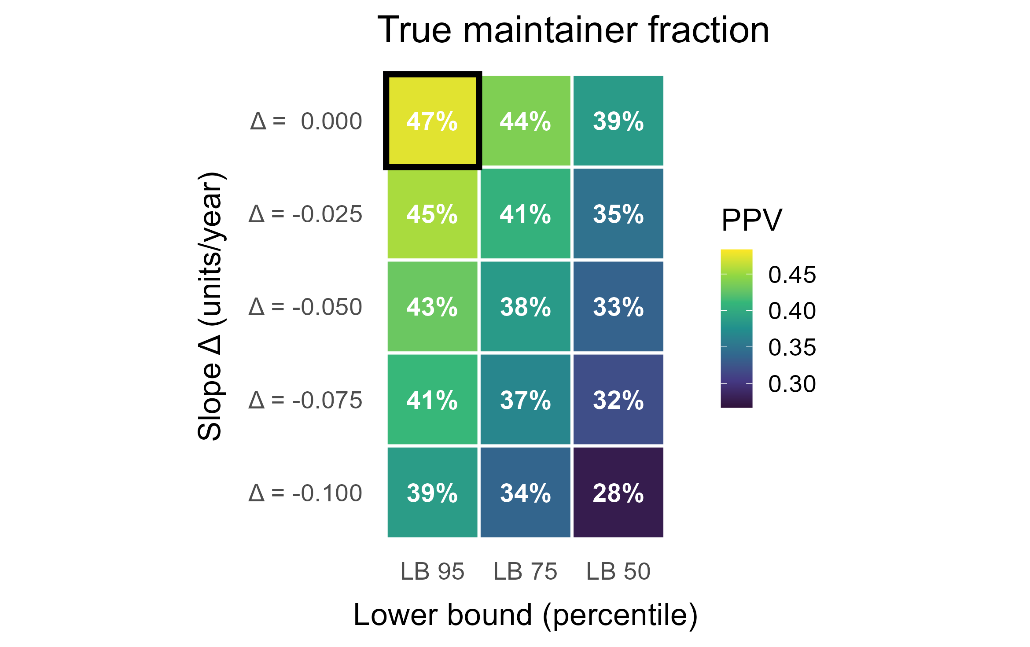 |
| --- | --- |

***Online Methods Figure 8.*** ***Calibrating stable performers definitions against a simulation-based null.*** *Left panel: Heatmap of an age-integrated separation metric quantifying how strongly the observed stable performers prevalence departs from chance for each rule definition. Separation is summarized as the area between the observed and null prevalence curves integrated over age (AUC gap); larger values indicate greater deviation from the null expectation. Right panel: Heatmap of the weighted mean positive predictive value (PPV) across age for each stable performers definition, indexed by the slope threshold (Δ; rows) and the jackknife lower-bound quantile (LB; columns). PPV represents the estimated fraction of observed stable performers that exceeds the simulation-based null prevalence (i.e., is likely genuine signal rather than noise-driven classification), using the same stable performers rule applied to each simulated dataset. Numeric labels within tiles report the corresponding statistic (PPV: 0–1; AUC gap: arbitrary units), and the highest value in each panel is indicated.*

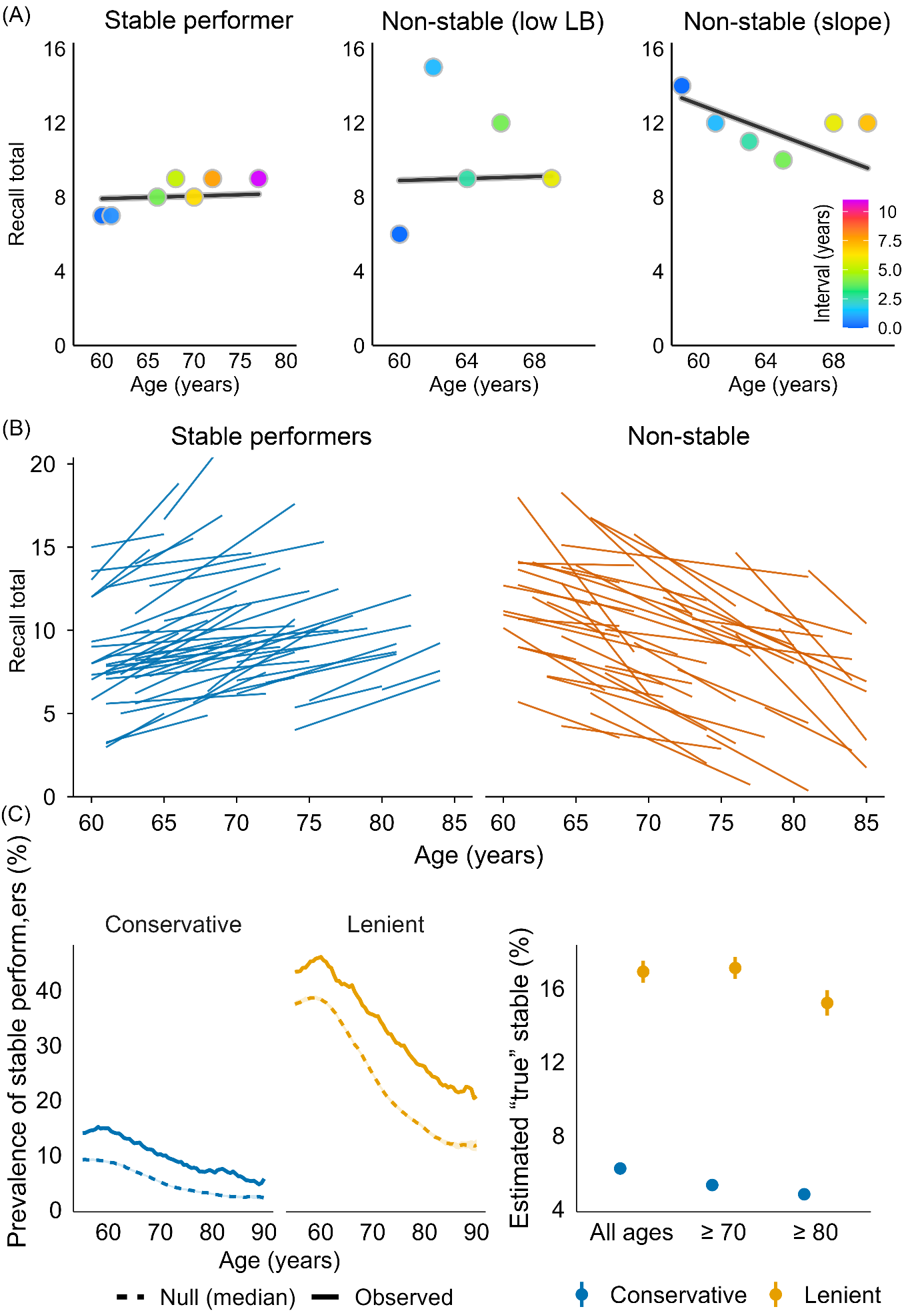

***Online methods Figure 9. Individual trajectories and randomly sampled longitudinal slopes in SHARE***. A: Left – Stable performer: near-flat Theil–Sen slope (> 0) with a 95% jackknife lower bound ≥ 0. Middle – Uncertain (non-stable performer): near-flat Theil–Sen slope (> 0) but the jackknife lower bound includes < 0. Right - Non-stable performer: negative slope (< 0) with high-confidence lower bound. B: Lines show Theil–Sen slopes for 50 randomly selected stable (𝛿 > 0, LB95%, blue) and non-stable (𝛿 < -0.1 or LB50%, orange) performers.

*Hierarchical Bayesian modelling*

We modelled performance on the word recall test using a hierarchical Bayesian model, which lets us describe both how recall typically changes with age in the population, and how individual participants vary around that population pattern. Because recall scores and the latent variables can be skewed, we model them with a skewed normal distribution. We use the term “spread” for the scale of this distribution. When there is no skewness, this spread is the same as the usual standard deviation. An overview of the model is shown in Online Methods Figure 10.

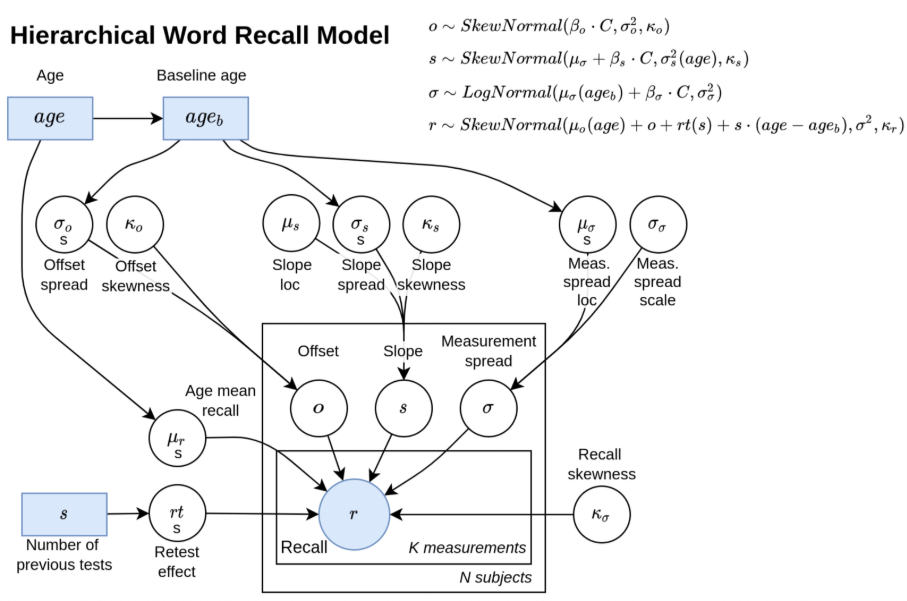

***Online Methods Figure 10. Hierarchical Bayesian model.***

The average recall in the population is modelled as a smooth function of age. For each participant, we estimate three subject-specific parameters: 1) An offset: how their typical recall level differs from the average participant of the same age. 2) A slope: how their recall changes with age, relative to the population trend. 3) A measurement spread: how much their scores vary from session to session. These subject-specific parameters are assumed to come from broader population distributions. This gives the priors for every participant.

The prior of the offsets is modelled with a skewed normal distribution with mean fixed at zero (on average, individuals do not systematically differ from the age-specific mean), spread given by a smooth function of baseline age, and a skewness parameter. The prior of the slopes are modelled with a skewed normal distribution with an estimated mean, spread given by a smooth function of baseline age, and a skewness parameter. The prior of the measurement spreads are modelled with a log-normal distribution, whose location (on the log scale) is given by a smooth function of baseline age, and with an estimated scale parameter. The practice effect is modelled as a smooth function of the number of previous sessions.

Finally, each observed recall score is modelled with a skewed normal distribution. The mean of this distribution combines the population mean for that age, the participant’s offset at baseline, the participant’s slope times the age since baseline, and the practice effect. The spread of this distribution is given by the participant’s own measurement spread, while the skewness is shared across the population.

This framework allows us to describe how recall changes with age at the population level, while also capturing individual differences, practice effects, and potentially skewed distributions of scores. The model parameters are inferred from the data using Markov Chain Monte Carlo and the NUTS kernel giving the posterior distribution of all the parameters given the data.

*Brain relationship analyses*

Longitudinal structural MRI data were available for each participant as a series of cortical and subcortical volumes across two or more time points. For each scan we obtained regional volumes from standard automated segmentation, including bilateral cortical regions and subcortical structures, as well as global measures such as total gray matter and estimated total intracranial volume (eTIV). Scanner/site effects were removed at the regional level by regressing each raw volume on scanner site and using the residuals as site-adjusted volumes.

For each participant and each brain region, we then modeled volume as a function of age using Theil–Sen regression, using all available brain scans. This yielded a robust linear slope (change in volume per year) and an intercept. To anchor slopes to a meaningful level, we computed each person’s “midpoint age” for brain imaging as the median of their observed scan ages and derived the corresponding estimated volume at that age by combining the intercept and slope. Percent change per year was then defined as the annual slope divided by this midpoint volume, multiplied by 100. This converts all regional slopes to a comparable percent-change metric.

To reduce the influence of extreme and likely artifactual slopes, we applied a two-stage trimming procedure to the percent change per year values for each region: first removing the most extreme tails based on a small percentile cut-off, and then excluding remaining values that lay more than a fixed number of standard deviations from the regional mean. For cortical areas measured separately in the left and right hemispheres, we averaged the left- and right-hemisphere percent-change measures to obtain a single bilateral slope.

We focused on the contrast between conservative stable performers and lenient non-stable performers. For each bilateral region of interest, we fitted a separate logistic regression model with conservative stable performers status as the binary outcome (1 = conservative stable performers, 0 = lenient non-stable performers). The main predictor was the regional annual percent change, standardized to have mean zero and unit variance across the analysis sample. All models were adjusted for age (continuous), sex, and eTIV (both also standardized). To account for the fact that slopes based on very short follow-up intervals are much noisier than slopes based on long intervals, we used interval weighting, so that each participant’s contribution to the likelihood was weighted in proportion to their brain follow-up span, after enforcing a small lower bound so that very short intervals did not receive zero weight (11). From each model we extracted the coefficient for the regional brain-change slope and converted it to an odds ratio per one standard deviation more positive slope, along with 95% confidence intervals and Wald p-values. We controlled the false discovery rate across all regions using the Benjamini–Hochberg procedure with a conventional 5% threshold.

*Univariate analyses of stable performers characteristics*

We examined which characteristics differentiate conservative stable performers from lenient non- stable performers using age-adjusted univariate logistic regressions, thereby excluding ambiguous cases. Thirty-seven candidate predictors plus sex spanned early-life factors, education, health and function, psychosocial measures, lifestyle, cognition, and income. For each predictor we took one value per participant from the visit closest to that participant’s analysis age (their within-person median age). Continuous predictors were z-standardized (mean = 0, SD = 1) so odds ratios (ORs) reflect change per +1 SD; strictly binary predictors were coded 0/1. Models used case-wise deletion for the predictor under test, with age and outcome always retained.

Our primary univariate models used inverse-prevalence class weights (0.5/π for each class, where π is the class prevalence) to mitigate outcome imbalance. For numeric/binary predictors we fit logit(maintainer) ~ age + predictor and report the predictor’s Wald p-value, OR, and 95% CI. For multi-level factors we compared an age-only model to an age+factor model via a likelihood-ratio test (overall p-value), with level-specific ORs plotted descriptively. To control multiplicity, Benjamini–Hochberg FDR was applied across all univariate tests (numeric/binary Wald p-values and factor likelihood ratio test p-values join we quantified each participant’s noise level as the within-person residual standard tly), with q < 0.05 considered significant. As robustness checks, we verified that (i) inverse-prevalence weighting vs. unweighted fits produced near-identical effect estimates and discoveries, (ii) restricting to predictors with ≤20% missingness did not change conclusions, and (iii) results were qualitatively unchanged when standardizing continuous predictors within sex strata.

*Multivariate analyses of stable performers characteristics with elastic nets*

To jointly model predictors of stable performers status, we fit elastic-net penalized logistic regression with age included unpenalized (forced into the model). Preprocessing matched the univariate setup. To avoid dropping participants due to scattered missingness, we applied simple single imputation before model fitting (numeric: median; binary/factor: mode), keeping the analysis sample intact. We used inverse-prevalence class weights (weight = 0.5/prevalence for each class) to reduce the impact of class imbalance.

We tuned the mixing parameter over a small grid (α = 0.10, 0.30, 0.50, 0.70) and selected the penalty λ by 10-fold cross-validation using CV-AUC and the “1-SE” rule; if the 1-SE model selected no predictors beyond age, we fell back to λ_min. After selecting (α, λ), we refit a standard (unpenalized) logistic regression on the set of non-zero predictors to obtain interpretable odds ratios with Wald standard errors and 95% CIs; FDR (Benjamini–Hochberg) was used across these multivariate p-values. Discrimination was summarized with ROC curves and AUCs from the elastic-net model and, for comparison, from the refit MLE model) evaluated on the full analysis sample. As a robustness check, we verified that using complete-case rows instead of imputation reduced sample size substantially without changing the main pattern of selected predictors or AUC, and that the MLE refit produced similar apparent AUC to the elastic-net model.

*Individual vs. group trajectory simulation*

We conducted a simulation experiment to examine how apparent age-related decline at the population level can arise from heterogeneous individual trajectories. We simulated 5,000 virtual participants observed annually from ages 30 to 90 on a hypothetical memory test ranging from 0 to 20 points. Each participant began near a mean score of 16 with small between-person variability and a near-flat baseline slope. At random ages, determined by an age-dependent hazard function, individuals experienced one to three short “bursts” of accelerated decline, lasting 2–4 years each, with steeper slopes at older ages. Memory scores were generated from a latent unbounded process mapped through a logistic function to constrain values to the 0–20 range. We added small measurement noise to mimic test variability. The simulated trajectories were then aggregated and modelled with a generalized additive model (GAM) to estimate the group-level age curve.

*Burst-decline analyses*

Data & preprocessing: We used the practice-corrected recall score across visits and used the Theil-Sen slope and a robust intercept as the median of estimated scores anchored at the median age of each participant. We then computed residuals (rij=yij−(b^0i+s^iaij) so that long-run age trends (including practice adjustment) are removed. Consecutive visits define intervals between visits with duration Δt=a_i,j+1_−a_ij_>0, raw change Δy=y_i,j+1_−y_ij_, residual change Δr=r_i,j+1_−r_ij_, and segment slope Δy/Δt.

Burst definitions: An interval is flagged as a burst if it is (i) fast (segment slope ≤ −0.50 words/year), (ii) substantial (Δy≤−1.50 words), and (iii) short (Δt≤3 years). For “big” bursts we additionally require z-scaled magnitude – Δy / SD_i_(r) ≥z_thr_, where SD_i_(r) is that person’s residual SD.

Recovery definition: For each burst with onset at time *t_on_* we scan future intervals at least ≥ the minimum gap after onset (gap ∈ (12) years) and take the best future residual value. A burst is considered recovered if

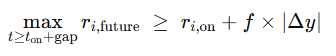

i.e., if the participant rebounds by at least a fraction *f* toward their pre-drop level, with **f** ∈ {0.40,0.50}. Otherwise, it is non-recovered. (Here *r_i,on_* is the residual at the burst’s onset interval.)

Permutation null (noise-only timing): To test whether the observed rates could arise from random timing of within-person fluctuations, we generated a permutation null that shuffles residuals within each person (preserving each ID’s residual distribution, number and spacing of visits, and fitted trajectory *^b_0i_+^s_i_a*). For each shuffle we reconstructed the time series y*_ij_ = fitted_ij_ + r(perm)_ij_, rebuilt intervals, re-applied the burst and recovery rules, and recomputed both outcomes. We ran 200 permutations.

*Stable performers state analyses in HRS - Maintainer bout detection and back projection analyses*

Bout detection and “time-limited state” summaries: For each participant, we searched all contiguous windows of length 4 up to the person’s total number of observations, starting with the longest possible window, and selected the first window that satisfied the maintainer criterion (thereby identifying the longest qualifying maintainer period). For the selected bout we recorded: bout length (waves) and duration (years; end age − start age), start age and end age, the LOO lower-bound statistic (LB_0.05)

To characterize stability as a time-limited state, we estimated robust pre- and post-bout slopes using Theil–Sen on observations outside the bout:

Pre-bout slope: TS slope of observations before the bout (requires ≥2 observations)

Post-bout slope: TS slope of observations after the bout (requires ≥2 observations)

We then summarized: (i) prevalence of having ≥1 qualifying bout (≥4 waves), (ii) mean duration of the longest bout among bout-positive participants, and (iii) the fraction and mean magnitude of decline (slope < 0) in pre- and post-bout segments, among participants with estimable pre/post segments.

Age-window definition (75–85 overlap): For age-specific prevalence in later life, we defined a bout as relevant to ages 75–85 if it overlapped the 75-85 year age window. For each participant, we computed the longest qualifying bout that overlapped ages 75–85 (if any). The denominator for 75–85 analyses comprised participants with at least one observed assessment within the 75–85 interval.

Because strict stability criteria can still yield false positives under noise and sparse sampling, we estimated an empirical null distribution using simulation calibrated to the observed data structure, following the same logic as for SHARE and BETULA. For each simulated dataset, we re-ran the identical bout-detection algorithm (overall and 75–85 overlap), yielding null prevalences for thresholds ≥4 to ≥13 waves.

PPV / Se-corrected “true prevalence”: Let Obs denote the observed prevalence of meeting a bout-length threshold (e.g., having a ≥10-wave maintainer bout), and Null denote the corresponding simulated prevalence under the null model. Treating Null as the false-positive rate and assuming sensitivity Se for detecting true stability periods, we estimated the underlying true prevalence:

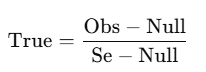

We set Se = 0.80 (primary analysis). Uncertainty bands were obtained by summarizing the null distribution across simulations (median and 5th/95th percentiles) and propagating these into conservative bounds for True by pairing Obs with the null high/low values.

All analyses were implemented in R using data.table for efficient data manipulation, mgcv for GAM fitting, and parallel for simulation. Random number streams were controlled via fixed seeds for reproducibility.

*Back-projection analyses in HRS: testing maintenance as trait versus state*

We used episodic memory in the Health and Retirement Study (HRS) after practice adjustment, analyzed on the age scale. All analyses were restricted to the simulation-aligned HRS analytic sample used throughout the manuscript. The long-format dataset contained participant identifier, age at each assessment (years), and practice-adjusted memory score. Birth year was obtained from the RAND HRS file (rabyear) and merged by participant identifier (hhidpn).

All rates of change were estimated using Theil–Sen (TS) regression. For each participant, we first estimated an overall TS slope using all available observations, representing the average rate of change across the full observation window. We paired this slope with a robust intercept defined consistently with TS estimation by centering the fitted line on the median of the participant’s observed scores relative to age.

To operationalize stability as a time-limited state, we then searched for “stable performance bouts.” For each participant we examined all contiguous sequences of assessments with a minimum length of four waves and evaluated each candidate segment using a conservative leave-one-out procedure. Specifically, within a candidate segment we repeatedly re-estimated the TS slope while omitting one observation at a time and summarized these leave-one-out slopes by a lower-tail value. A segment was classified as stable if this conservative lower bound was not negative. For each participant, we retained the longest segment meeting this stability requirement as the participant’s longest stable bout and recorded its duration in waves, age span, midpoint age, and the TS slope estimated within the bout.

A central design choice in back-projection is how to anchor the linear trajectory used for extrapolation to midlife. We therefore implemented two anchoring strategies that map directly onto the trait versus state interpretations of maintenance. In the within-bout back-projection (the primary state-based test), bout maintainers were projected using the TS slope and robust intercept estimated from observations inside their stable bout, so that the extrapolated trajectory is anchored specifically to the plateau-like segment. In contrast, non-stable performers were projected using their overall TS slope and robust intercept estimated from all observations, reflecting their typical change across the observed period. In the whole-window back-projection (a complementary trait-like projection), bout stable performers were also projected using their overall TS slope and robust intercept computed from the full observation window, matching the approach used for non-stable performers; this makes the stability projection reflect the participant’s full observed trajectory rather than being anchored to the stable segment.

For each participant we generated a predicted memory score at age 55 by extrapolating the chosen TS line to that age (within-bout line for the within-bout analysis; overall line for the whole-window analysis). To interpret back-projected scores relative to same-cohort peers, we standardized predicted age-55 values within birth cohort. Birth cohorts were defined in decade bins (e.g., 1930s). Cohort norms were estimated empirically from participants with midlife observations (ages 50–60). For each participant contributing to the norms, we estimated a TS slope using their observations in the 50–60 age range and predicted their score at age 55; the cohort-specific mean and standard deviation of these predicted age-55 values were then used to standardize all back-projected age-55 scores. This procedure ensures that cohort standardization is anchored to the midlife distribution rather than being influenced by late-life performance.

Back-projection analyses were carried out among older participants, defined as those whose maximum observed age was at least 75 years. Within this subset, bout stable performers were defined as participants with at least one qualifying stable bout of length four waves or longer, while non-maintainers were those meeting the older criterion but without any qualifying stable bout.

We quantified the “back-projection paradox” by evaluating how often back-projected standardized age-55 values fell below cohort-relative thresholds corresponding to the bottom quartile and progressively more extreme cutoffs (bottom 15%, bottom 10%, and bottom 5%). For each threshold we computed the proportion below the cutoff separately for bout stable performers and non-stable performers, and summarized group differences using risk ratios and risk differences.

Conceptually, the within-bout back-projection is designed as a direct test of whether late-life stability reflects a trait-like near-flat trajectory that plausibly extends back across adulthood. If stability were lifelong resistance, then trajectories anchored to the stable segment should not imply implausibly low midlife performance. By contrast, if stability is typically a time-limited state embedded within a broader non-monotonic trajectory, then anchoring projections to the plateau itself can yield paradoxical reconstructions at age 55. The whole-window back-projection provides a complementary comparison by anchoring stable performers to their full observed trajectory; under a state interpretation this should attenuate the paradox because the overall line incorporates surrounding decline episodes rather than reflecting only the stable segment.

**Online Methods References**

1.	A. Borsch-Supan *et al.*, Data Resource Profile: the Survey of Health, Ageing and Retirement in Europe (SHARE). *Int J Epidemiol* **42**, 992–1001 (2013).

2.	S. Gruber, C. Hunkler, S. Stuck (2014) Generating easySHARE: guidelines, structure, content and programming. in *SHARE Working Paper Series* (Munich: MEA, Max Planck Institute for Social Law and Social Policy).

3.	M. Bergmann, T. Kneip, G. De Luca, A. Scherpenzeel (2019) Survey participation in the Survey of Health, Ageing and Retirement in Europe (SHARE), Wave 1-7. Based on Release 7.0.0. . in *SHARE Working Paper Series* (Munich: MEA, Max Planck Institute for Social Law and Social Policy).

4.	A. Börsch-Supan, S. Gruber, easySHARE. Release version: 8.0.0. SHARE-ERIC. 10.6103/SHARE.easy.800 (2022).

5.	T. Mehrbrodt, S. Gruber, M. Wagner (2019) Scales and Multi-Item Indicators in the Survey of Health, Ageing and Retirement in Europe. in *SHARE Working Paper Series* (Germany).
