## Supplemental Information for "Punctuated memory change: The temporal dynamics and brain basis of memory stability in aging"

| Section | Title | Page |
| --- | --- | --- |
| 1 | Memory change over age in SHARE | 2 |
|  | Supplementary Figure 1 | 2 |
|  | Supplementary Figure 2 | 3 |
| 2 | Distinguishing “true stable performers” from chance | 4 |
|  | Supplementary Figure 3 | 4 |
| 3 | Frequentialist vs. Bayesian modelling | 5 |
|  | Supplementary Figure 4 | 5 |
|  | Supplementary Figure 5 | 5 |
| 4 | Example plots of individual stable performers | 6 |
|  | Supplementary Figure 6 | 6 |
|  | Supplementary Figure 7 | 7 |
|  | Supplementary Figure 8 | 8 |
|  | Supplementary Figure 9 | 9 |
| 5 | Memory analyses in the MRI subsample | 10 |
|  | Supplementary Figure 10 | 10 |
|  | Supplementary Figure 11 | 10 |
| 6 | Brain atrophy analyses | 11 |
|  | Supplementary Table 1 | 11 |
|  | Supplementary Figure 12 | 13 |
|  | Supplementary Figure 13 | 14 |
| 7 | Robustness checks for analyses of what characterizes stable performers | 15 |
| 8 | Robustness checks for mortality analyses | 16 |
| 9 | Identification of burst-decline in individual SHARE participants | 18 |
|  | Supplementary Figure 14 | 19 |
| 10 | Offset – slope analyses | 20 |
|  | Supplementary Figure 15 | 20 |
|  | Supplementary Figure 16 | 20 |
|  | Supplementary Figure 17 | 21 |
| 11 | Maintenance bout analyses – stability as a state | 22 |
|  | Supplementary Figure 18 | 22 |
|  | Supplementary Figure 19 | 23 |
| 12 | Additional information on MRI subsample datasets | 24 |
|  | Supplementary Table 2 | 24 |
| 13 | Funding, acknowledgements and additional cohort information | 29 |
| 14 | Additional SHARE and Betula analyses | 32 |
|  | Supplementary Figure 20 | 32 |
|  | Supplementary Figure 21 | 33 |
| 15 | Consortia authors | 36 |

**Section 1: Memory change over age**

The overall median slope was -0.197 words/year (95% CI for the population median: -0.200, -0.192. To summarize age trends (Figure 1), we plotted a longitudinal curve obtained by smoothing the per-participant Theil–Sen slopes as a function of each participant’s median age using a GAM (F = 212.2, p < 2×10⁻¹⁶). This was compared to a cross-sectional curve computed as the first derivative of a GAM fitted to recall versus age using all visits (F = 12,231.1, p < 2×10⁻¹⁶). Over most of the age span (≈60–80 years), the curves were approximately parallel, but the longitudinal decline was consistently steeper with narrower CIs. Restricting the cross-sectional model to the same ≥4-visit subsample used for the longitudinal analyses produced identical shapes (SI Figure X). The discrepancy is expected under several well-documented mechanisms. First, selective survival and participation enrich older age groups with healthier, higher-functioning individuals, inflating means at later ages and attenuating cross-sectional age contrasts. Second, cohort differences can bias cross-sectional comparisons toward smaller declines. Third, the longitudinal estimator is a secant (interval-average) while the cross-sectional derivative is a tangent (instantaneous rate). With accelerating decline, interval-averaged slopes are expected to be more negative.

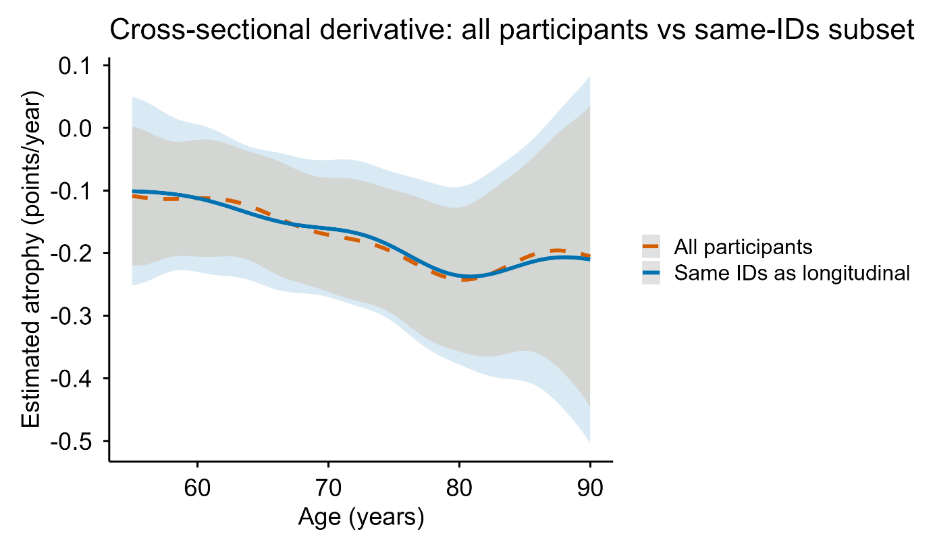

**Supplementary Figure 1 Estimated memory decline from cross-sectional age-trends.** A cross-sectional curve computed as the first derivative of a GAM fitted to recall versus age using all visits was used as an approximation of age-decline. The orange line shows the results for the full SHARE sample (n = 158,291) and the blue line represents the results for the longitudinal subsample with > 4 visits (n = 55,318). Shaded ribbons denote 95% CI.

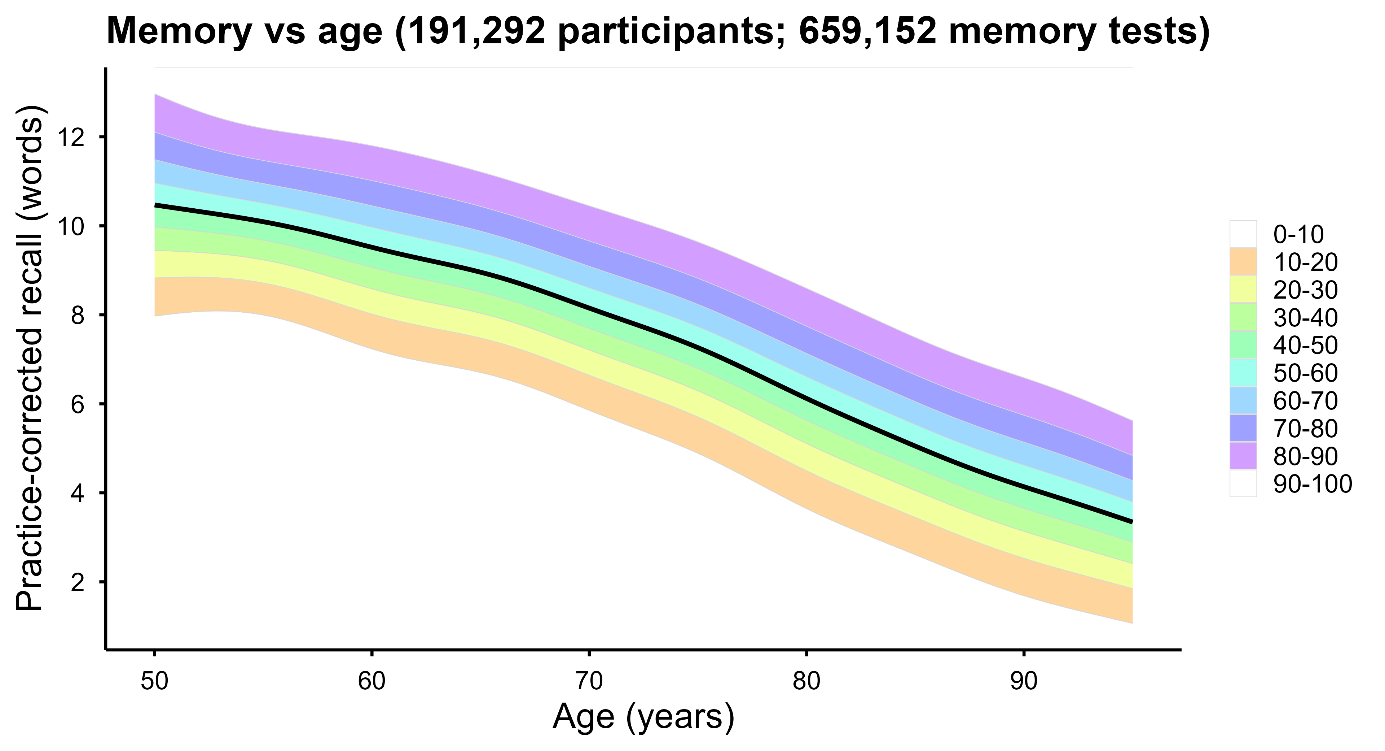

**Supplementary Figure 2. Pooled age trend in practice-corrected episodic memory in SHARE and HRS with model-based predictive bands.** Practice-corrected recall (words; immediate + delayed 10-word recall) is plotted against age (50–95 years) using pooled data from SHARE and HRS. The black curve shows the estimated mean trajectory from a generalized additive model (GAM) fitted to all observations with a penalized spline for age (mgcv::bam; smoothing basis with k=12). To avoid over-weighting participants with many repeated assessments, observations were weighted by the inverse of each participant’s number of available visits (each participant contributing total weight ≈1). Shaded bands represent model-based predictive dispersion around the mean trajectory. The shaded colors show the predictive distribution into 10 bands (0–10%, …, 90–100%), with extreme bands optionally de-emphasized for visualization. Band width is allowed to vary with age by fitting a second GAM to the log of squared residuals as a smooth function of age and converting predictions to an age-specific residual SD; predictive bands are then computed as the mean curve plus quantile-specific multipliers of this SD at each age.

**Section 2: Distinguishing “true stable performers” from chance**

To separate real stable performers from residual noise, we ran a simulation experiment. We generated a dataset where all participants followed the estimated age-expected memory decline, and added each participant’s own within-person noise, estimated as deviations from the participant’s linear slope based on the real data. This allowed us to estimate how often our rules would flag stable performers if stable performers did not actually exist.

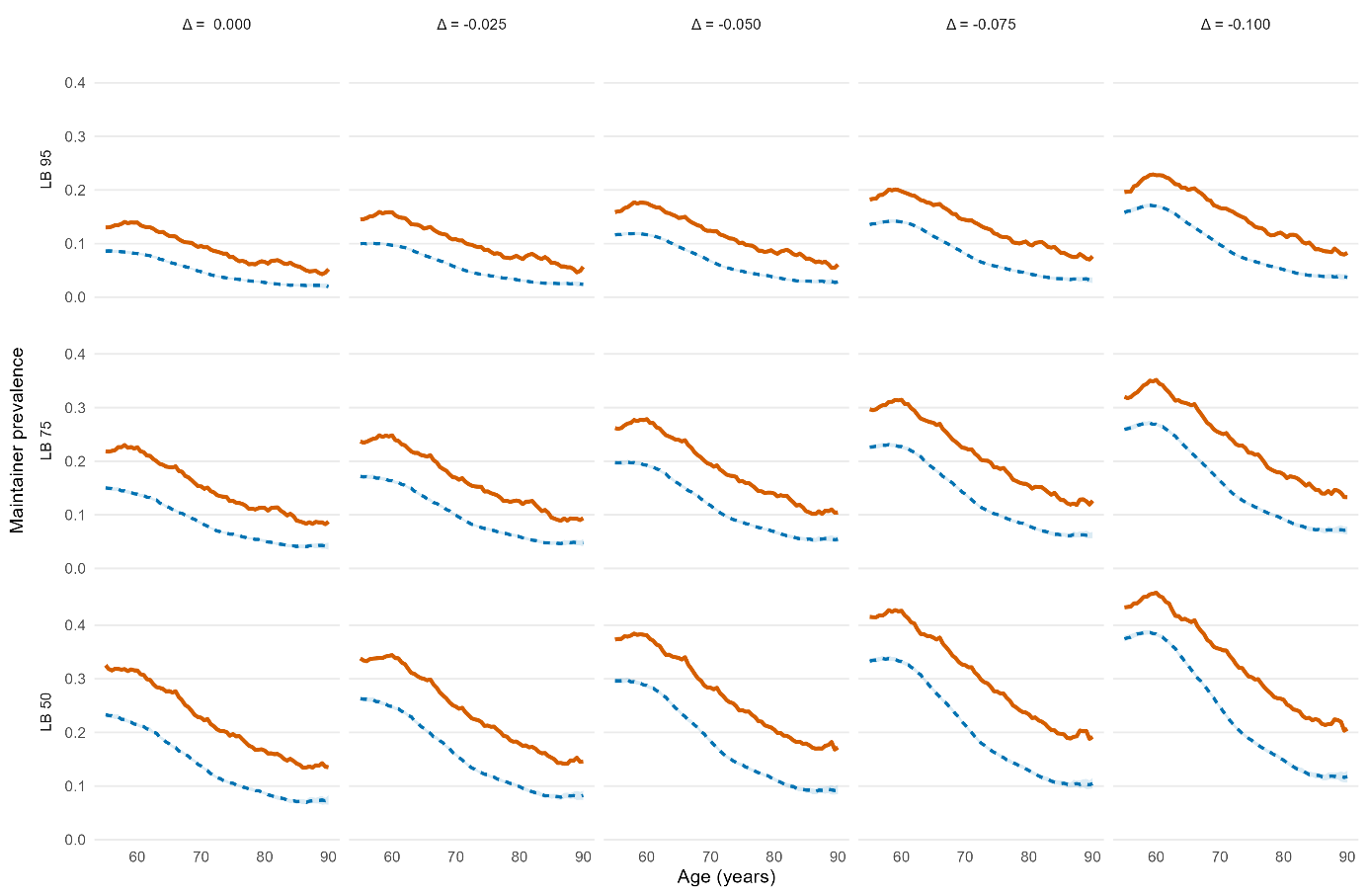

**Supplementary Figure 3 Simulation results across thresholds.** We compared the observed stable performers prevalence (orange solid lines) to the null (chance-only) prevalence for a grid of definitions that vary the slope threshold and the confidence requirement (LB threshold) (blue dotted lines). Plotting the observed prevalence against the simulation median and interquartile range (IQR) showed that for all the 15 tested rules (5 slope × 3 LB variations), the observed curve exceeds the null IQR.

**Section 3: Frequentialist vs. Bayesian modelling**

| Frequentialist framework | Bayesian framework |
| --- | --- |
| 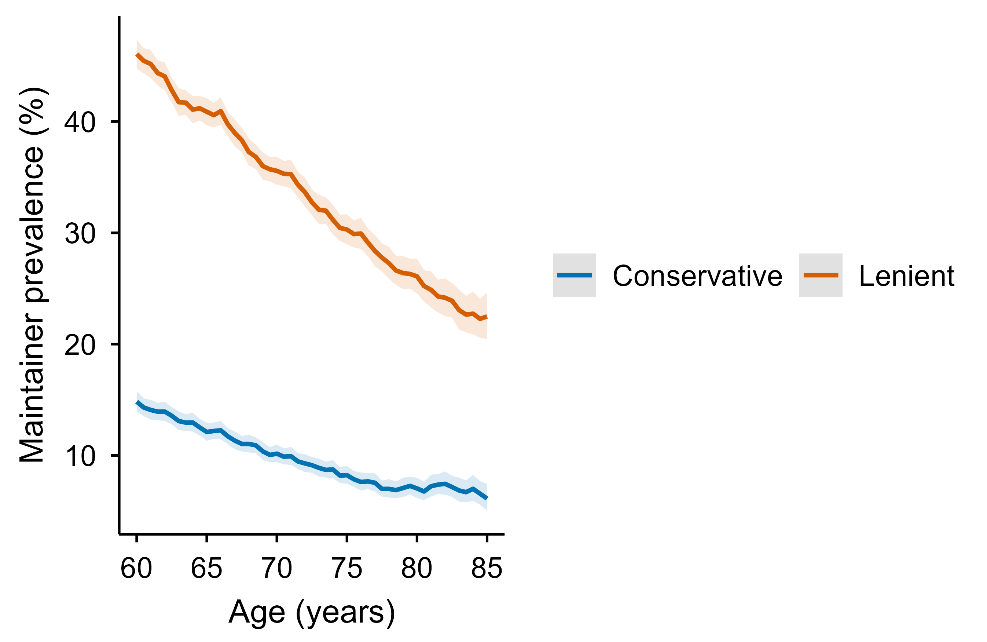 | 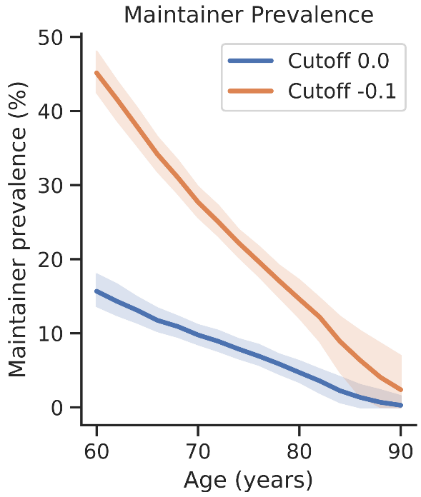 |

**Supplementary Figure 4. Prevalence of observed stable performers across age using robust Theil-Sen regression and hierarchical Bayesian modelling in SHARE.** Left panel: The lines show the observed percentages of stable performers within a ±1.5-year moving window using the Thiel-Sen approach. Shaded bands are 95% Wald CIs. The conservative rule (blue) has 95% LOO lower bound (LB) and slope ≥ 0, whereas the lenient rule (orange) has 50% LOO LB and slope ≥ -0.1. Right panel: Stable performers prevalence with age in the hierarchical Bayesian modelling framework. Shaded bands are 95%.

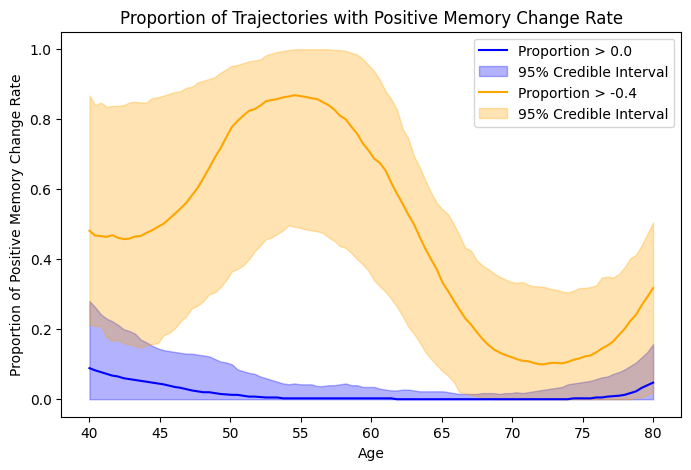

**Supplementary Figure 5.** Stable performers prevalence as a function of age in Betula under the hierarchical Bayesian framework according to a conservative (> 0) and lenient (> -0.4) slope criterion.

**Section 4: Example plots of individual stable performers**

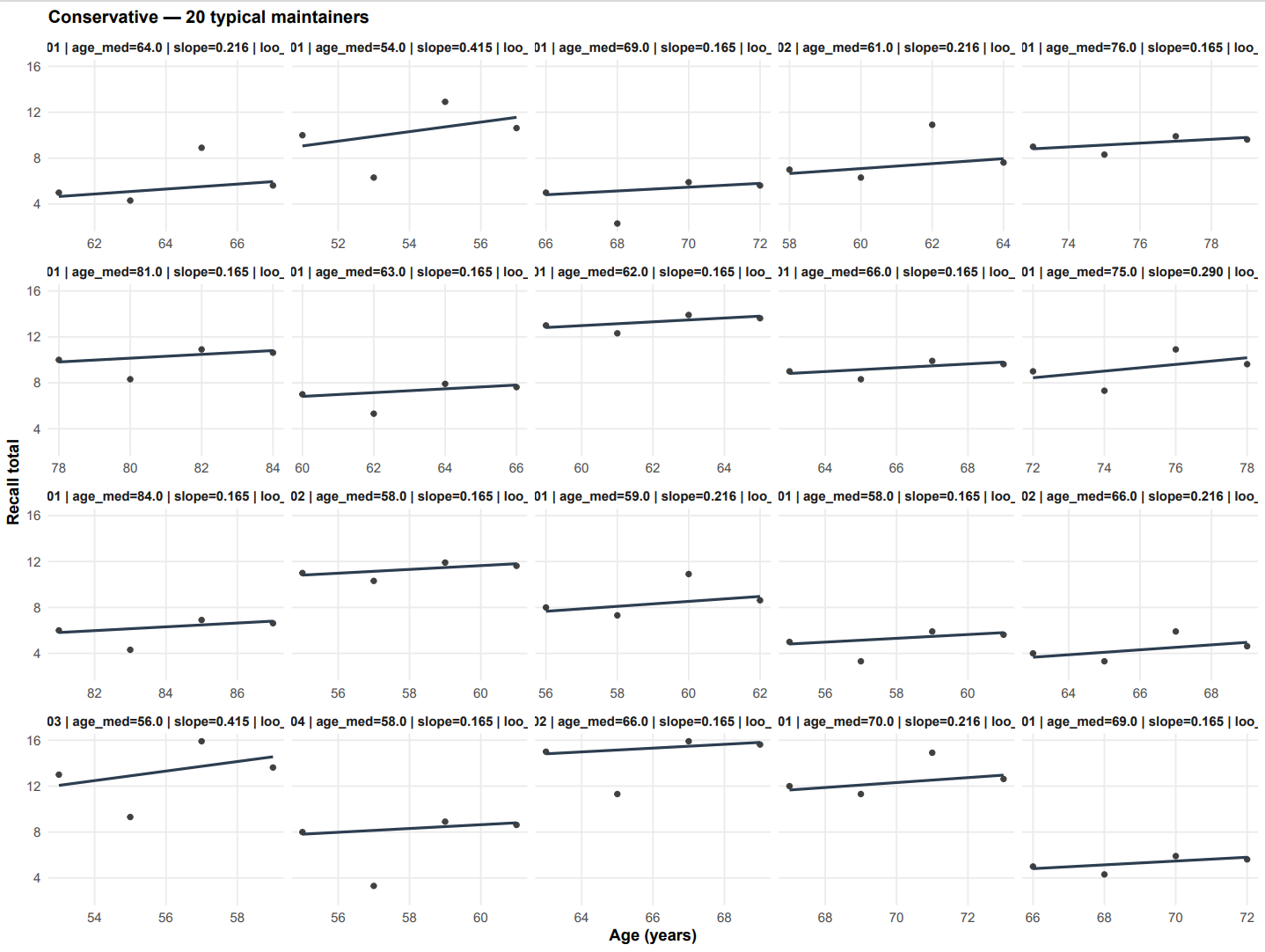

**Supplementary Figure 6. Examples of stable performers according to the conservative rule (slope ≥ 0, LB 95%).**

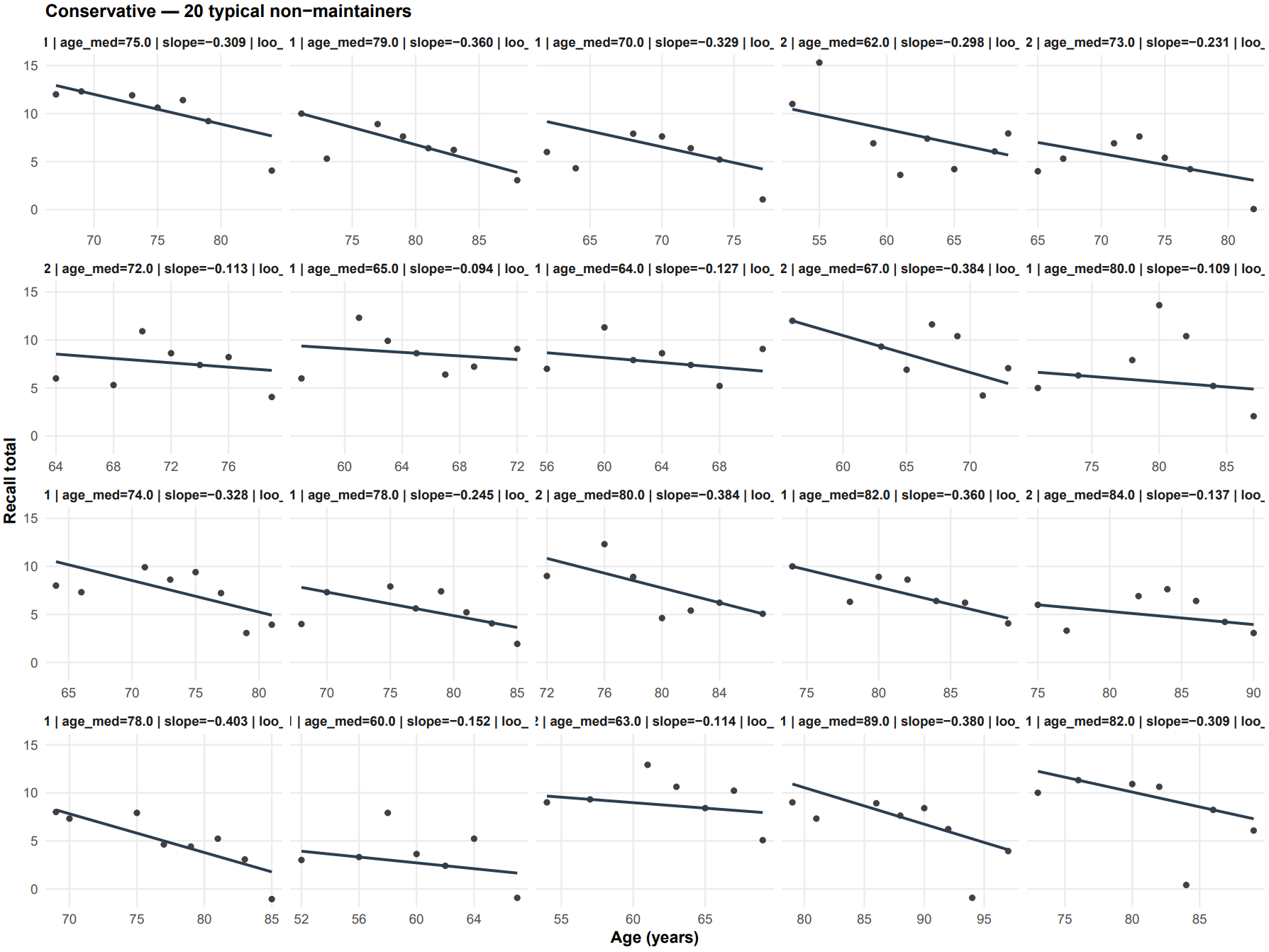

**Supplementary Figure 7. Examples of non-** **stable performers according to the conservative rule (slope ≥ 0, LB 95%).**

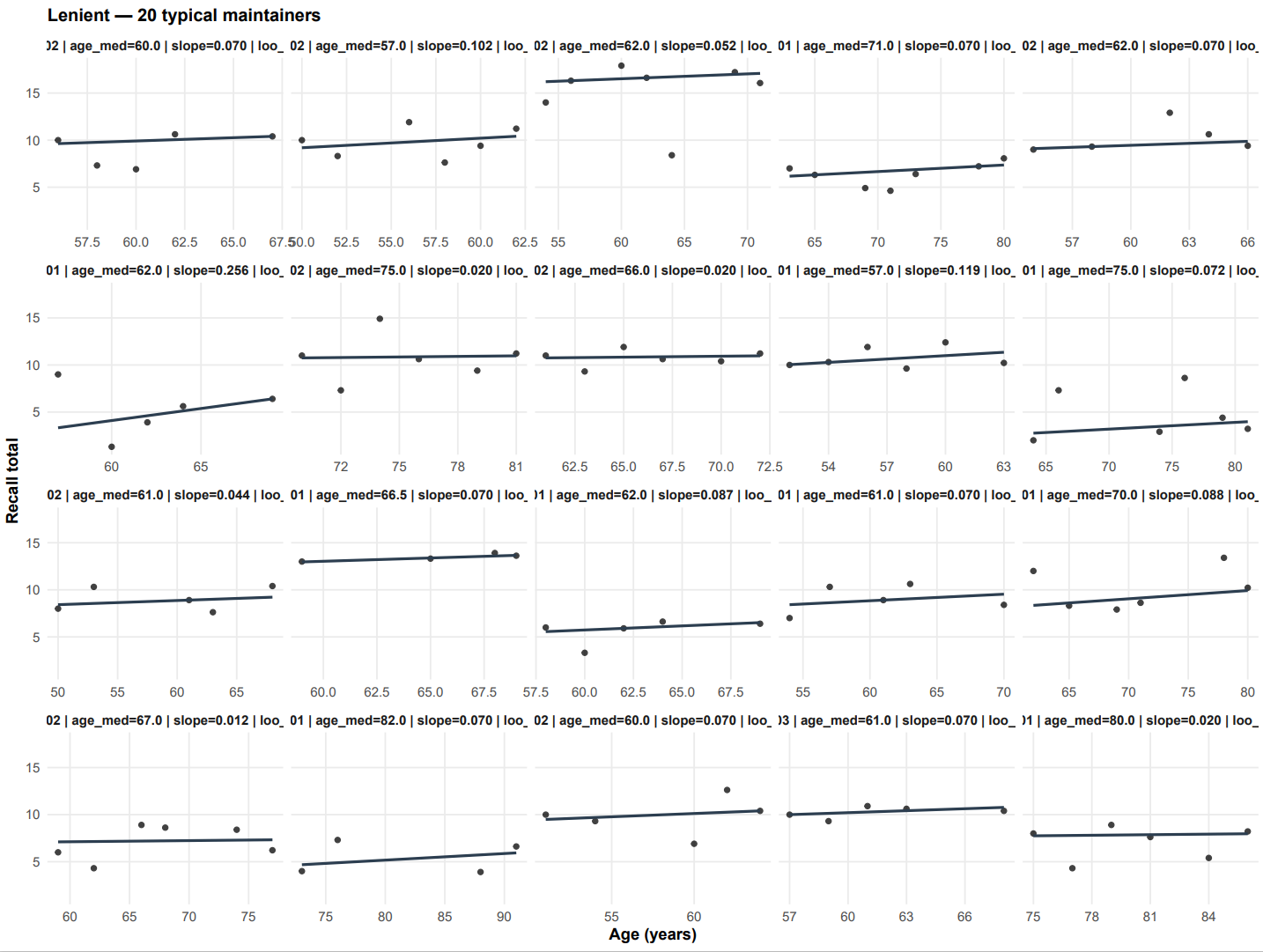

**Supplementary Figure 8. Examples of stable performers according to the lenient rule (slope ≥ -0.1, LB 50%).**

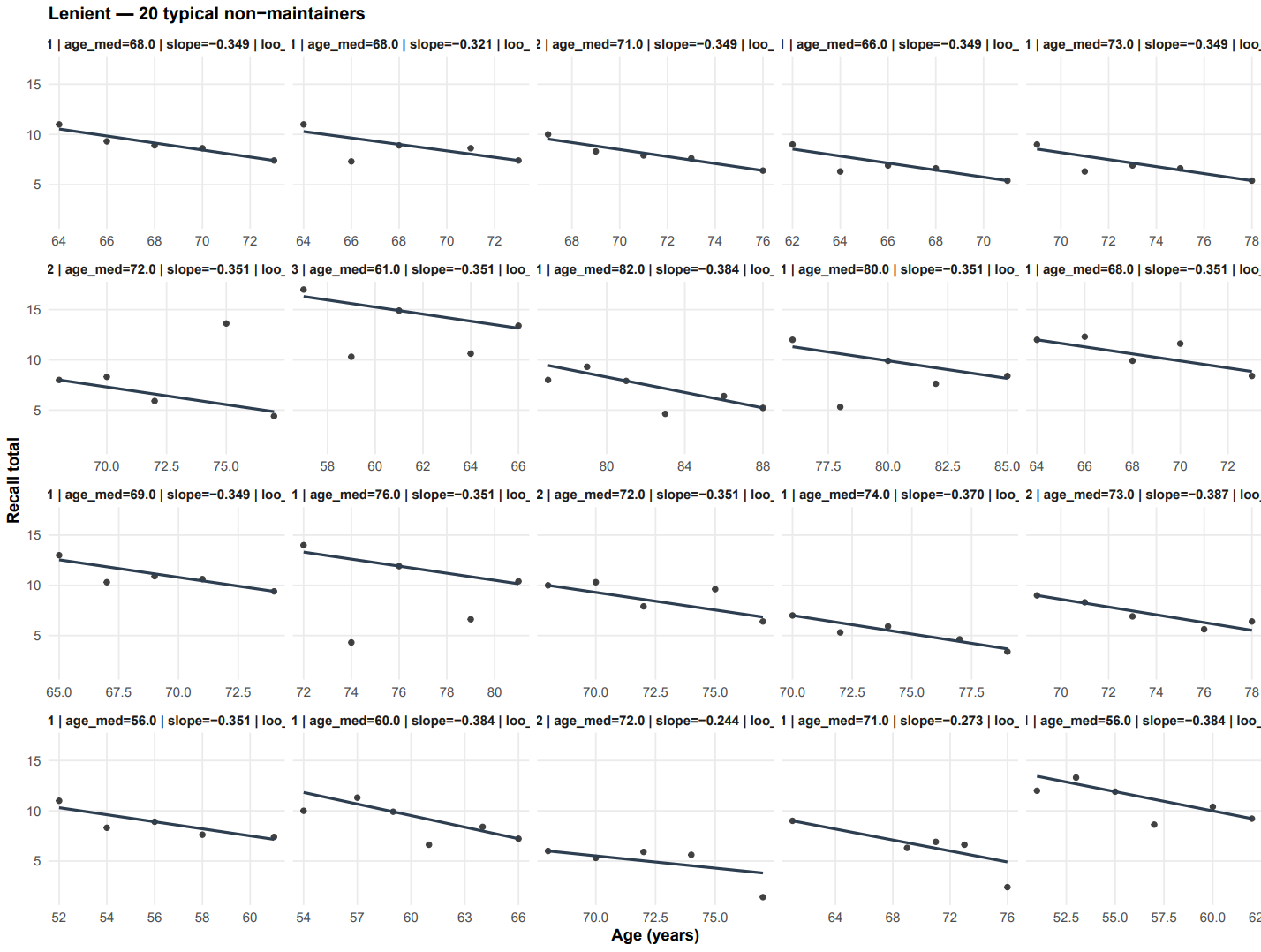

**Supplementary Figure 9. Examples of non-** **stable performers according to the lenient rule (slope ≥ -0.1, LB 50%).**

**Section 5: Memory analyses in the MRI subsample**

We analyzed longitudinal MRI data from 10 independent studies (n =1,978 participants aged > 50 with > 4 episodic-memory test sessions covering 16,342 tests and 7,206 scans). We extracted a principal component across memory subtests within each study and classified stable performers using the same Theil–Sen–based framework. The PPV-estimated proportion of true stable performers was 9.8% (95% CI: 9.3–10.4) for ages ≥70 (95% CI: 8.2–9.2) for ages ≥80, slightly higher than in SHARE and BETULA, likely reflecting more selectivity and selection bias in MRI cohorts.

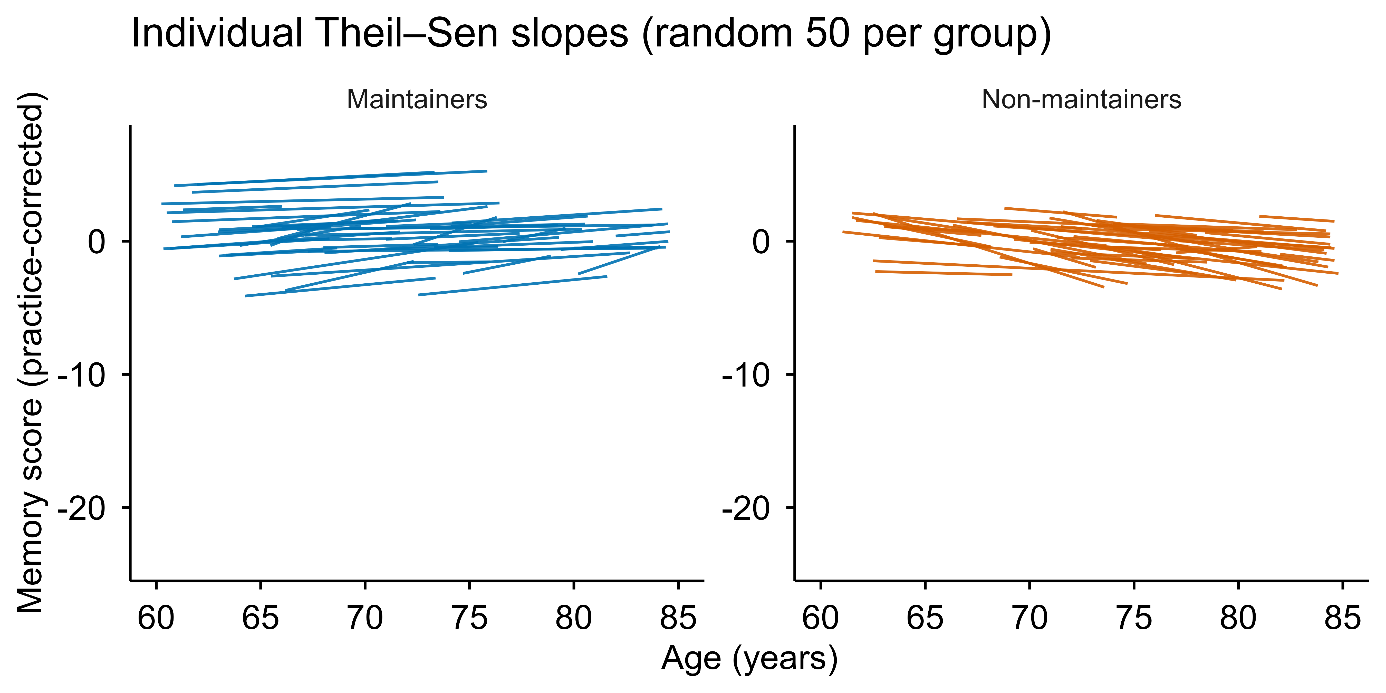

**Supplementary Figure 10. Individual memory trajectories in the MRI subsample.** Lines show Theil–Sen slopes for 50 randomly selected stable performers (𝛿 > 0, LB > 95, blue) and non- stable performers (𝛿 < -0.1 or LB < 50, orange).

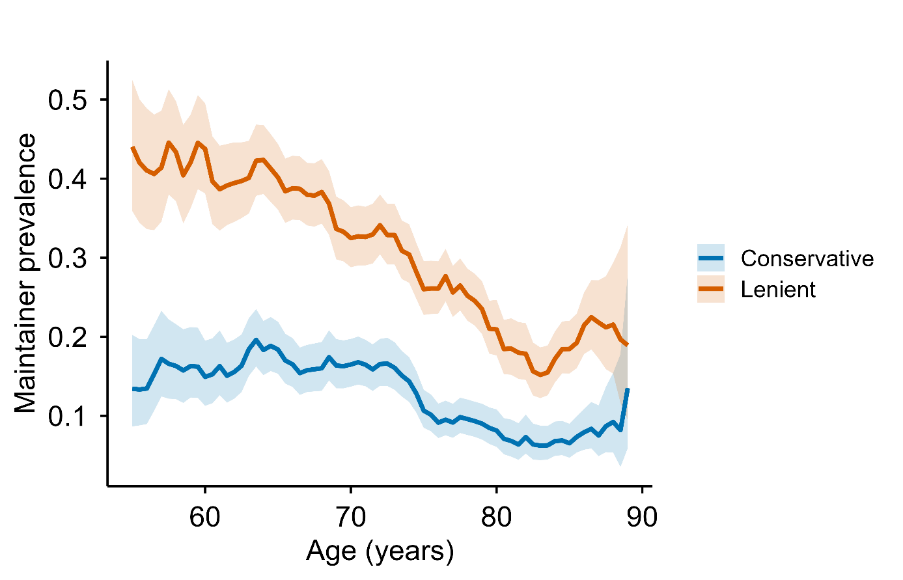

**Supplementary Figure 11. Prevalence of observed stable performers across age using robust Theil-Sen regression in the MRI subsample.** The lines show the observed percentages of stable performers within a ±1.5-year moving window using the Thiel-Sen approach. Shaded bands are 95% Wald CIs. The conservative rule (blue) has 95% LOO lower bound (LB) and slope > 0, whereas the lenient rule (orange) has 50% LOO LB and slope > -0.1.

**Section 6: Brain atrophy analyses**

We used the same Theil–Sen framework to 44 volumetric brain regions, and fitted interval-weighted logistic regressions comparing conservative stable performers (n = 215; mean age 68.5 years [SD 7.5, range 54.0-88.9], 124 women/ 91 men; mean memory follow-up 9.3 years [SD 5.4, range 2.0-21.4]; mean MRI follow-up 6.1 years [SD 4.6, range 1.0-17.4]) to lenient non- stable performers (n = 1,387; mean age 72.4 years [SD 8.5, range 51.6-91.2], 711 women/ 676 men; mean memory follow-up 8.1 years [SD 4.2, range 2.0-22.3]; mean MRI follow-up 5.0 years [SD 3.3, range 1.0-18.1]). Models were adjusted for age at the midpoint of brain follow-up, sex, and intracranial volume. Participants with longer MRI follow-up contributed more strongly to the estimation via weights proportional to their brain follow-up interval (weight = participant’s follow-up span in years / sample mean follow-up span), thereby downweighing noisy slopes from short intervals. FDR was controlled across regions by the Benjamini–Hochberg procedure.

Conservative stable performers showed less atrophy and ventricular expansion than lenient non- stable performers in 27 of 44 regions from the Desikan-Killany cortical atlas and the FreeSurfer ASEG segmentation (Supplementary Information Table 1, Supplementary Figure 12). The pattern was robust to analyses without covariates and to alternative stable performers definitions. As a comparison, the same analyses were run using robust volume levels, with no regions surviving FDR-correction (Supplementary Information Figure 13).

| Brain region | beta | or | lcl | ucl | p | q bh |
| --- | --- | --- | --- | --- | --- | --- |
| Cerebellum-Cortex | 0.29 | 1.34 | 1.20 | 1.49 | 1.29e-07 | 2.84e-06 |
| Hippocampus | 0.35 | 1.42 | 1.23 | 1.63 | 1.07e-06 | 9.44e-06 |
| Inf-Lat-Vent | -0.23 | 0.79 | 0.70 | 0.89 | 0.00 | 0.00 |
| Cerebellum-White-Matter | 0.19 | 1.20 | 1.08 | 1.34 | 0.00 | 0.00 |
| Amygdala | 0.20 | 1.22 | 1.07 | 1.39 | 0.00 | 0.01 |
| Thalamus-Proper | 0.14 | 1.15 | 1.01 | 1.30 | 0.03 | 0.05 |
| Lateral-Ventricle | -0.07 | 0.93 | 0.83 | 1.05 | 0.24 | 0.31 |
| Pallidum | 0.06 | 1.06 | 0.94 | 1.20 | 0.35 | 0.45 |
| Putamen | -0.02 | 0.98 | 0.87 | 1.10 | 0.69 | 0.76 |
| Caudate | -0.02 | 0.98 | 0.86 | 1.11 | 0.73 | 0.78 |
| cuneus | 0.40 | 1.49 | 1.34 | 1.65 | 3.52e-14 | 1.56e-12 |
| isthmuscingulate | 0.29 | 1.33 | 1.19 | 1.48 | 3.78e-07 | 5.54e-06 |
| precuneus | 0.25 | 1.29 | 1.16 | 1.42 | 7.22e-07 | 7.94e-06 |
| pericalcarine | 0.25 | 1.28 | 1.15 | 1.42 | 3.55e-06 | 2.39e-05 |
| superiorparietal | 0.27 | 1.31 | 1.17 | 1.46 | 3.80e-06 | 2.39e-05 |
| caudalmiddlefrontal | 0.23 | 1.26 | 1.14 | 1.41 | 1.81e-05 | 9.98e-05 |
| inferiorparietal | 0.21 | 1.23 | 1.12 | 1.36 | 3.10e-05 | 0.00 |
| parahippocampal | 0.21 | 1.23 | 1.11 | 1.36 | 4.81e-05 | 0.00 |
| precentral | 0.24 | 1.28 | 1.13 | 1.45 | 0.00 | 0.00 |
| supramarginal | 0.18 | 1.20 | 1.09 | 1.32 | 0.00 | 0.00 |
| temporalpole | 0.19 | 1.21 | 1.09 | 1.35 | 0.00 | 0.00 |
| lateraloccipital | 0.17 | 1.18 | 1.07 | 1.30 | 0.00 | 0.00 |
| insula | 0.16 | 1.17 | 1.07 | 1.28 | 0.00 | 0.00 |
| paracentral | 0.18 | 1.20 | 1.06 | 1.35 | 0.00 | 0.01 |
| superiorfrontal | 0.14 | 1.15 | 1.05 | 1.25 | 0.00 | 0.01 |
| inferiortemporal | 0.12 | 1.13 | 1.04 | 1.23 | 0.01 | 0.01 |
| superiortemporal | 0.12 | 1.13 | 1.03 | 1.23 | 0.01 | 0.01 |
| postcentral | 0.14 | 1.15 | 1.04 | 1.27 | 0.01 | 0.02 |
| parsopercularis | 0.12 | 1.13 | 1.03 | 1.24 | 0.01 | 0.02 |
| parstriangularis | 0.11 | 1.12 | 1.02 | 1.23 | 0.02 | 0.04 |
| bankssts | 0.13 | 1.14 | 1.02 | 1.28 | 0.02 | 0.04 |
| transversetemporal | 0.10 | 1.11 | 1.00 | 1.23 | 0.06 | 0.10 |
| posteriorcingulate | 0.09 | 1.10 | 0.99 | 1.21 | 0.08 | 0.11 |
| fusiform | 0.08 | 1.08 | 0.99 | 1.18 | 0.09 | 0.13 |
| rostralmiddlefrontal | 0.07 | 1.07 | 0.97 | 1.18 | 0.15 | 0.21 |
| lingual | 0.07 | 1.07 | 0.98 | 1.17 | 0.15 | 0.21 |
| medialorbitofrontal | -0.04 | 0.96 | 0.88 | 1.05 | 0.36 | 0.45 |
| lateralorbitofrontal | 0.04 | 1.04 | 0.95 | 1.13 | 0.41 | 0.50 |
| middletemporal | 0.04 | 1.04 | 0.95 | 1.14 | 0.43 | 0.51 |
| rostralanteriorcingulate | 0.03 | 1.03 | 0.94 | 1.13 | 0.51 | 0.59 |
| frontalpole | -0.02 | 0.98 | 0.88 | 1.09 | 0.67 | 0.76 |
| parsorbitalis | 0.01 | 1.01 | 0.92 | 1.10 | 0.91 | 0.95 |
| entorhinal | -0.01 | 0.99 | 0.89 | 1.11 | 0.92 | 0.95 |
| caudalanteriorcingulate | 0.00 | 1.00 | 0.91 | 1.11 | 0.95 | 0.95 |

**Supplementary Table 1. Exact statistics for volumetric analyses.** For each bilateral region of interest, a separate logistic regression model with conservative stable performers status as the binary outcome (1 = conservative stable performers, 0 = lenient non- stable performers) was fitted. The main predictor was the regional annual percent change, and all models were adjusted for age, sex, and eTIV (all standardized). We used interval weighting, so each participant’s contribution to the likelihood was weighted in proportion to their brain follow-up span, after enforcing a small lower bound so that very short intervals did not receive zero weight. False discovery rate across all regions was controlled using the Benjamini–Hochberg procedure with a conventional 5% threshold (q bh).

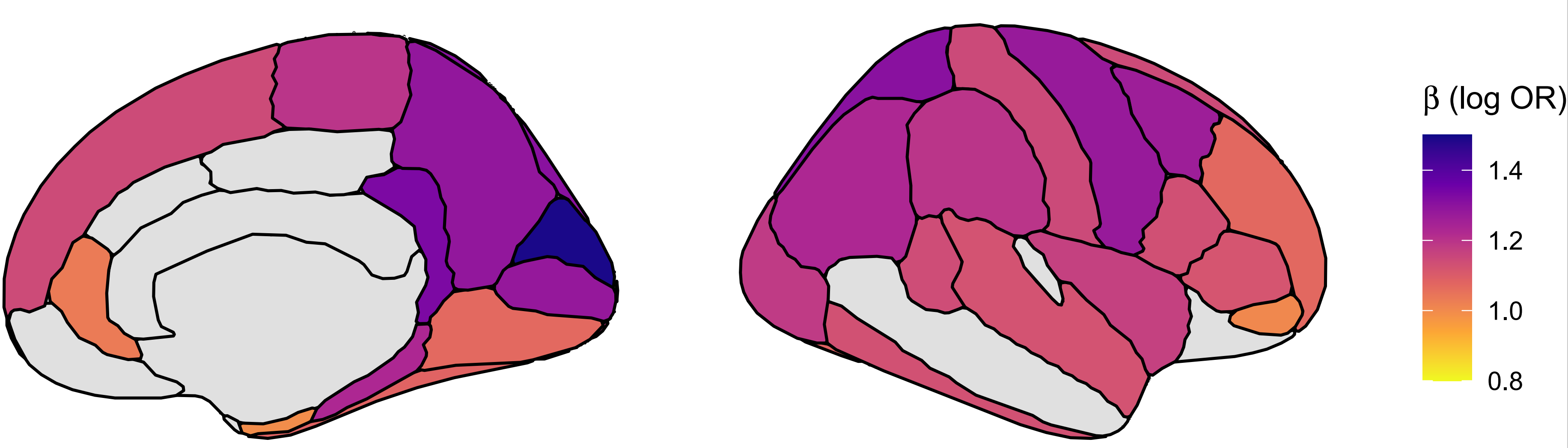

**Supplementary Figure 12. Regional distribution of odds ratio results.** The figure shows the ORs pr region, identical to what is presented in the forest plot in Figure 4 in the main manuscript. The annual percent volume change was averaged across hemispheres, and the results bare displayed on the right hemisphere for convenience. Inferior lateral ventricles are not shown. The figures is created using *ggseg software (21).*

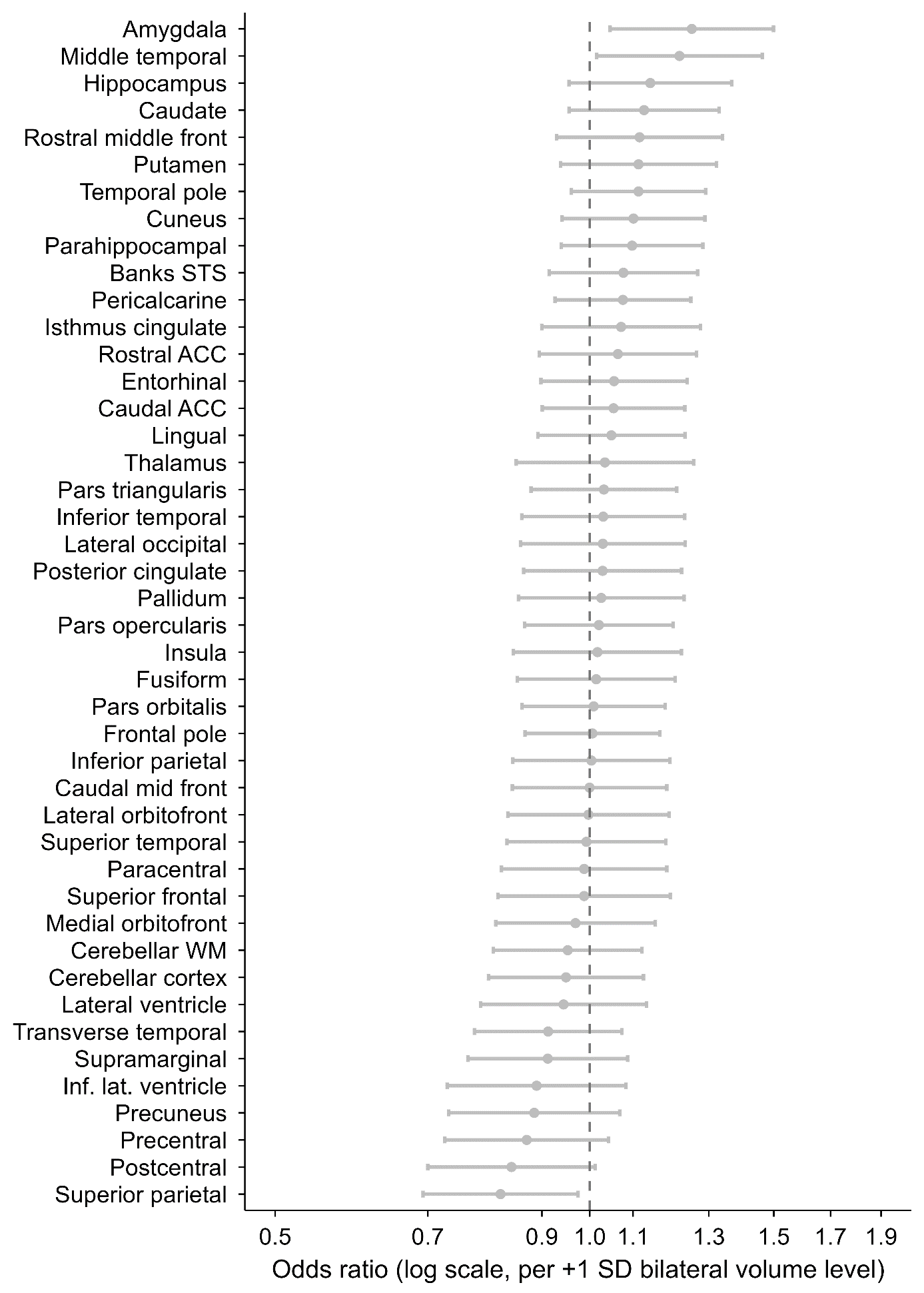

**Supplementary Figure 13. Brain volume levels in stable performers versus non-** **stable performers.** Odds ratios (ORs) with 95% confidence intervals from univariate logistic regression models relating bilateral regional brain volume levels to conservative stable performers status. Each point represents the OR associated with a 1 SD higher bilateral volume in a given region, comparing conservative stable performers to lenient non- stable performers. Analyses were restricted to the same participants as in the main brain-change (slope) analysis. For each region, a robust intercept and median-age level were estimated from Theil–Sen regression, and these median-age levels were z-standardized across participants. Logistic models were then fit separately for each region, with stable performers status (1 = conservative stable performers, 0 = lenient non- stable performers) as the outcome and the regional level (per +1 SD), age at the midpoint of brain follow-up, sex, and estimated total intracranial volume as predictors. Odds ratios > 1 indicate higher odds of being a conservative stable performers with larger regional volume; odds ratios < 1 indicate higher odds of being a non- stable performers with larger volume. The x-axis is on a logarithmic scale from 0.5 to 1.9. No regions survived false discovery rate control at q < .05 (Benjamini–Hochberg) across all 44 bilateral subcortical and cortical regions. This figure complements the main-text slope analysis.

**Section 7: Robustness checks for analyses of what characterizes stable performers**

As a comparability check, we re-ran all univariate models without weights (full results in SI). Effect sizes were essentially identical across approaches (correlation of log-ORs: r = 0.995), and every variable that was significant in the unweighted analysis also remained significant with weighting, while the weighted analysis identified a small number of additional, borderline associations. Thus, weighting improved sensitivity to signals in the minority class without changing the substantive pattern of effects.

To assess whether variable missingness affected conclusions, we re-estimated all age-adjusted univariate models restricting predictors to those with ≤20% missing data. Effect sizes were virtually unchanged (correlation of log-odds ratios with the main analysis: r = 1.000), indicating that missingness did not bias estimates. Further, re-estimating all univariate models after standardizing continuous predictors within sex yielded highly concordant effect sizes (β correlation = 0.985).

The negative association between numeracy (percentage) and being a stable performer, combined with the positive association between numeracy (subtraction) and being a stable performer in the univariate analysis, was puzzling. The subtraction test was introduced in SHARE wave 3 and has limited amount of variation in scores at young ages, with substantial ceiling effects; more than 50% of the sample below age 80 years gets the highest possible score on the test. In a post hoc model including both numeracy measures and age, the percentage numeracy score was negatively related to stable performers status (β = −0.124 ± 0.016; OR per +1 SD ≈ 0.88, p < 4e−14), whereas the subtraction numeracy was positively related (β = 0.075 ± 0.017; OR ≈ 1.08, p ≈ 1.7e−05). The scores were positively correlated (Pearson’s r ≈ .38, Spearman ≈ 0.35), and both correlated negatively with age (Pearson’s r ≈ -.16 and -.17, respectively) and positively with ISCED education (Pearson’s r ≈ .40 and .34, respectively). The effects persisted with education in the model (ORs ≈ 0.87 and ≈ 1.07, respectively). The general pattern replicated across different coding and data extraction schemes, within 10-year age-groups, in sex- and education-stratified analyses, and when running a joint model forcing both numeracy variables in, thereby controlling for each other, the opposite directions still existed (percentage OR = 0.88, subtraction OR = 1.08). Hence, although unexpected, the results suggest construct differences between the subtraction and the percentage tests - they share variance (positive correlation), but the unique parts push odds in opposite directions.

To assess the joint predictive value of the candidate factors, we fit an age-adjusted elastic-net logistic regression with simple median/mode imputation for missingness, yielding modest but above chance discrimination (AUC = 0.614, CI: 0.606-0.622) (Figure 9). Four variables were retained with significant associations in the maximum likelihood estimation logistic regression refit used for inference: stable performers had better orientation performance (OR = 1.08 per +1 SD, 95% CI 1.04–1.12, p = 4.8×10⁻⁵), while non- stable performers reported worse health (OR = 0.96 per +1 SD toward “more problems”, 95% CI 0.93–0.99, p = 8.2×10⁻³), more depressive symptoms (OR = 0.94 per +1 SD, 95% CI 0.91–0.98, p = 1.2×10⁻³) and higher scores on the percentage math test (OR = 0.88 per +1 SD, 95% CI 0.86–0.91, p = 1.1×10⁻^14^). The overall AUC indicates limited out-of-sample discrimination, suggesting that no single domain strongly separates stable performers from non- stable performers when considered jointly. An age-only model achieved close to the same performance (AUC = 0.608, 95% CI: 0.600–0.616). Although the difference between the full model and the age-only model was statistically significant (ΔAUC = 0.003, DeLong p = 0.006), the incremental gain was negligible, indicating that age captures nearly all discriminative signal.

**Section 8: Robustness checks for mortality analyses**

We asked whether mortality differed between stable performers and non- stable performers. Using Cox models with chronological age as the time scale, follow-up began at each person’s classification age (the visit closest to their median age across waves) and ended at death or last observation. The landmark cohort included 48,185 individuals (4,853 deaths), with 5,261 stable performers (419 deaths; 4,842 censored) and 42,924 non-stable performers (4,434 deaths; 38,490 censored). Because age was the analysis time, risks were compared at the same ages over follow-up (age is controlled by design). In the sex-adjusted landmark model, stable performers (conservative rule) had a higher hazard of death, HR = 1.155 (95% CI 1.045–1.278), p = 0.005, while females had lower hazard, HR = 0.616 (0.582–0.652), p < 0.001. Stratifying baseline hazards by country, given heterogeneity in mortality and drop-out across SHARE countries, slightly strengthened the contrast, HR = 1.168 (1.056–1.293), p = 0.003.

To probe whether differential drop-out could bias this contrast by stable performers level, we estimated inverse-probability-of-censoring weights (IPCW) from a visit-to-visit “return alive to next wave” model (logistic regression with interval midpoint of age, sex, country, stable performers label; in a sensitivity model also education [ISCED-97], self-rated health, ADL, IADL, and depressive symptoms). We stabilized by the marginal retention probability and truncated extremes. Person-level weights were near 1 (median 1.009; 95th percentile 1.047; 99th percentile 1.174), and the weighted landmark estimate at the classification anchor was essentially unchanged, HR = 1.167 (1.060–1.285), p = 0.002, indicating little evidence that informative censoring drives the primary label contrast (weights near 1; weighted HR ≈ unweighted). We also isolated the most distinct groups by comparing conservative stable performers to lenient non- stable performers only. In sex-adjusted, country-stratified Cox models, the hazard was higher for stable performers overall, HR = 1.156 (1.042–1.281), p = 0.006. Weighted age-banded analyses suggested this was most apparent from age 70: HRs were 1.156 (0.979–1.366), p = 0.090 for ages 55–70; 1.234 (1.048–1.454), p = 0.014 for 70–80; 1.277 (1.051–1.551), p = 0.009 for 80–90; and 0.496 (0.217–1.134), p = 0.062 for 90–100. Collapsing to ≥70 yielded HR = 1.164 (1.029–1.317), p = 0.013. A weighted stable performers × age interaction did not improve fit (interaction ≈ 0; LR test p ≈ 0.98), consistent with level shifts rather than a smooth interaction.

Finally, we re-anchored follow-up at each person’s last observed age. Under this specification, the label contrast attenuated to the null—country-stratified HR = 1.048 (0.940–1.169), p = 0.40; without strata HR = 1.050 (0.947–1.164), p = 0.35, suggesting that the initial HR>1 is better explained by selection and timing around when the label is assigned rather than by stable performers status itself being causally related to higher mortality.

Because a binary label discards information about degree and trajectory, we then tested memory level and slope as continuous features derived from each person’s robust Theil–Sen line. Level was evaluated at the classification age and at a common age 70. Per 1-SD higher level at the classification age, the sex-adjusted, country-stratified hazard ratio was 0.787 (0.760–0.815), attenuating to 0.850 (0.818–0.883) with adjustment for education, self-rated health, chronic conditions, ADL, and IADL. Evaluated at age 70, associations were similarly strong and robust, with sex-adjusted HR = 0.818 (0.796–0.839) and fully adjusted HR = 0.845 (0.823–0.868). Standardizing level within age bands by z-transforming each person’s memory level relative to peers of similar classification age yielded HR = 0.684 (0.663–0.706). In a joint model including both level at the classification age and slope, both contributed, with a larger effect for level (level HR = 0.787; slope HR = 0.904; both p < 0.001). The IPCW weighted model with continuous features only (level and slope) closely matched these results (level HR = 0.799, 0.773–0.827; slope HR = 0.908, 0.882–0.934), and adding the stable performers label on top of level and slope further improved fit (label HR = 1.512, 1.353–1.691; LR test versus features-only: χ² = 49.26, df = 1, p = 2.24×10⁻¹², AIC reduced from 58,564.9 to 58,517.6), indicating that the label still captures residual information not fully contained in level and slope (Figure 9).

Taken together, the apparent HR > 1 for stable performers in the primary landmark contrast was sensitive to design choices, i.e. how labels are set and where follow-up is anchored, and showed its clearest separation from age 70 onward, but without evidence for a continuous age interaction. In contrast, memory level showed a clear, stable association with mortality, especially on a common age scale or when standardized within age bands, and slope retained a modest independent contribution. Cognition related to survival most cleanly through level, and, to a lesser degree, slope, while the stable performers label was a coarser, timing-sensitive summary that nonetheless added some information beyond the continuous features.

**Section 9: Identification of burst-decline in individual SHARE participants**

Having shown that a burst-decline process can reproduce the group trend, we next asked whether intervals where memory drops quickly and stabilizes at a lower level can be detected in individual SHARE participants. We removed each person’s gradual age trend by subtracting their Theil–Sen line from the practice-adjusted scores and flagged an interval as a burst if it was (i) fast (≤ −0.50 points/year), (ii) substantial (drop ≥ 1.5 points), and (iii) short (≤ 3 years). Recovery was evaluated ≥2 years after onset: if the best future residual regained ≥50% of the drop, the burst was labeled “recovered”.

We found that bursts were common and sizeable. Typical drops were ~3.4 points (IQR ≈2.3) over ~2 years (IQR 2–3), and most did not recover (≈80–94% across recovery thresholds and time-windows). Varying drop size, speed, and duration produced similar or higher rates, indicating that the phenomenon is not sensitive to specific thresholds. Bursts were meaningfully related to stable performers status: stable performers (conservative rule) showed fewer bursts overall and fewer unresolved bursts (any burst 0.51 vs 0.85; any non-recovered 0.39 vs 0.47; RR 0.60 and 0.83). After adjusting for number of intervals exposure, stable performers still had lower odds of a non-recovered burst (OR ≈0.70, 95% CI 0.66–0.74). Using a stricter definition to find “big bursts”, defined as drops ≥2× each person’s residual SD, stable performers remained much less likely to exhibit any non-recovered event (adjusted OR ≈0.16, 95% CI 0.151–0.174).

To test whether such patterns could arise from noise alone, we permuted residuals within person, preserving spacing and variability, and reapplied the burst rules. In the observed data, 54.1% of participants had ≥1 non-recovered big burst (0.55 events per 10 person-years). Under the permutation null, medians were 45.1% and 0.463. Observed rates exceeded permutation medians in all age bands (50–80+, p ≈ 0). Together, these analyses indicate that bursts are real, time-structured phenomena rather than artifacts of noise or regression to the mean. Stable performers have fewer bursts overall, and especially fewer unresolved big bursts, which can help explain how some individuals remain stable for a decade while still performing at relatively low levels.

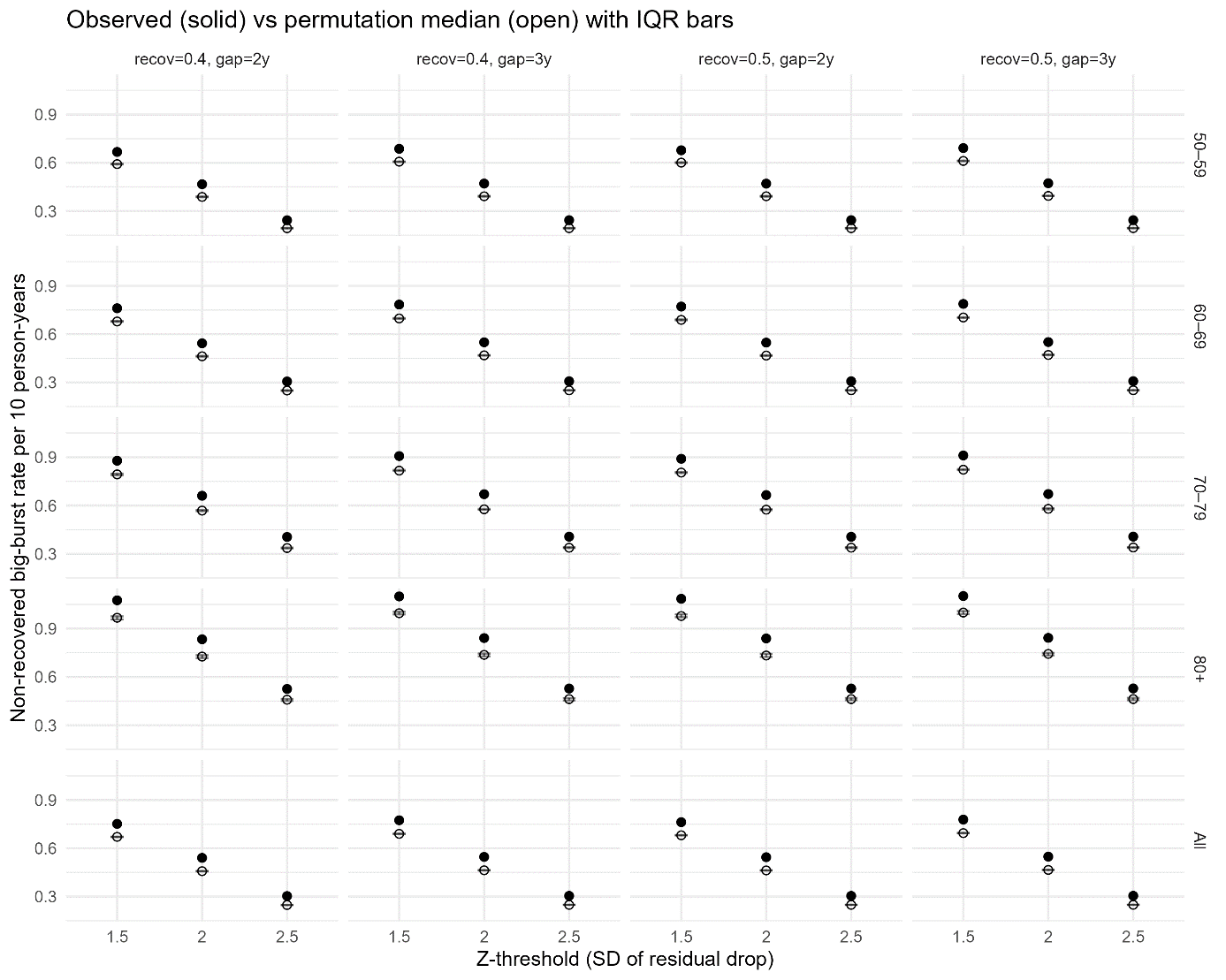

**Supplementary Figure 14. Non-recovered “big burst” events are more frequent than expected under a noise-only timing model.** The figure compares observed rates of non-recovered big drops (“bursts”) in episodic-memory scores with permutation-based null expectations that preserve each person’s variability but randomize the timing of fluctuations. Results are shown across age bands (50–59, 60–69, 70–79, 80+, and All ages) and across analysis settings that vary the drop size threshold and the recovery rule. Points show observed values; bands/bars show the permutation distribution (median and interquartile range, IQR). Right-tailed permutation p-values (as or more extreme than observed) are annotated; all tested configurations yielded p < .001. Solid symbols = observed outcome; hollow symbols/lines or shaded bands = permutation null (median with IQR). Error bars depict IQR.

**Section 10: Offset-slope analyses**

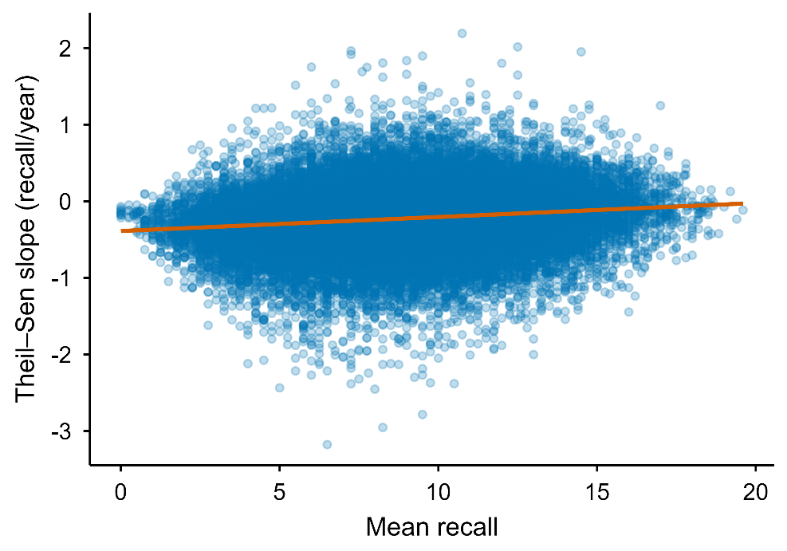

**Supplementary Figure 15. Thiel-Sen slope vs Thiel-Sen estimated recall.**

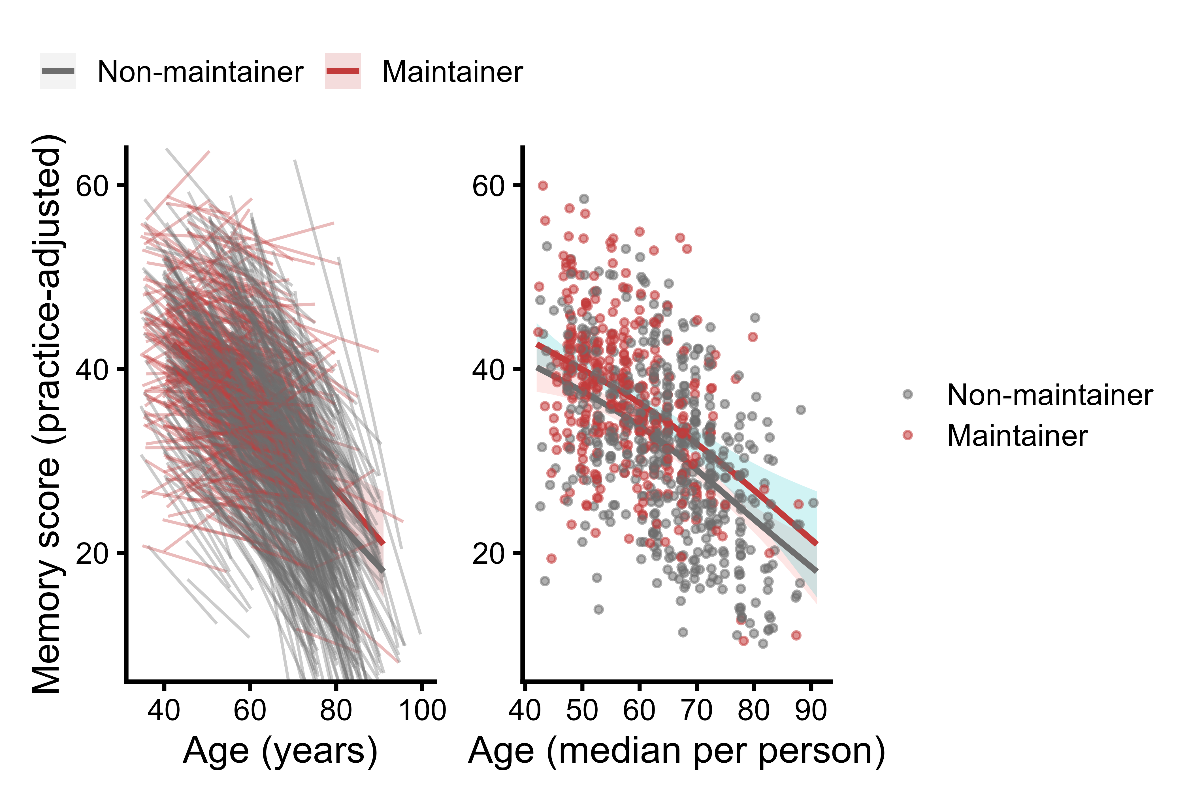

**Supplementary Figure 16. Thiel-Sen-estimated memory in all BETULA participants and stable performers**. Left: Slopes for all stable performers (red) and non- stable performers (grey) in Betula. Right: Memory level for each stable performer (red) and non- stable performer (grey) and the GAM-derived age-trajectories. Shaded area denotes 95% CI.

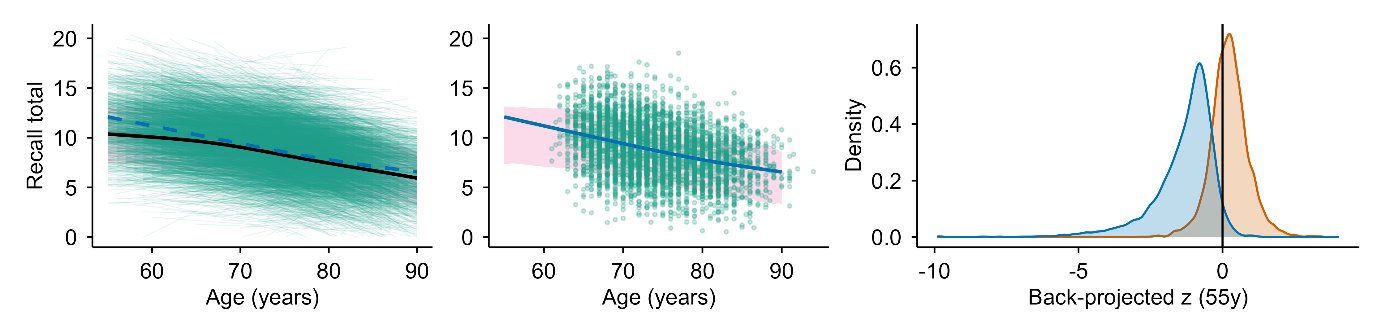

**Supplementary Figure 17. The back-projection paradox in HRS under a transient stable performers (bout) framework**. Left panel: Theil–Sen (TS)–estimated episodic memory level (practice-corrected recall total) as a function of age. Each participant contributes one TS-defined level at their median observed age; the black curve shows the smoothed mean across all eligible HRS participants (≥4 observations), and the dashed curve shows the corresponding mean for participants who exhibited at least one contiguous “stable” bout meeting the conservative criterion (leave-one-out TS lower 95% bound ≥ 0). The shaded band denotes the central 75% of the distribution of TS-estimated levels at each age (moving window), and faint individual lines show the TS-implied trajectories for bout stable performers across their observed age ranges. Middle panel: Same background band as in (A), with points showing TS-estimated levels at the median age for bout stable performers only; the solid curve is the smoothed mean for bout stable performers. Right panel: Distribution of cohort-standardized back-projected performance at age 55 among older participants (defined as having a maximum observed age ≥75). For each participant, a linear TS model was used to back-project to age 55: bout stable performers were projected using the slope and intercept estimated within their identified stable bout (anchored to the bout), whereas non- stable performers were projected using their TS line fit across all observations. Back-projected values were z-standardized within birth cohort using cohort norms at age 55 derived from participants observed in midlife (ages 50–60). The vertical line marks the cohort mean (z = 0). The pronounced left-shift among bout stable performers indicates that extrapolating the stable-period dynamics backward implies implausibly low midlife performance for many individuals, consistent with stability being predominantly a time-limited plateau embedded within a broader decline trajectory rather than a lifelong trait.

**Section 11: Stable bout analyses – stability as a state**

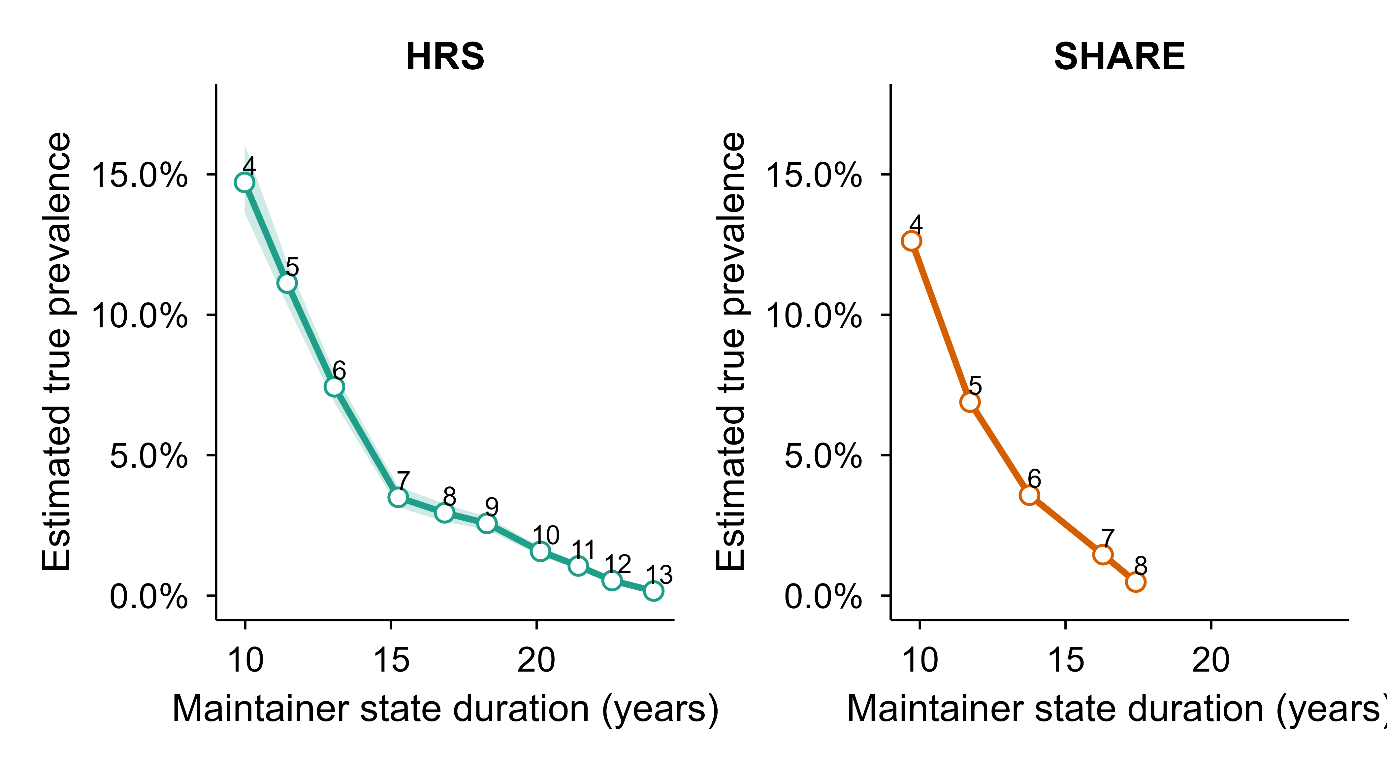

**Supplementary Figure 18. True prevalence of stable periods across the full follow-up (any age) in HRS and SHARE.** Panels show the estimated true prevalence of having at least one contiguous stable period of length ≥W waves (W = 4–13) for the HRS (left) and SHARE (right). The x-axis gives the mean duration (years) of stable periods among participants meeting the corresponding threshold, and the y-axis gives the PPV/Se-corrected true prevalence (Se = 0.80). Stable periods were defined using the conservative criterion of a leave-one-out Theil–Sen 5th-percentile (95% lower bound) ≥ 0 within the candidate window. Shaded ribbons show uncertainty from the simulation-derived null (constructed from the 5th and 95th percentiles of null prevalence and propagated through the correction). Numbers next to points indicate the wave threshold W. Axes are identical across panels to facilitate direct comparison of prevalence and duration between cohorts.

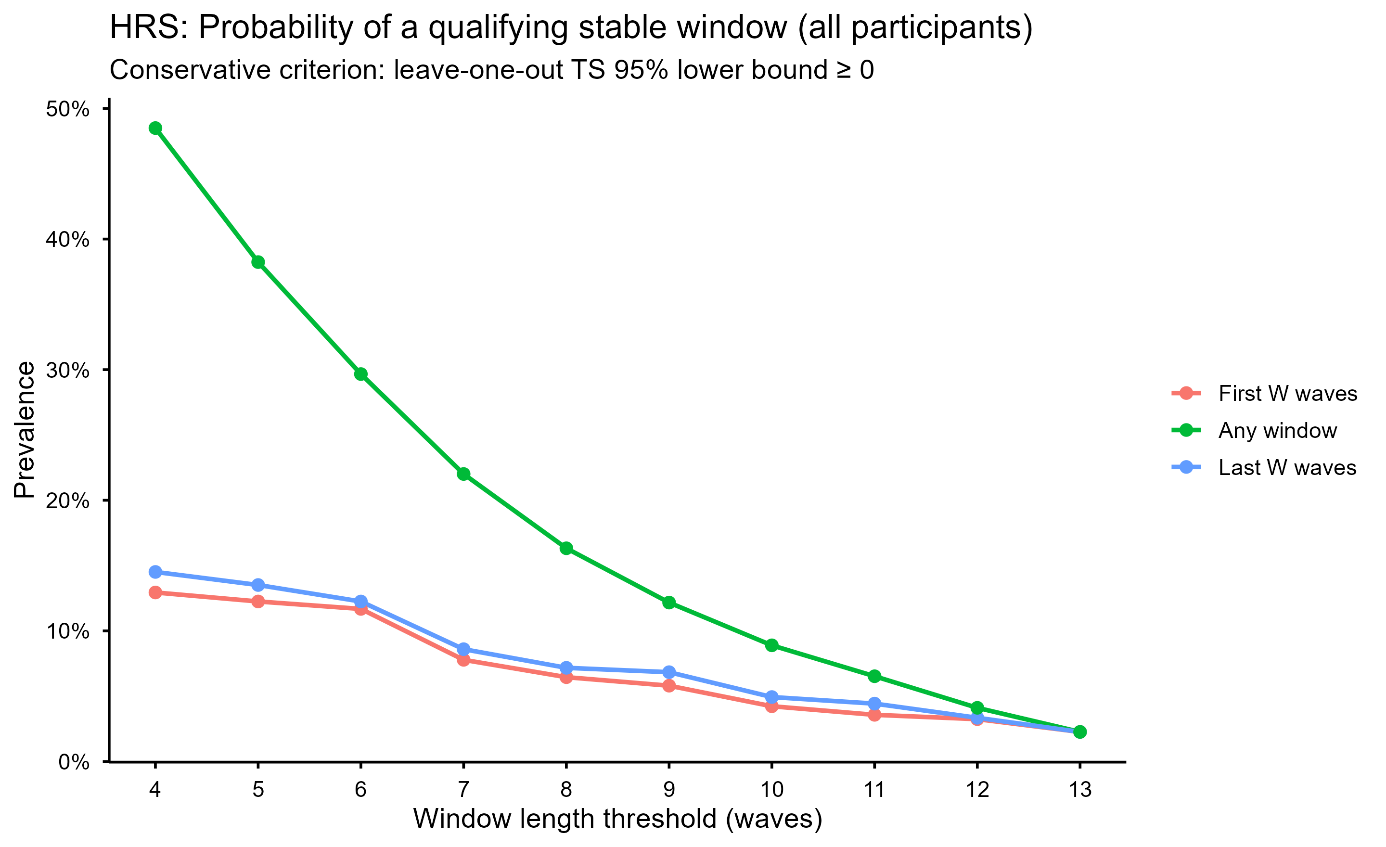

**Supplementary Figure 19. Timing of stable windows within follow-up in HRS supports stability as a time-limited state.** This figure compares the prevalence of qualifying stable windows when stability is required to occur (i) in the first possible contiguous window of length *W* (“First *W* waves”), (ii) in the last possible contiguous window of length *W* (“Last *W* waves”), or (iii) anywhere within the observation series (“Any window”). Analyses were conducted in HRS using practice-corrected episodic memory scores and the conservative stable performers criterion: a contiguous window qualifies as stable if the 5th percentile lower bound of leave-one-out Theil–Sen slopes within the window is non-negative (LOO TS 95% lower bound ≥ 0). For each threshold *W* (4–13 waves), participants were included if they had at least *W* assessments (N_eligible shown in the accompanying table output), and prevalence was computed as the proportion of eligible participants meeting the stability criterion under each timing constraint. The “Any window” definition yields substantially higher prevalence because it allows the stable window to occur at any point in the series (multiple opportunities), whereas the “First” and “Last” definitions are each restricted to a single fixed segment. Importantly, prevalence in the first and last windows is similar across thresholds (e.g., *W*=4: 12.9% vs 14.5%; *W*=10: 4.2% vs 4.9%), indicating that stable periods are not preferentially concentrated at either end of follow-up. This pattern is consistent with decline occurring both before and after stable periods, rather than stability reflecting a baseline trait or an end-stage plateau. Mean duration (years) of qualifying windows is shown in the accompanying summary table (computed as end age minus start age for windows meeting the criterion).

**Section 12: Additional information on MRI subsample and Betula datasets**

| **Sample** | **Link** | **PI and/or Admin Contact** | **IRB** |
| --- | --- | --- | --- |
| ADNI | <https://adni.loni.usc.edu/> (O) | Weiner MW; (PI)  (AC) | Approved by the Institutional Review Boards of all of the participating institutions |
| BETULA | <http://www.ufbi.umu.se/english> (R) | Lars Nyberg; (PI) | Regional Ethical Vetting Board at Umeå University |
| OUS/COGNORM | <https://www.med.uio.no/klinmed/english/research/groups/delirium/index.html> | Leiv Otto Watne (PI); Anders Martin Fjell; (PI) | Norwegian Regional Committees for Medical and Health Research Ethics and the Data Protector Officer at Oslo University Hospital |
| HABS | <https://habs.mgh.harvard.edu> (O) | Reisa Sperling; (PI); (AC) | Partners Healthcare Human Research Committee |
| LCBC | <http://lcbc.uio.no> (R) | Anders M Fjell; / Kristine B. Walhovd; (PI) | Norwegian Regional Committee for Medical and Health Research Ethic; Regional Ethical Committee of South Norway |
| OASIS3 | <https://www.oasis-brains.org/> (O) | Pamela J. LaMontagne; (PI); Daniel Marcus; (PI); <https://www.oasis-brains.org/#contact> (AC) | Institutional Review Board of Washington University School of Medicine |
| preventAD | [https://prevent-alzheimer.net](https://prevent-alzheimer.net/?page_id=42&lang=en); <https://openpreventad.loris.ca/> (O) | Jennifer Tremblay; j (PI); <https://openpreventad.loris.ca/contact/> (AC) | The McGill Institutional Review Board and the Douglas Mental Health University Institute Research Ethics Board |
| WRAP | <https://wrap.wisc.edu/> (O) | (AC); Sterling C. Johnson (PI) | Health Sciences IRB of the University of Wisconsin-Madison |
| Mayo/ MCSA | <https://www.gaaindata.org/partner/MCSA> (O) | [](https://www.google.com/search?q=mcsaadrcdatasharing%40mayo.edu&client=firefox-b-d&hs=b5TU&sca_esv=e373368215e95de4&channel=entpr&sxsrf=ANbL-n4x9dLs7EYYukrU2ozGrkEy63vVqw%3A1768222281876&ei=Se5kaY-gNarYwPAP4euuqQo&ved=2ahUKEwj7mYurhYaSAxXfSFUIHXL1ID4QgK4QegQIBRAB&uact=5&oq=MCSA+study+contact+us+mayo&gs_lp=Egxnd3Mtd2l6LXNlcnAiGk1DU0Egc3R1ZHkgY29udGFjdCB1cyBtYXlvMgUQIRigATIFECEYoAEyBRAhGKABSOwMUJQHWJILcAF4AZABAJgBcaABigOqAQM0LjG4AQPIAQD4AQGYAgagAqIDwgIKEAAYRxjWBBiwA5gDAIgGAZAGApIHAzUuMaAHhw2yBwM0LjG4B50DwgcDMC42yAcKgAgB&sclient=gws-wiz-serp&mstk=AUtExfC4Fxtg3K_M7WKx04yfX0bpl99xkUWFgR0f-PFpP7F8C5qNPWGd-UTXzzSXDvFnNHcozWw4sKCveIm3PbYPuDKBabWfP_HU3H1JRZRd0H2qV8scU0oJIuCLM63HZgywUH5lJSdPJ3BlV8EooOh6WQGGcTwYTDNYC9s30tUapHYwYN0KoCU_vT-7m457YPvlv3M92AFcESk0L7k_T10QqK9G3eoK2jEcCUe8lS2NF08TQM5kmY4_2bIXWiUd68YOAJhL9fKkzxwaV8g2OU5ElZGB&csui=3) (AC); Ronald Petersen; [](https://neurodegenerationresearch.eu/cohort/the-mayo-clinic-study-of-aging/) (PI); Clifford R. Jack, Jr. (PI) | Institutional review boards of the Mayo Clinic and Olmsted Medical Center |
| BLSA | <https://www.blsa.nih.gov/> (O) | (AC); Luigi Ferruci (PI); Susan Resnick; (PI-MRI) | Internal Review Board of the Intramural Research Program of the National Institutes of Health |
| AIBL | <https://aibl.csiro.au/research/> (O) | Christopher Rowe; (PI) | Institutional ethics committees of Austin Health, StVincent’s Health, Hollywood |

**Supplementary Table 2: Links, data owner and IRB approvals for each dataset**. Data availability, contact and principal investigator information, and ethical approval for the different datasets used. PI = Principal Investigator. AC = Administrative contact. IRB = Institutional Review Boards. O = Openly available. Automatic or semi-automatic data agreements. R = Restricted. Ad-hoc permission is required. Contact PI or AC for specific details on access to data.

Dataset descriptions

ADNI: The Alzheimer’s Disease Neuroimaging Initiative is a multi-site project led by Doctor Michael W. Weiner to assess the progression of mild cognitive impairment (MCI) and early Alzheimer’s Disease (AD). The present study includes participants from ADNI 1, ADNIGO, ADNI2, and ADNI 3, who were cognitively healthy at baseline (*DX_bl* variable). Only observations in which participants were still cognitively healthy were included as determined by the ADNI team (*DX* variable). Participants were required to have no evidence of ischemic stroke (Hachinski Ischemic Score ≤ 4), a Geriatric Depression scale score < 6, stable medications for 4 weeks before the screening, good auditory and visual acuity, good general health, no medical contraindications to MRI and at least 6 grades of education/work history. General inclusion and exclusion criteria are described elsewhere ^1^. All participants signed an informed consent form and the protocols were approved by the corresponding regional ethical committees in the US and Canada. Data was retrieved in April 2021.

PreventAD: The Pre-symptomatic Evaluation of Experimental or Novel Treatments for AD is a retrospective, long-term study that follows cognitively healthy older individuals with a familiar history of AD. It includes participants enrolled either from an observational cohort or the clinical trial of PREVENT-AD. This study comprises MRI images, blood and CSF samples, and clinical and neuropsychological assessments. Participants had to be at least 60 years old with ≥6 years of education and be cognitively unimpaired at baseline. The Montreal Cognitive Assessment (MoCA) (≥ 26/30) and CDR (= 0) scales were used to assess cognitive abilities. Other exclusion criteria at baseline included medical conditions that prevented longitudinal participation or medical contraindications to MRI, use of acetylcholinesterase inhibitors, other approved prescription cognitive enhancers, hypertension, or substance abuse. The protocols, consent forms, and study procedures were approved by the McGill Institutional Review Board and the Douglas Mental Health University Institute Research Ethics Board. Observations with RBANS > 1SD below the mean and probable MCI, as evaluated by a clinician, were excluded.

HABS: The Harvard Aging Brain Study is an ongoing, long-term observational study that aims to enhance our understanding of brain aging and the early stages of Alzheimer's disease. The study collects PET, MRI data, neuropsychological and clinical assessments. The age range was 50-90 years at the time of baseline assessment and all patients were considered non-clinically impaired at the start of the study. Further, participants had a CDR score of 0, MMSE score ≥ 25, < 11 on the Geriatric Depression Scale, and scored above age- and education-adjusted cutoffs on the 30-Minute Delayed Recall of the Logical Memory Story A, to be included in the study. Participants with a history of alcoholism, drug abuse, head trauma, or current serious medical/psychiatric illness were excluded. Observations with MCI or AD diagnostic (DX variable) were excluded. All participants signed informed consent and the protocol was approved by the Partners Healthcare Human Research Committee. HABS data release 2.20, retrieved in August 2022 via habs.mgh.harvard.edu.

LCBC: The Center for Lifespan Changes in Brain and Cognition cohort (Oslo, Norway) consists of cognitively healthy, community-dwelling participants across the lifespan and is drawn from studies coordinated by the LCBC Research Center (LCBC www.lcbc.uio.no), approved by a Norwegian Regional Committee for Medical and Health Research Ethics. Written informed consent was obtained from all participants. The samples were recruited by a variety of methods such as newspapers and webpage ads. Most participants were recruited for observational studies, while a minority were recruited to cognitive training. All participants had to undergo a standardized health interview before being included in the study, and those with a history of neurological or psychiatric conditions or who reported concerns about their cognitive function were excluded. Additionally, all participants over the age of 40 years were required to score at least 25 on the Mini-Mental State Examination, and observations paired with MMSE ≤ 25 were excluded. Data was retrieved in November 2022.

OASIS3: The Open Access Series of Imaging Studies is a retrospective collection of multimodal data that focuses on aging and AD and is openly accessible to the scientific community. OASIS-3 includes neuroimaging, clinical and neuropsychological data. Participants were recruited through the Washington University Knight Alzheimer Disease Research Center via flyers, word of mouth, and community engagements, and were between 42 and 95 years of age. Only participants deemed cognitively normal at baseline were included. Exclusion criteria included medical conditions that precluded longitudinal participation or medical contraindications for the different study arms. All participants consented to Knight ADRC-related projects following procedures approved by the Institutional Review Board of Washington University School of Medicine. Observations were included until the last observation in which a subject was deemed cognitively healthy as determined by the Clinical Dementia Rating Scale (CDR) ^3^.

AIBL: The Australian Imaging, Biomarker & Lifestyle Flagship Study of Ageing (AIBL) (Ellis et al., 2009) is a prospective study including cognitively normal participants, and patients with mild cognitive impairment, or AD aged 60 years or older. The study assesses the biomarkers, genetic factors, cognitive characteristics, and health and lifestyle factors that are associated with the development of AD, combining techniques such as MRI, positron emission tomography (PET), blood tests, and fluid sample analysis, as well as neuropsychological and clinical assessments. Healthy participants must meet specific criteria, including being free of cognitive impairments and having test performance within 1.5 standard deviation *(SD)* of age-adjusted norms. The test battery and sample description, as well as in-detail general inclusion criteria, have been described in detail previously (Ellis et al., 2009). All individuals from cognitively normal individuals at baseline were included, as defined by the AIBL team (DX variable). All participants signed an informed consent form and the protocol was approved by the institutional human research committees of Austin Health, St Vincent’s Health, Hollywood Private Hospital, and Edith Cowan University (Australia). Data was retrieved in February 2023. See [www.aibl.csiro.au](http://www.aibl.csiro.au/) for further details.

BLSA: The National Institute on Aging Baltimore Longitudinal Study of Aging (BLSA) is Americas longest-running scientific study of human aging. It began in 1958, and neuroimaging was introduced in 1994. The Open dataset includes defaced brain MRIs conducted 2009–2020 for a subset of the BLSA participants scanned in a single 3.T dataset (Ferrucci, 2008). From this open subsample, all participants cognitively healthy at their first observation were included in the analysis. All participants signed an informed consent form, and the protocol was approved by the Internal Review Board of the Intramural Research Program of the National Institutes of Health. Data was retrieved in August 2025.

Mayo: The Mayo Clinic and Olmsted Medical Center Study (MCSA) is a longitudinal, population-based study of residents of Olmsted County, Minnesota. The long-term goals of the Mayo Clinic Study of Aging are to develop tools to predict and prevent cognitive decline and dementia, develop risk-prediction models for cognitive impairment, and conduct aging-related research to promote successful aging. MCSA uses comprehensive medical record linkage to follow residents longitudinally. The study provides detailed clinical, cognitive, and demographic data and has been widely used to examine aging processes, cognitive decline, and neurological disease within a well-defined community cohort. We used an openly released subset (Schwarz et al., 2025). See recruitment details in (Roberts et al., 2008) which included a stratified random selection of subjects from the sampling frame, a letter of invitation followed by a recruitment telephone call, and finally an in-person evaluation or a telephone assessment. For the present study only participants age > 50 and cognitively healthy at baseline were selected. The project was approved by the institutional review boards of the Mayo Clinic and Olmsted Medical Center. All persons examined as part of the study were informed of the scope of the project and signed an informed consent form including a Health Insurance Portability and Accountability Act (HIPAA) authorization. The data was retrieved in June 2025.

WRAP: The Wisconsin Registry for Alzheimer’s Prevention (WRAP) is a longitudinal cohort study enriched for Alzheimer’s disease risk, combining detailed cognitive assessments with multimodal neuroimaging. The study follows initially late–middle-aged adults to characterize early brain changes associated with Alzheimer’s disease risk, progression, and preclinical pathology. See (Sager et al., 2005) for detailed enrollment criteria. Briefly, individuals required to be between 40 and 65 years, speak English, and have a parent with either autopsy-confirmed or probable AD as defined by NINCDS-ADRDA research criteria. That is, WRAP volunteers were recruited from adult children of persons with AD evaluated in the Memory Assessment Clinic at the University of Wisconsin–Madison or an affiliated memory clinic. Enrollment into WRAP began in November 2001 and is currently ongoing. The assessments include information on cognition, questionnaires, lab tests, genotyping, and MRI and PET imaging. For this study, we used cognitively healthy individuals aged 50 or more at baseline. All study procedures have been approved by the Health Sciences IRB of the University of Wisconsin-Madison and the participants gave informed consent. Data was retrieved in July 2025.

**Section 13: Funding**

LCBC is supported by the European Research Council under grant agreements no. 283634 and no. 725025 (to A.M.F.) and no. 313440 (to K.B.W.), as well as the Norwegian Research Council (325878, 262453 to A.M.F.; 325001, 301395, 239889 to K.B.W.; 249931 to A.M.F & K.B.W.; 324882 to DVP; 325415 to HG), the National Association for Public Health’s dementia research program, Norway (to A.M.F.), and the University of Oslo through the UiO:Life Science convergence environment (to A.M.F). Betula is supported by a scholar grant from the Knut and Alice Wallenberg foundation to L.N. Additional funding include:

The SHARE data collection has been funded by the European Commission, DG RTD through FP5 (QLK6-CT-2001-00360), FP6 (SHARE-I3: RII-CT-2006-062193, COMPARE: CIT5-CT-2005-028857, SHARELIFE: CIT4-CT-2006-028812), FP7 (SHARE-PREP: GA N°211909, SHARE-LEAP: GA N°227822, SHARE M4: GA N°261982, DASISH: GA N°283646) and Horizon 2020 (SHARE-DEV3: GA N°676536, SHARE-COHESION: GA N°870628, SERISS: GA N°654221, SSHOC: GA N°823782, SHARE-COVID19: GA N°101015924) and by DG Employment, Social Affairs & Inclusion through VS 2015/0195, VS 2016/0135, VS 2018/0285, VS 2019/0332, VS 2020/0313, SHARE-EUCOV: GA N°101052589 and EUCOVII: GA N°101102412. Additional funding from the German Federal Ministry of Education and Research (01UW1301, 01UW1801, 01UW2202), the Max Planck Society for the Advancement of Science, the U.S. National Institute on Aging (U01_AG09740-13S2, P01_AG005842, P01_AG08291, P30_AG12815, R21_AG025169, Y1-AG-4553-01, IAG_BSR06-11, OGHA_04-064, BSR12-04, R01_AG052527-02, R01_AG056329-02, R01_AG063944, HHSN271201300071C, RAG052527A) and from various national funding sources is gratefully acknowledged (see [www.share-eric.eu](http://www.share-eric.eu)).

Data from the Health and Retirement Study (HRS) were obtained from the University of Michigan. HRS is sponsored by the U.S. National Institute on Aging (grant U01AG009740) and conducted by the University of Michigan, with additional support from the U.S. Social Security Administration. RAND HRS Longitudinal File (1992–2022 release, Stata format) was produced by the RAND Center for the Study of Aging, with funding from the National Institute on Aging (grant numbers NIA U01AG009740 and NIA R01AG073289) and the Social Security Administration. Santa Monica, CA.

ADNI: The ADNI was launched in 2003 as a public-private partnership, led by Principal Investigator Michael W. Weiner, MD. The primary goal of ADNI has been to test whether serial magnetic resonance imaging (MRI), positron emission tomography (PET), other biological markers, and clinical and neuropsychological assessment can be combined to measure the progression of mild cognitive impairment (MCI) and early Alzheimer’s disease (AD). For up-to-date information, see <https://adni.loni.usc.edu/>. As such, the investigators within the ADNI contributed to the design and implementation of ADNI and/or provided data but did not participate in analysis or writing of this report. A complete listing of ADNI investigators can be found at: <http://adni.loni.usc.edu/wp-content/uploads/how_to_apply/ADNI_Acknowledgement_List.pdf>. Data collection and sharing for this project were funded by the ADNI (NIH Grant U01 AG024904). ADNI is funded by the National Institute on Aging, the National Institute of Biomedical Imaging and Bioengineering, and through generous contributions from the following: AbbVie, Alzheimer's Association; Alzheimer's Drug Discovery Foundation; Araclon Biotech; BioClinica, Inc.; Biogen; Bristol-Myers Squibb Company; CereSpir, Inc.; Cogstate Eisai Inc.; Elan Pharmaceuticals, Inc.; Eli Lilly and Company; EuroImmun; F. Hoffmann-La Roche Ltd and its affiliated company Genentech, Inc.; Fujirebio; GE Healthcare; IXICO Ltd.; Janssen Alzheimer Immunotherapy Research & Development, LLC.; Johnson & Johnson Pharmaceutical Research & Development LLC.; Lumosity; Lundbeck; Merck & Co., Inc.; Meso Scale Diagnostics, LLC.; NeuroRx Research; Neurotrack Technologies; Novartis Pharmaceuticals Corporation; Pfizer Inc.; Piramal Imaging; Servier; Takeda Pharmaceutical Company; and Transition Therapeutics. The Canadian Institutes of Health Research is providing funds to support ADNI clinical sites in Canada. Private sector contributions are facilitated by the Foundation for the National Institutes of Health (http://www.fnih.org). The grantee organization is the Northern California Institute for Research and Education, and the study is coordinated by the Alzheimer's Therapeutic Research Institute at the University of Southern California. ADNI data are disseminated by the Laboratory for Neuro Imaging at the University of Southern California.

OUS-COGNORM is funded by the South-Eastern Norway Regional Health Authorities (#2017095) The Norwegian Health Association (#19536) and by Wellcome Leap’s Dynamic Resilience Program (jointly funded by Temasek Trust) #104617).

AIBL: Data used in the preparation of this article was obtained from the Australian Imaging Biomarkers and Lifestyle flagship study of ageing (AIBL) funded by the Commonwealth Scientific and Industrial Research Organisation (CSIRO) which was made available at the ADNI database (www.loni.usc.edu/ADNI). The AIBL researchers contributed data but did not participate in analysis or writing of this report. AIBL researchers are listed at [www.aibl.csiro.au](http://www.aibl.csiro.au).

BLSA: This study uses research data from The Baltimore Longitudinal Study of Aging study (BLSA) that has been made available through the LONI IDA and GAAIN. The BLSA is supported by the Intramural Research Program (IRP) of the National Institute on Aging (NIA). The authors thank the investigators, staff, and participants of the BLSA study and the neuroimaging sub-study for making the data available. BLSA investigators have not contributed to nor approved, and are not in any way responsible for, the contents of this article.

HABS_HD: Research reported on this publication was supported by the National Institute on Aging of the National Institutes of Health under Award Numbers R01AG054073, R01AG058533, R01AG070862, P41EB015922 and U19AG078109. The content is solely the responsibility of the authors and does not necessarily represent the official views of the National Institutes of Health.

PREVENT-AD: PREVENT-AD was funded by the Canadian Institutes of Health Research, McGill University, the Fonds de Recherche du Québec – Santé, Alzheimer’s Association, Brain Canada, the Government of Canada, the Canada Fund for Innovation, the Douglas Hospital Research Centre and Foundation, the Levesque Foundation, an unrestricted research grant from Pfizer Canada. Private sector contributions are facilitated by the Development Office of the McGill University Faculty of Medicine and by the Douglas Hospital Research Centre Foundation (<http://www.douglas.qc.ca/>).

OASIS3: Longitudinal Multimodal Neuroimaging: Principal Investigators: T. Benzinger, D. Marcus, J. Morris; NIH P30 AG066444, P50 AG00561, P30 NS09857781, P01 AG026276, P01 AG003991, R01 AG043434, UL1 TR000448, R01 EB009352. AV-45 doses were provided by Avid Radiopharmaceuticals, a wholly owned subsidiary of Eli Lilly.

MCSA: Data used in preparation of this article were shared by the Mayo Clinic Study of Aging (MCSA). The MCSA is funded by the following sources: NIH U01 AG006786, R01 AG034676, R37 AG011378, R01 AG041851, R01 NS097495, R01 AG056366, R01 AG068206, P30 AG062677, GHR Foundation, Elsie and Marvin Dekelboum Family Foundation, Liston Award, Schuler Foundation, Alexander Foundation, Mayo Foundation for Medical Education and Research.

WRAP: Data used in preparation of this articles was made kindly available by the University of Wisconsin. The Wisconsin Registry for Alzheimer's Prevention (WRAP) was funded by the following NIH sources: R01AG027161, and R01AG021155, R01AG062285, and R01AG062167 for MRI data collection.

VETSA: VETSA is supported by National Institute of Health grants NIA R01 AG018384 (PI: Lyons), R01 AG018386 (PI: Kremen), R01 AG022381 (PI: Kremen), and R01 AG022982 (PI: Kremen), and, in part, with resources of the VA San Diego Center of Excellence for Stress and Mental Health. The Cooperative Studies Program of the Office of Research & Development of the United States Department of Veterans Affairs has provided financial support for the development and maintenance of the Vietnam Era Twin (VET) Registry. The content of this manuscript is solely the responsibility of the authors and does not necessarily represent the official views of the NIA/NIH, the Veterans’ Administration, or the VETSA investigators

**Section 14: Additional SHARE and Betula analyses**

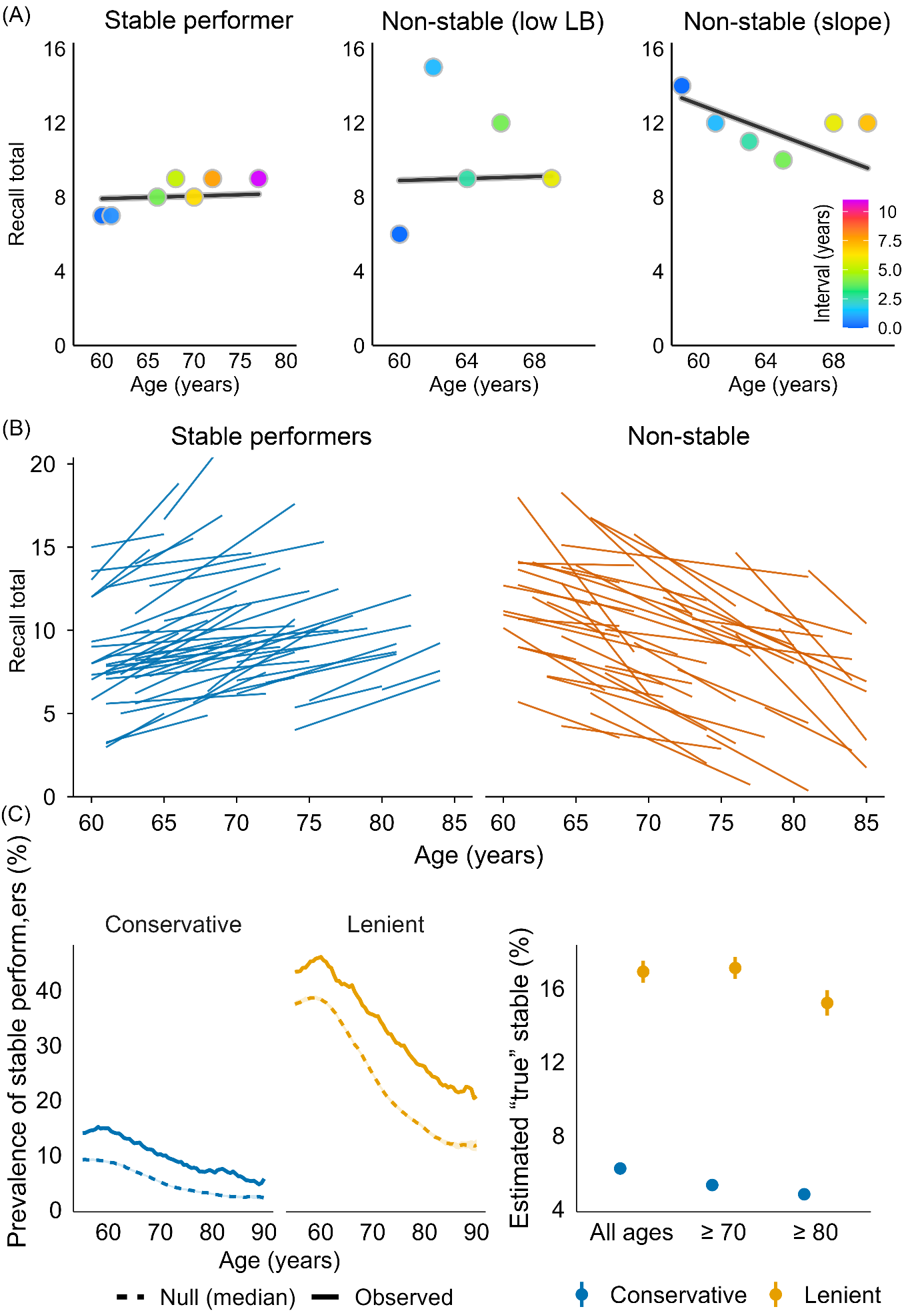

**Supplementary Figure 20. Prevalence of stable performers in SHARE**. Left - Observed performance prevalence (solid line) vs. null expectation (dashed line) if all participants follow the group-average age-expected decline and given the measurement noise in the data (median, shaded band = null IQR), separately for the conservative and the lenient stability criteria. Right - Points show the estimated proportion of true stable performers for the conservative (blue) and lenient (orange) rules, evaluated across all ages, ≥70, and ≥80 years. Vertical bars indicate uncertainty bounds derived from the null simulation.

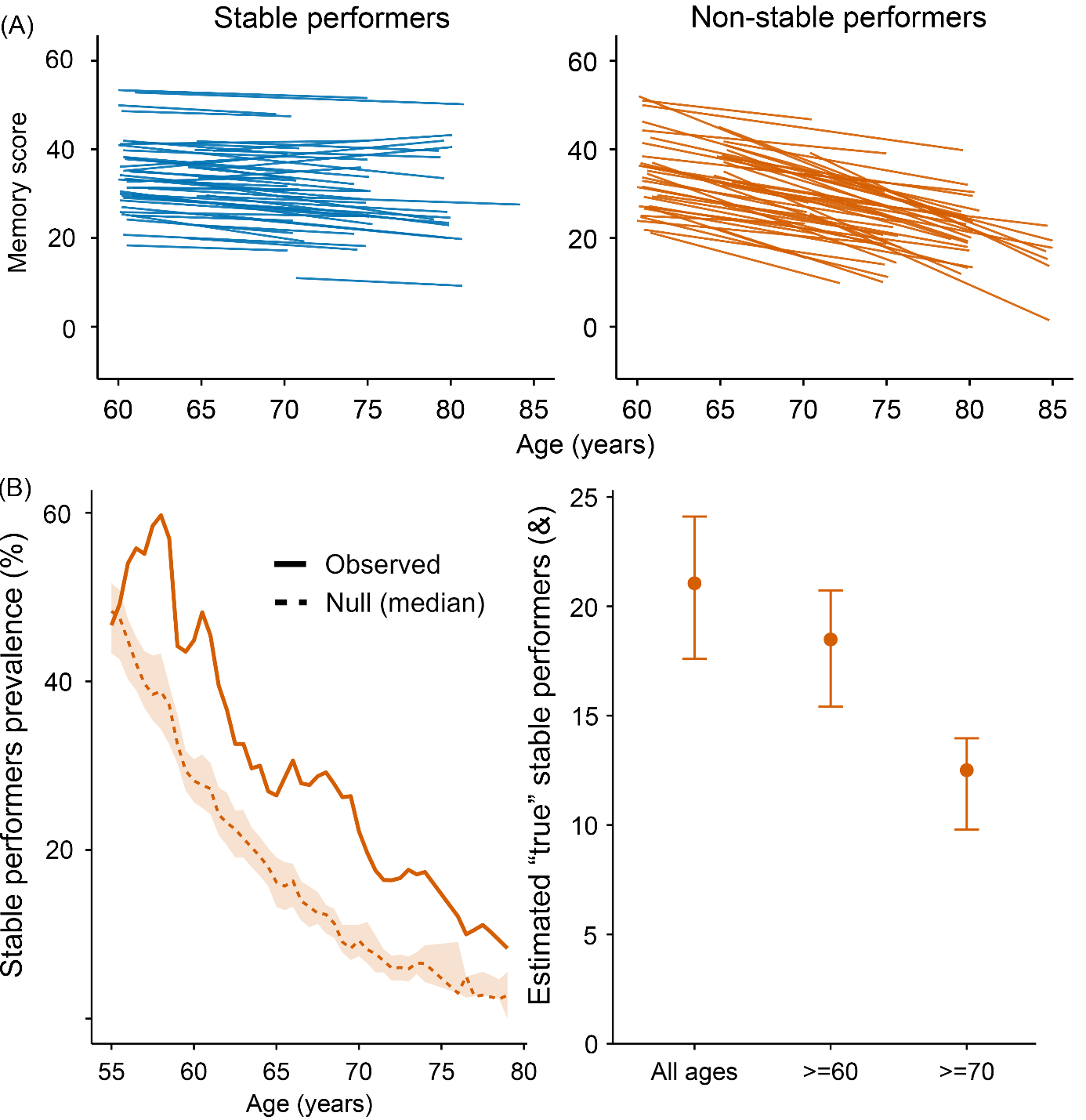

**Supplementary Figure 21. Prevalence of stable performers in Betula.** A: Lines show Theil–Sen slopes for 50 randomly selected stable performers (blue) and non-stable performers (orange). B: Left - Observed prevalence of stable performers (solid line) vs. null expectation (dashed line) (median, shaded band = null IQR). Right - Points show the PPV-adjusted estimated proportion of true stable performers (lenient rule: 𝛿 ≥ -0.4, LB50%) assuming a sensitivity of 0.80. Vertical bars indicate uncertainty bounds derived from the null simulation (optimistic/pessimistic envelopes).

**Section 15: Consortia authors**

Data used in preparation of this article were obtained from the Alzheimer’s Disease Neuroimaging Initiative (ADNI) database (adni.loni.usc.edu). As such, the investigators within the ADNI contributed to the design and implementation of ADNI and/or provided data but did not participate in analysis or writing of this report. A complete listing of ADNI investigators can be found at: <http://adni.loni.usc.edu/wp-content/uploads/how_to_apply/ADNI_Acknowledgement_List.pdf>

The Health and Aging Brain Study (HABS-HD) Study Team:

HABS-HD MPIs: Sid E O'Bryant, Kristine Yaffe, Arthur Toga, Robert Rissman, & Leigh Johnson; and the HABS-HD Investigators: Meredith Braskie, Kevin King, James R Hall, Melissa Petersen, Raymond Palmer, Robert Barber, Yonggang Shi, Fan Zhang, Rajesh Nandy, Roderick McColl, David Mason, Bradley Christian, Nicole Phillips, Stephanie Large, Joe Lee, Badri Vardarajan, Monica Rivera Mindt, Amrita Cheema, Lisa Barnes, Mark Mapstone, Annie Cohen, Amy Kind, Ozioma Okonkwo, Raul Vintimilla, Zhengyang Zhou, Michael Donohue, Rema Raman, Matthew Borzage, Michelle Mielke, Beau Ances, Ganesh Babulal, Jorge Llibre-Guerra, Carl Hill and Rocky Vig.
